# Probabilities of HIV-1 bNAb development in healthy and chronically infected individuals

**DOI:** 10.1101/2022.07.11.499584

**Authors:** Christoph Kreer, Cosimo Lupo, Meryem S. Ercanoglu, Lutz Gieselmann, Natanael Spisak, Jan Grossbach, Maike Schlotz, Philipp Schommers, Henning Gruell, Leona Dold, Andreas Beyer, Armita Nourmohammad, Thierry Mora, Aleksandra M. Walczak, Florian Klein

**Affiliations:** Laboratory of Experimental Immunology, Institute of Virology, Faculty of Medicine and University Hospital Cologne, University of Cologne, 50931 Cologne, Germany; Laboratoire de physique de l’Ecole normale supérieure, CNRS, PSL University, Sorbonne Université, and Université Paris Cité, 75005 Paris, France; German Center for Infection Research, Partner Site Bonn-Cologne, 50931 Cologne, Germany; Excellence Cluster on Cellular Stress Responses in Aging Associated Diseases, University of Cologne, 50931 Cologne, Germany; Department I of Internal Medicine, Faculty of Medicine and University Hospital Cologne, University of Cologne, 50937 Cologne, Germany; Department of Internal Medicine I, University Hospital of Bonn, Bonn, Germany; German Center for Infection Research (DZIF), Partner Site Bonn-Cologne, Bonn, Germany; Center for Molecular Medicine Cologne (CMMC), University of Cologne, 50931 Cologne, Germany; Max Planck Institute for Dynamics and Self-Organization, Am Faßberg 17, 37077 Göttingen, Germany; Department of Physics, University of Washington, 3910 15th Ave Northeast, Seattle, WA 98195, USA; Fred Hutchinson Cancer Research Center, 1100 Fairview Ave N, Seattle, WA 98109, USA

**Author notes:** These authors contributed equally (Christoph Kreer, Cosimo Lupo, Meryem S. Ercanoglu). Shared senior authorship (Thierry Mora, Aleksandra M. Walczak, Florian Klein). Istituto Nazionale di Fisica Nucleare (INFN), Sezione di Roma I, 00185 Rome, Italy. Corresponding author: Florian Klein. **Author Contributions:** Conceptualization, C.K., T.M., A.M.W., and F.K.; methodology, C.K., C.L., M.S.E., N.S., J.G., A.B., L.D., A.N., T.M., A.M.W, and F.K.; investigation, C.K., L.G., M.S.E.; resources, L.G., M.S., L.D., P.S., and H.G.; formal analysis, C.K., C.L., N.S., and J.G.; writing-original draft, C.K., T.M., A.M.W and F.K.; writing-reviewing and editing: all authors; visualization, C.K.; supervision, A.B., T.M., A.M.W., F.K.; funding acquisition, A.N., T.M., A.M.W., C.K., and F. K.

**Keywords:** HIV-1, broadly neutralizing antibody, bNAb, B cell receptor repertoire, probability

## Abstract

HIV-1 broadly neutralizing antibodies (bNAbs) are able to suppress viremia and prevent infection. Their induction by vaccination is therefore a major goal. However, in contrast to antibodies that neutralize other pathogens, HIV-1-specific bNAbs frequently carry uncommon molecular characteristics that might prevent their induction. Here, we performed unbiased sequence analyses of B cell receptor repertoires from 57 healthy and 46 chronically HIV-1- or HCV-infected individuals and learned probabilistic models to predict the likelihood of bNAb development. We formally show that lower probabilities for bNAbs are predictive of higher HIV-1 neutralization activity. Moreover, ranking of bNAbs by their probabilities allowed to identify highly potent antibodies with superior generation probabilities as preferential targets for vaccination approaches. Importantly, we found equal bNAb probabilities across infected and healthy donors. This implies that chronic infection is not a prerequisite for the generation of bNAbs, fostering the hope that HIV-1 vaccines can induce bNAb development in healthy individuals.

**Significance Statement:** While HIV-1 broadly neutralizing antibodies (bNAbs) can develop in chronically HIV-1-infected individuals, they could not yet be elicited by active vaccination. Here, we computationally demonstrate that HIV-1 bNAbs carry distinct sequence features making them unlikely outcomes of the antibody evolution. However, our approach allowed us to identify bNAbs with higher probabilities of being generated. These candidates can now serve as the most promising targets to be induced by vaccination. Moreover, we show that chronic infection has no influence on the probabilities of finding typical bNAb sequence features in the memory B cell compartment. Both findings are critical to design effective vaccination strategies.

## Introduction

The adaptive immune system is able to cope with a plethora of different antigenic structures by employing a diverse repertoire of lymphocytes, which express unique immune receptors as a consequence of V(D)J recombination during lymphopoiesis (1). B cell receptors (BCRs) further diversify during affinity maturation by somatic hypermutation (SHM) (2, 3), leading to the generation of antibodies with high affinities that can target and neutralize infectious pathogens.

The human immunodeficiency virus-1 (HIV-1), however, is able to outpace the adaptive immune system by quickly evolving into antigenically diverse quasispecies due to its error-prone replication machinery (4). These quasispecies contain viral variants that can escape from autologous circulating antibodies. As a consequence, the immune system adapts to the emerged variants, resembling an ongoing immunological arms race (5). Notably, there is a rare fraction of HIV-1-infected individuals who develop a broad serum neutralization response against numerous viral variants, and from whom monoclonal broadly neutralizing antibodies (bNAbs) have been isolated (6–9; reviewed in 10–13). These antibodies target various sites on the homotrimeric envelope glycoprotein (Env), including the CD4 binding site (CD4bs), the variable loop 1 and 2 apex region (V1/V2 apex), the V3 loop base with its surrounding glycans (V3 loop), the gp120/gp41 interface region with the fusion peptide, the membrane-proximal external region (MPER) of gp41, and the so-called ‘silent-face’ (14). Importantly, bNAbs are able to prevent and treat HIV-1 as well as chimeric simian-(S)HIV-1 infections in animal models (15–20), and have been demonstrated to suppress viremia in HIV-1-infected individuals without notable adverse events or side effects (21–30). For instance, a combination of the CD4bs antibody 3BNC117 and the V3 loop antibody 10-1074 was able to control viremia in 76% of study participants for at least 20 weeks in the absence of antiretroviral therapy (ART) (27). Moreover, preventive treatment with the CD4bs bNAb VRC01 significantly reduced infections with VRC01-sensitive HIV-1 (31), demonstrating that bNAbs are in principle able to prevent infections in humans. Importantly, technical advances and efforts to screen thousands of HIV-1-infected individuals have promoted the isolation of nearly pan-neutralizing bNAbs (9, 32, 33). This next generation of bNAbs, engineered bi- or trispecific antibodies (34–37), and combination therapies with complementary bNAbs (25, 26, 29) have the potential to constrict viral escape and thus improve HIV-1 treatment and prevention strategies.

Despite the progress in passive administration, all efforts to induce highly potent bNAbs through vaccination have failed so far. bNAbs tend to accumulate unusual sequence properties, including high numbers of somatic mutations (6–9, 32, 33, 38), insertions/deletions (6, 8, 9, 38, 39), distinct V_H_ gene segment use (40–42), or exceptionally long complementarity determining regions (CDRs) (7). Long CDRH3 regions as well as the usage of VH4-34 have previously been associated with self-reactive antibodies in autoimmune diseases (43–45) and several bNAbs proved indeed to be auto- and polyreactive (46, 47). Since autoreactive B cells are counter-selected during B cell development (45), it has been speculated that bNAb development is normally blocked by immune checkpoints that can only be bypassed through chronic infection (48). In line with this, BCRs of chronically Hepatitis C virus (HCV)-infected individuals show increased CDRH3s lengths (i.e. potentially higher self-reactivity) in comparison to healthy controls (49). In addition, many potent HIV-1 bNAbs have been isolated after several years of infection (9, 42, 50), suggesting that a prolonged virus-antibody co-evolution is a requirement for their induction. Guided vaccine design attempts to mimic this co-evolution by serial immunizations with varying immunogens (51, 52). Yet it is currently unclear which bNAbs have the highest chance to be elicited and should thus be selected for these strategies. Previously, precursor frequencies of VRC01-like bNAbs and BG18 have been estimated in healthy individuals by CDRH3 similarity searches (52–54) and probabilities of distinct mutations have been determined for a subset of bNAbs (55). However, comprehensive methods and analyses that compare the combined probabilities of V(D)J recombination and overall mutation patterns across different bNAb classes are still lacking and it remains elusive how these probabilities are influenced by chronic infection.

Here, we performed unbiased next-generation sequencing (NGS) and learned probabilistic models for somatic point mutations and V(D)J recombinations on BCR repertoire data from 57 healthy individuals. We applied these models to determine and compare the generation probabilities of 75 bNAbs. By correlating probabilities with neutralization efficacies, we identified broad and potent candidate bNAbs that are more likely to be elicited than others and are thus particularly suited as targets for vaccination strategies. Finally, we sequenced 34 HIV-1- and 12 hepatitis C virus (HCV)-infected individuals, to infer repertoire characteristics and learn models to determine bNAb generation probabilities in the presence of chronic infections.

## Results

### Performing unbiased sequencing of the B cell receptor repertoire

In order to determine and compare the probability of developing an antibody with a specific sequence, we aimed to establish a pipeline for collecting unbiased BCR repertoire data from peripheral B cells and infer antibody sequence statistics with high confidence (**Figure 1 A**). To this end, we set up a sorting strategy to isolate naive or IgG-class-switched (i.e. antigen-experienced) B cells from peripheral blood mononuclear cells (PBMCs, **Supplementary Figure 1**) and developed a 5’-rapid amplification of cDNA ends (RACE)-based sequencing protocol including unique molecular identifiers (UMIs) for computational error correction (**Figure 1 A**).

**Figure 1:**
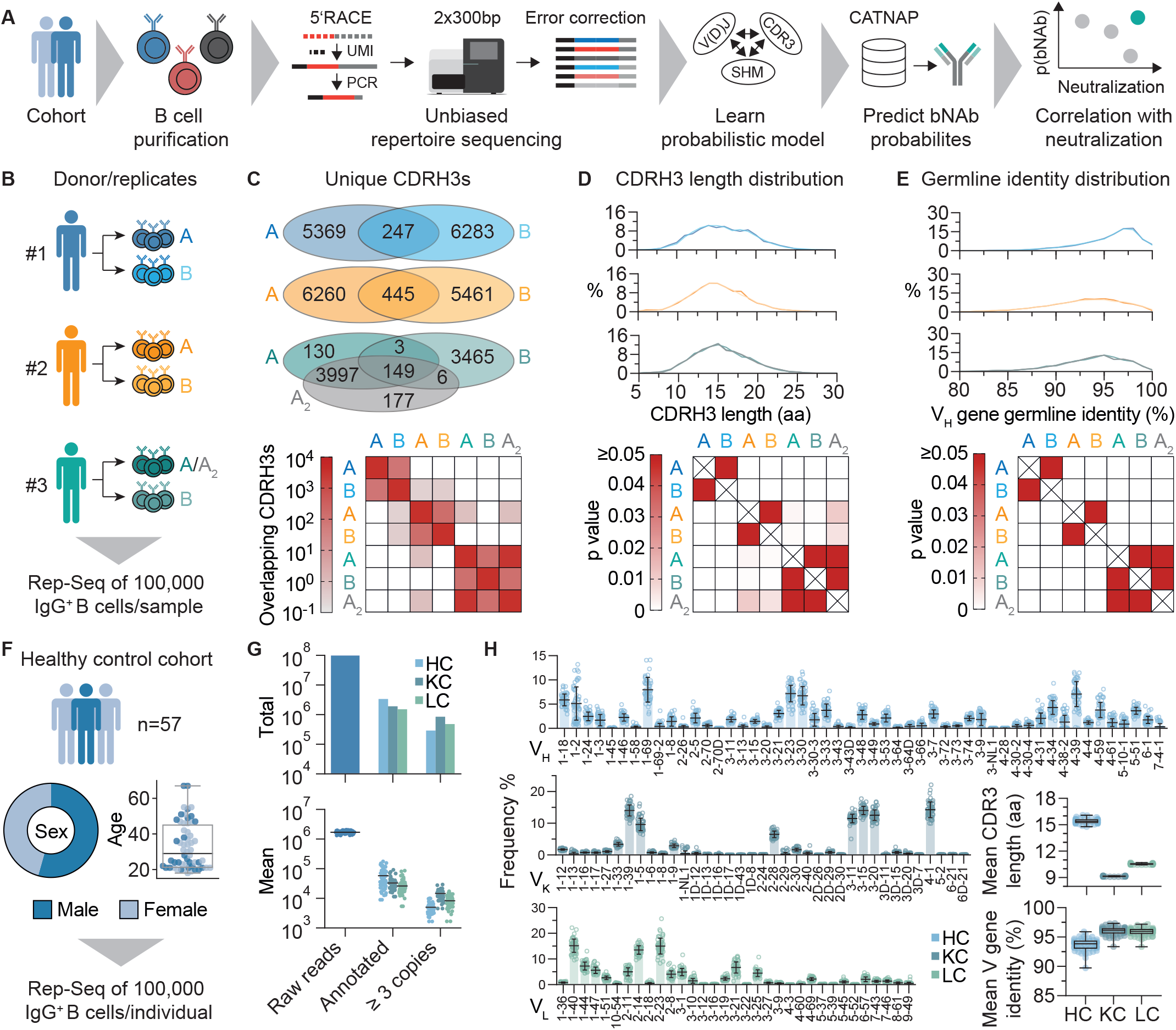
Overall approach and unbiased repertoire sequencing. (**A)** B cells are purified from peripheral blood and total mRNA is isolated. 5’ rapid amplification of cDNA ends (5’RACE) is performed to add unique molecular identifiers (UMI) and amplify variable regions. Amplicons are sequenced by 2×300 bp Illumina sequencing and UMIs are exploited for error correction. High-quality sequences are used to learn probabilistic models and predict probabilities of bNAb sequences, which are derived from the CATNAP database. Correlation of probabilities and neutralization allows for identifying highly probable and potent bNAbs. **(B)** Biological replicate samples of 100,000 lgG^+^ B cells were isolated from three healthy donors for sequencing pipeline validation. A_2_ represents a technical replicate of A (i.e., the same PCR product, for which library preparation and sequencing have been performed independently). **(C)** Overlap of unique CDRH3s between replicates from the samples in (B). Upper panel shows the total number of overlapping CDRH3s between biological and technical replicates. Lower panel shows mean CDRH3 overlap across all samples after 100 iterations of subsampling n=3623 CDRH3s from each dataset. **(D)** CDRH3 length distributions from samples in (B). Upper panel shows overlayed distributions for biological replicates. Lower panel shows p-values from Dunn post-hoc test after global Kruskal Wallis test. (E) V_H_ gene germline identity distributions from samples in (B). As for (D), upper panel shows overlayed distributions for biological replicates and lower panel p-values from Dunn post-hoc test after global Kruskal Wallis test. (F) Cohort sex and age distributions of n=57 healthy individuals for lgG^+^ repertoire sequencing. (G) Total and mean number of raw reads, annotated reads, and identical sequences (identified by the same UMI) that were found at least three times. (H) V gene distributions, mean CDR3 lengths and mean V gene germline identities for heavy, kappa, and lambda chains of all 57 individuals. CDR: Complementarity determining region; SHM: Somatic hypermutation; HC: heavy chain; KC: kappa chain; LC: lambda chain.

To test whether this sequencing approach yields sufficient high-quality reads, we analyzed biological duplicates of 100,000 IgG^+^ B cells from three healthy blood donors (**Figure 1 B**). From 1,354,097 to 2,552,903 raw reads per replicate, we reconstituted on average 6,121 IgG heavy, 18,868 kappa, and 12,000 lambda chains after filtering for error-corrected and productive sequences (**Supplementary Figure 2**, **Supplementary Table 1**). There was substantial overlap in unique CDRH3s within samples from the same individuals (on average 2.4 to 4.1% for biological replicates and 81% for the technical replicate) in comparison to the overlap between different donors (<0.02%, **Figure 1 C**). Moreover, repertoire features such as CDRH3 length distribution (**Figure 1 D**) or V_H_ gene germline identity (**Figure 1 E**) were indistinguishable between biological replicates, but significantly differed across individuals. To estimate the resolution of the sequencing approach, we spiked in varying concentrations (0.01 to 10%) of two B cell lymphoma (BCL) cell lines into naive B cells from a healthy donor. Spiked-in cells could be detected at all concentrations, including as few as 10 in a total of 100,000 cells (**Supplementary Figure 3**). Concluding that the sequencing pipeline yields reproducible and representative statistics, we collected blood samples from 57 healthy individuals that comprised 54% male and 46% female donors with a mean age of 33 years (**Figure 1 F**). By sorting and sequencing in total 5,700,000 IgG positive B cells we generated 96,825,014 raw reads, yielding high-quality productive sequences for 287,505 heavy, 839,776 kappa, and 476,835 lambda chains, with on average 5,044 heavy, 14,733 kappa, and 8,366 lambda chains per individual (**Figure 1 G**, **Supplementary Table 2**). Although sequence features were predictive of the repertoire’s origin (**Figure 1 D, E**), they are relatively conserved between individuals on a global level (**Figure 1 H**). In accordance with previous studies (56–58) we find particular V genes (such as V_H_1-69 or V_H_3-23) to be highly abundant in our cohort, CDR3 lengths to average around 15-16, 9, and 11 amino acids and average mutational loads to peak around 93, 96, and 96% nucleotide V gene germline identity for heavy, kappa, and lambda chains, respectively.

We conclude that the presented pipeline allows for inferring high-quality repertoire statistics from a starting material of 100,000 B cells, and that this sample is representative of the unique IgG BCR repertoire feature distributions of a single individual. On average, IgG repertoires show conserved sequence feature distributions that reflect different probabilities for specific sequence characteristics to develop during B cell development and maturation.

### Predicting bNAb probabilities for V(D)J recombination and somatic hypermutation

To predict the probabilities of developing HIV-1 specific bNAbs, we retrieved sequence and neutralization data for 75 HIV-1 neutralizing antibodies with varying neutralization breadths and potencies against a panel of 56 different HIV-1 strains (**Supplementary Figure 4** and **Supplementary Table 3**) from the CATNAP database (59).

The 75 antibodies target various sites on the envelope spike-protein (**Figure 2A,** adapted from (11, 60)) and cover geometric mean IC_50_ values from 0.005 to 16.991 μg/ml as well as neutralization breadths between 10.7 and 98.2%, both of which we combined into a single neutralization score, to compare and rank bNAbs by their overall neutralization efficacy (**Supplementary Table 4, Figure 2B**). Overall, the 75 bNAbs show broad distributions for V_H_ gene usage (**Figure 2C**), CDRH3 lengths (**Figure 2D**), and V_H_ gene germline identities (**Figure 2E**), which differ substantially from the averaged IgG memory B cell repertoire statistics of the 57 healthy individuals. Notably, separating bNAbs into binding classes demonstrates that individual bNAbs are not necessarily extreme in all features. CD4 binding site antibodies, for example, often incorporate V_H_1-2 or 1-46 and are highly mutated, while their CDRH3 lengths are within the range of the memory IgG reference distribution (**Figure 2 C**, **D**, and **E**, blue bars). V2-apex antibodies, on the other hand, typically exhibit long CDRH3s, but are also less mutated (**Figure 2 D**, and **E**, yellow bars). Both classes of antibodies can be found among the top neutralizing antibodies (**Figure 2B**).

**Figure 2:**
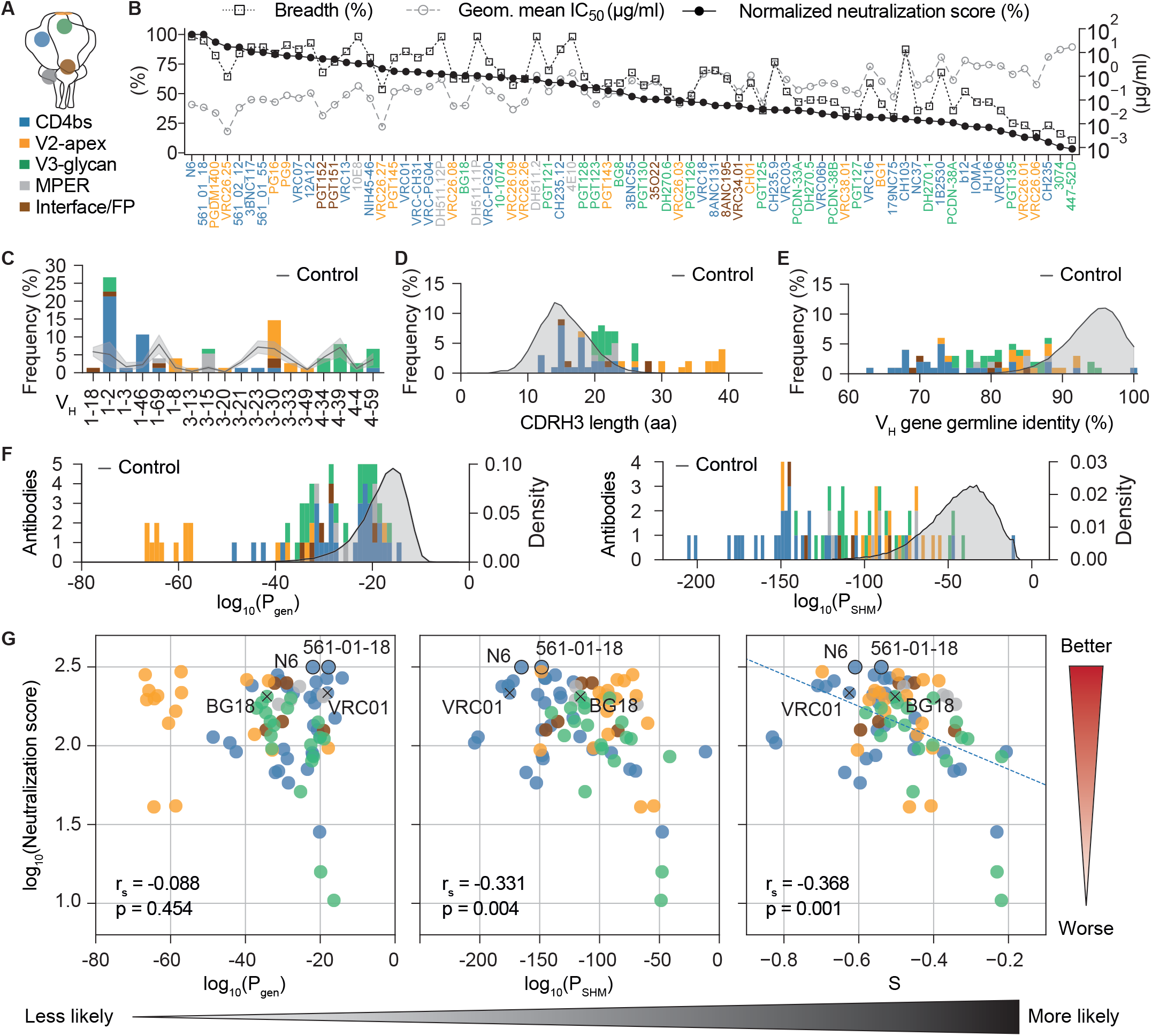
Neutralization efficacy and sequence characteristics of broadly neutralizing antibodies targeting HIV-1. (**A**) bNAb epitopes on the HIV-1 envelope spike. MPER: membrane proximal external region; CD4bs: CD4 binding site; FP: fusion peptide. (**B**) Breadth, geometric mean half inhibitory concentration (IC_50_) of neutralized strains, and normalized neutralization score for 75 bNAbs that have been tested against the same 56 HIV-1 strains. (**C**) Heavy chain V gene frequencies for the selected bNAbs. (**D**) Heavy chain CDR3 length distribution for the selected bNAbs in amino acids (aa). (**E**) Heavy chain V gene germline identity distribution for selected bNAbs. Controls in (C), (D) and (E) refer to sequence features from productive sequences of the 57 healthy individuals (Figure 1 F–H). (**F**) Heavy chain P_gen_ and P_SHM_ distribution for selected bNAbs derived by a model learned on productive sequences from 57 healthy individuals. Controls represent P_gen_ and P_SHM_ distributions for productive sequences from the 57 healthy individuals. (**G**) Correlation plots of bNAb neutralization scores against heavy chain P_gen_, P_SHM_ and a combined probability score S = c_1_log_10_(P_gen_) + c_2_log_10_(P_SHM_), which was derived by a linear regression (dashed line) with c_1_ = 4.0×10^-03^ and c_2_ = 3.2×10^-03^. Spearman correlation coefficients r_s_ and p values are given in the figure. Correlation coefficient and p value from linear regression for S are r = −0.446 and p = 6.031×10^-5^. Two near pan-neutralizing antibodies (N6 and 561-01-18) are highlighted by black outlines, two antibodies that have been used for structure-guided vaccine approaches (VRC01 and BG18) are highlighted by ‘x’. Gradients on the right and bottom show directions for increasing neutralization activity and generation probability, respectively.

While long CDRH3s and high levels of somatic mutations could in part explain the rare occurrence of bNAbs, they do not account for heterogeneity in gene segment selection probabilities, or for any sequence context, which is known to bias V(D)J recombination and somatic hypermutation (61). Moreover, simple descriptive statistics may suffer from undersampling of rare B cell sequences. We therefore applied the Inference and Generation Of Repertoires (IGoR) tool (62), to provide quantitative and comprehensive estimates for the probabilities of generating a given CDRH3 (P_gen_) and accumulating a unique pattern of point mutations (P_SHM_). IGoR infers a probabilistic model that accounts for the statistics of V(D)J usage as well as insertions and deletions in the junctional regions and sums over all generation scenarios consistent with the given BCR sequence to evaluate its overall generation probability. The SHM model learns the identities of mutated 5-mer subsequences. Importantly, estimating probabilities with models allows for overcoming limitations of sequencing depth through generalization. To investigate probabilities of bNAb sequence features after selection and affinity maturation, we took the combined quality-filtered and productive IgG sequences of all 57 healthy individuals to learn the models and determined P_gen_ and P_SHM_ for the 75 bNAb heavy chain sequences (**Figure 2F**, **Supplementary Table 5**). Almost all bNAbs show lower P_gen_ and P_SHM_ values (i.e., are less likely) than the median healthy IgG repertoire distribution, resembling the CDRH3 length and V_H_ gene germline identity comparison in **Figure 2D** and **E**. Indeed, there is a strong correlation between these sequence features and the probabilities (**Supplementary Figure 5**), suggesting that CDRH3 length and numbers of mutations are strong determinants of antibody probabilities. However, there are also antibodies with equal CDRH3 lengths (e.g. VRC03/DH511.11P) or similar amounts of SHM (e.g. VRC06b/3BNC117) but substantially different probabilities (over several log steps), highlighting the contribution of additional factors such as biases in V(D)J recombination and SHM (**Supplementary Figure 5**).

Given the broad distributions of P_gen_ and P_SHM_ for the different bNAbs, we asked whether they are correlated with neutralization efficacy, and whether we can identify highly potent bNAbs that are more likely to be generated. The neutralization score is not correlated with P_gen_ but slightly correlated with P_SHM_ alone for the 75 bNAbs (**Figure 2G** left and middle panel). Performing a linear regression that takes into account the logarithms of P_gen_ and P_SHM_ yields a combined probability score S (**Figure 2B** right panel, **Supplementary Table 5**) that is highly predictive of the neutralization score (r = −0.446 and p = 6.031×10^-5^ for the linear regression). Based on this regression, lower probability scores (i.e. less likely bNAbs) are correlated with better neutralization. Importantly, the correlation allows for identifying bNAbs that are highly potent, but easier to generate than others (**Figure 2G** right panel, upper right corner). Similar results for the correlation of P_gen_, P_SHM_, and the probability score S with the neutralization score were also obtained for light chains (**Supplementary Figure 6**, **Supplementary Table 5**).

We conclude that the linear combination of the logarithms of P_gen_ and P_SHM_ (i.e. the combined probability score S) is highly predictive of the neutralization capacity with the overall tendency of less likely bNAbs to be the most potent ones. In addition, our modeling approach not only confirms that bNAbs are in general unlikely outcomes of the B cell development, but also provides a framework for ranking bNAbs by their overall sequence probabilities.

### Global BCR repertoire features under chronic infections are similar to healthy repertoires with marginal differences

Since chronic infections have been described to interfere with B cell development and functionality (63), we wondered whether and to what extent they influence memory IgG BCR sequence characteristics. To answer this question, we sequenced BCRs from the IgG^+^ memory compartment of 34 HIV-1 and 12 hepatitis C virus (HCV)-infected individuals and compared them to the 57 healthy repertoires (**Figure 3 A**).

**Figure 3:**
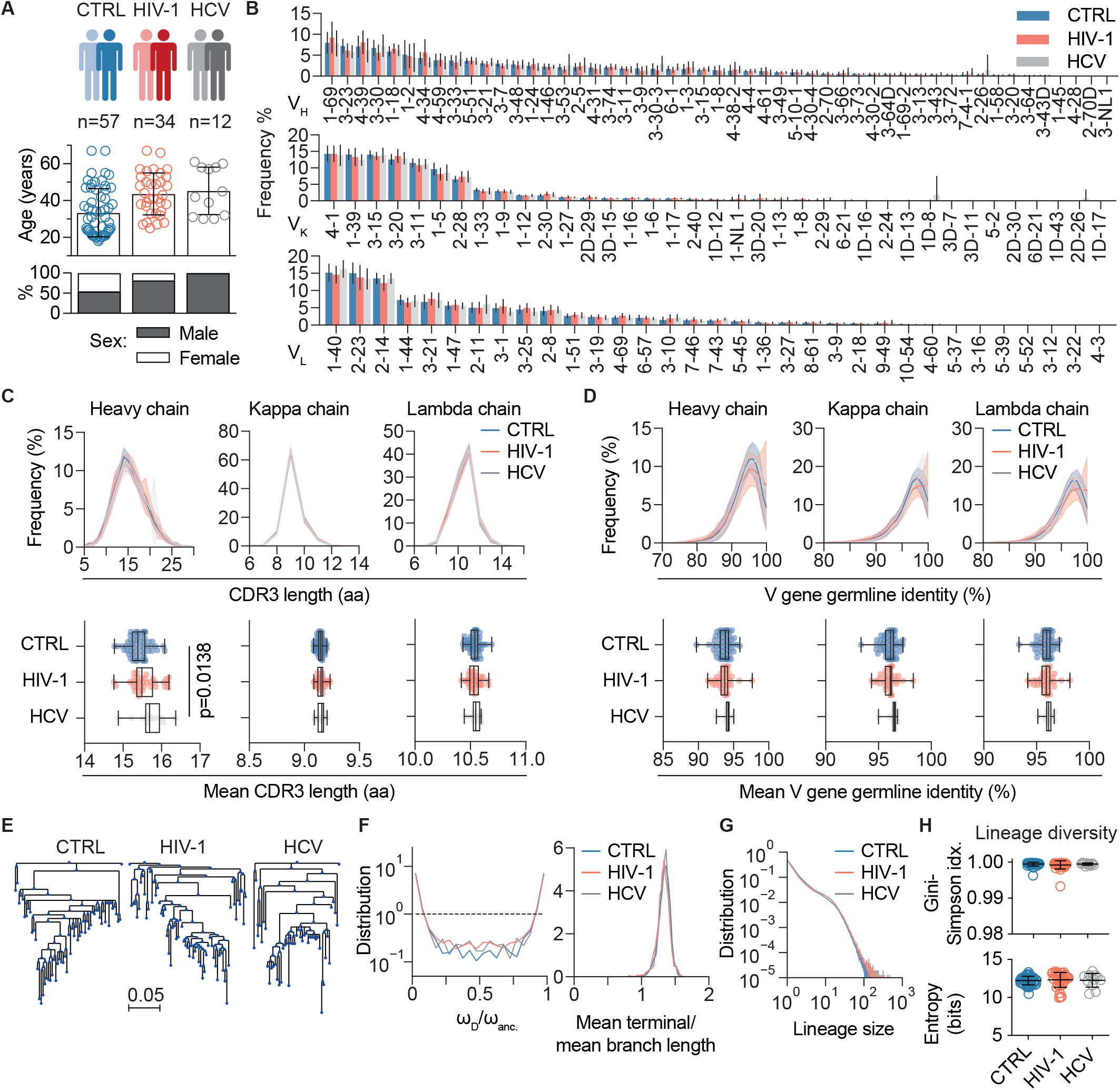
IgG heavy chain repertoire characteristics of healthy and chronically infected individuals. (**A**) Cohort overview with age and sex distributions. Single dots in age distributions represent individuals, bar graphs and error bars depict mean cohort age with standard deviations. (**B**) V gene distributions for heavy, kappa, and lambda chains. V genes were ordered by descending frequencies according to the healthy control group. Bars depict mean frequencies across individuals for each cohort, error bars show standard deviations. (**C**) Mean CDR3 length distributions for heavy, kappa, and lambda chains in amino acids (aa). Upper panel shows the mean CDR3 length distributions across individuals for each cohort as solid lines and standard deviations as shaded areas. Lower panel shows mean CDR3 amino acid length as dots for each individual and cohort statistics as box plots on top. (**D**) Mean V gene germline nucleotide identity distributions for heavy, kappa, and lambda chains. Representation of mean distributions (upper panel) and individual means (lower panel) as in C. One-way ANOVA with Tukey-HSD post-hoc test was performed on the means in (C) and (D). (**E**) Representative examples of lineages for the different cohorts based on heavy chain sequences. (**F**) Skewedness distribution of lineage trees (left panel, dashed line = neutral) and mean terminal/mean branch length distribution for lineage trees. (**G**) Lineage size distribution for the three cohorts. (**H**) Lineage diversity determined by Gini-Simpson index and entropy. CTRL = healthy control.

Processing and filtering of 54,628,577 HIV-1 and 19,767,705 HCV raw reads yielded in total 1,382,301 (HIV-1) as well as 486,488 (HCV) heavy and light chain sequences (**Supplementary Tables 6 and 7**). Overall, sequence characteristics, including V gene segment usage, CDR3 length, and V gene germline identity, were comparable to the healthy control cohort for both chain types (**Figures 3B, 3C, 3D**), although HCV-infected individuals showed slightly longer mean CDRH3 lengths (**Figure 3C**). Similarly, no obvious differences were observed in B cell lineage tree structures (**Figure 3E**) with respect to skewedness (**Figure 3F**), size distribution (**Figure 3G**), or diversity (**Figure 3H**). Since antiretroviral therapy (ART) is suppressing virus replication and thus may dampen anti-HIV-1 immune responses, we stratified the HIV-1 cohort by treatment (**Supplementary Figure 7**). Whereas repertoires from treated HIV-1-infected individuals were indistinguishable from healthy individuals, we found a substantially higher fraction of V_H_ genes 1-69 and 4-34 in untreated HIV-1-infected individuals, which did not reach significance after correcting for multiple testing. In addition, mean CDRH3 lengths were slightly but significantly longer in untreated individuals. Mean V gene germline identities did not differ significantly, although the untreated subgroup contains one individual with noticeably less somatic mutations, which is mainly responsible for the visible shift in the V gene germline identity distributions (**Figure 3D**, **Supplementary Figures 7**). We also measured serum-derived poly-IgG neutralization breadth against the 12-strain global panel (64) for 33 of the 34 HIV-1-infected individuals (**Supplementary Table 8**). Of note, broad serum neutralization is found in ART-naive (untreated) as well as ART-treated individuals of our cohort (**Supplementary Table 8**). Stratifying BCR repertoires by neutralization breadth into high (≥66%), intermediate (<66% and ≥33%), or low (<33% and >0%) breadth and no neutralization (0%) did not yield any evidence that serum neutralization breadth is linked to relevant global changes in BCR repertoire features (**Supplementary Figure 8**). Neither of the stratifications had any influence on the lineage characteristics (data not shown).

We conclude that V gene usages and CDRH3 lengths are marginally skewed in untreated, chronically infected individuals. However, on average, repertoire features remain almost constant between the cohorts.

### Equal probabilities for bNAb development in healthy and chronically infected individuals

To investigate whether chronic infection shifts the probability to develop distinct bNAb sequences, we used the IgG repertoire data from HIV-1 and HCV-infected individuals to train IGoR and learn models for P_gen_ and P_SHM_.

Whole repertoire P_gen_ distributions of both cohorts were almost identical to the healthy control data for heavy, kappa, and lambda chains (**Figure 4A**). Repertoire P_SHM_ distributions on the other hand were slightly shifted towards less negative values for HCV- and HIV-1-infected individuals (**Figure 4B**), which resembles the increase in V_H_ gene germline identity (i.e. less somatic mutations) that was already observed in the repertoire data (**Figure 3D**). We speculated that marginal differences in these distributions could still influence the probability for certain rare V(D)J recombinations or mutational patterns of distinct bNAb sequences. Therefore, we calculated P_gen_ and P_SHM_ for the individual 75 bNAbs with the models that have been trained on the repertoire data from infected individuals (**Supplementary Table 5**). When comparing those to the values that have been inferred from the healthy cohort, there is only little deviation in P_gen_ or P_SHM_ except for three probabilities that deviate more than 25% from the healthy cohort (**Figure 4C and D**). Interestingly, P_gen_ for the light chain of 10-1074 is >5 log-fold higher (i.e., more likely) in healthy individuals than in both chronically infected cohorts, suggesting that the selection of this particular light chain sequence is somehow hampered by chronic infections. There is however no tendency for a whole antibody epitope group to be favored in one or the other cohort. To finally compare the overall probabilities of individual bNAbs between the cohorts, we finally calculated the combined probability score (S) using the cohort-specific P_gen_ and P_SHM_ values together with the coefficients from the healthy repertoire linear regression model of Figure 2G (**Supplementary Table 5**). In line with the similar P_gen_ and P_SHM_ values (**Figure 4C** and **D**), combined probability scores were almost identical between healthy and infected individuals (**Figure 4E**). Despite the slight differences in the repertoire characteristics of untreated individuals (**Supplementary Figure 7**), treatment status had no influence on bNAb probability scores (**Figure 4F**). Similarly, we found no global change in probability scores when stratifying by serum neutralization breadth, except for three less probable chains in high (CH103 light chain), intermediate (10-1074 heavy chain), and low breadth subgroups (179NC75 heavy chain, **Figure 4G**). Taken together, these results suggest no substantial differences in the overall probability of bNAb generation in the absence or presence of chronic HIV-1 or HCV infection.

**Figure 4:**
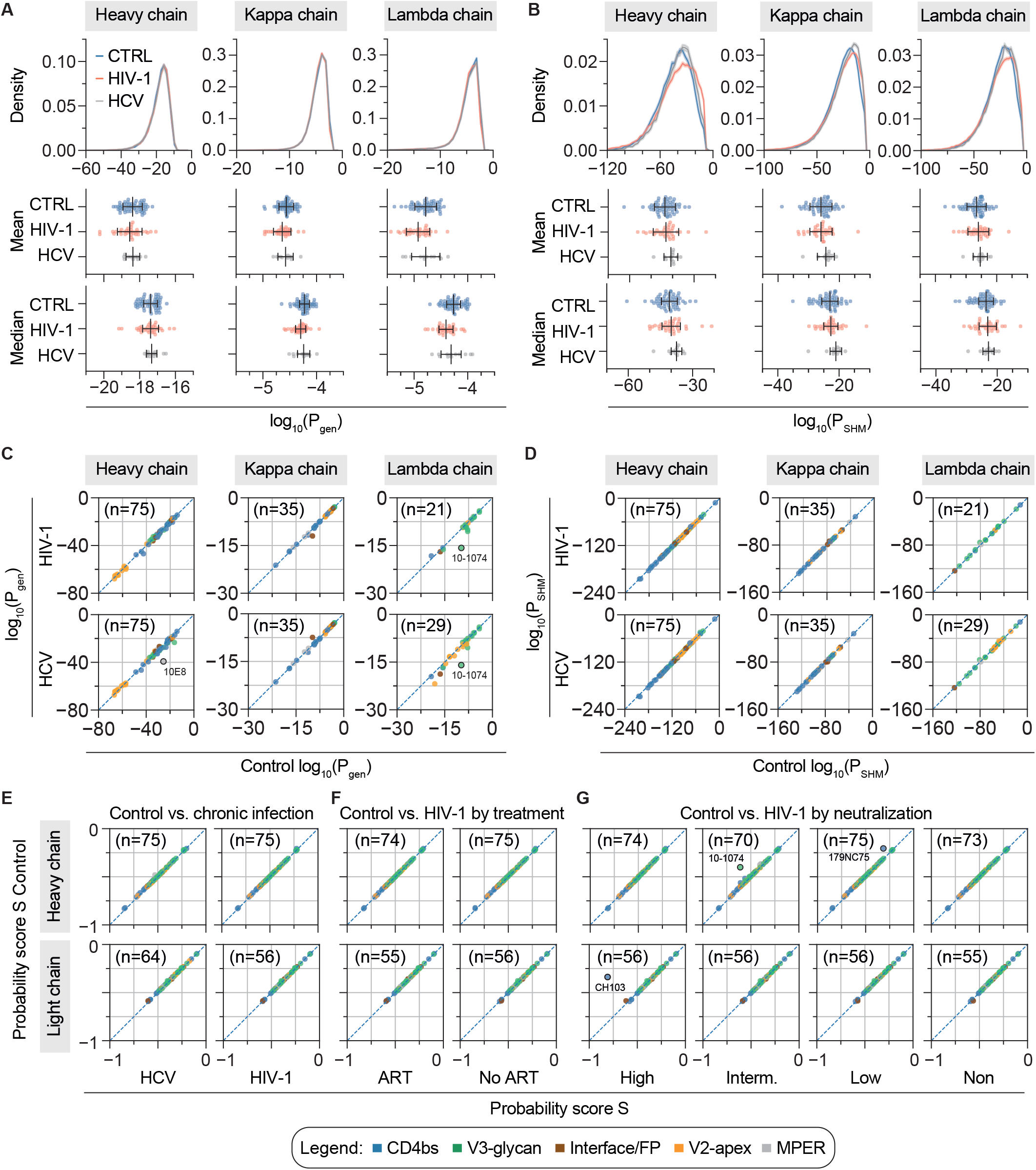
Probability distributions and bNAb probability scores across cohorts. (**A**) P_gen_ distribution as well as mean and median P_gen_ for heavy and light chains derived from healthy control (CTRL), HIV-1-infected, and HCV-infected cohorts. (**B**) P_SHM_ distributions for heavy and light chains as in (A). (**C**) Correlation plots of P_gen_ for neutralizing antibody heavy, kappa, and lambda chains derived from either healthy control (x-axis) or chronically infected cohorts (HIV-1, HCV, y-axis). (**D**) Correlation plots of P_SHM_ as in (C). Highlighted antibodies in (C) and (D) deviate >25% of the range of all observed values in comparison to the healthy cohort. (**E**) Correlation plot for bNAb heavy and light chain probabilty scores derived from HCV or HIV-1 cohorts (x-axis) in comparison to healthy control (CTRL, y-axis). Cohort-specific P_gen_ and P_SHM_ of bNAbs were used to calculate the bNAb probability scores S using coefficients from the linear regression on healthy individuals (Figure 3G). The dashed line represents identity. (**F**) Comparison as in (E) with the HIV-1 cohort stratified by antiretroviral therapy (on ART, n=22; off ART, n=12). (**G**) Comparison as in (E) with the HIV-1 cohort stratified by serum neutralization breadth against the global HIV-1 panel into high (≥66%, n=8), intermediate (≥33%, n=5), or low (>0%, n=16) breadth and no neutralization (0 %, n=4) to recalculate P_gen_ and P_SHM_ of bNAbs for these cohorts. Serum neutralization data for one individual was not available. ART: anti-retroviral therapy. Numbers in brackets in (C - G) indicate bNAbs for which P_gen_/P_SHM_ and scores could be inferred.

## Discussion

Broadly HIV-1 neutralizing antibodies can effectively prevent and treat HIV-1 infections in animal models (16, 17, 19, 65) and are considered promising tools for HIV-1 prevention and therapy in humans (66, 67). However, bNAbs have only been identified in a minor fraction of HIV-1-infected individuals and to elicit highly broad and potent bNAbs by vaccination remains a yet unreached goal (11, 68, 69). The presence of high amounts of somatic mutations, insertions/deletions, or exceptionally long CDRH3s in bNAbs raised the questions of (i) whether these outstanding features explain their rareness, (ii) whether rareness is correlated with neutralization capacity, and (iii) if chronic infection is a prerequisite for their induction. By learning models from IgG^+^ memory BCR sequences of 57 healthy individuals, we estimated the probability for specific V(D)J recombinations (P_gen_) and for accumulating patterns of somatic point mutations (P_SHM_) in a representative set of HIV-1-neutralizing antibodies. We also tried to include insertions and deletions (indels). However, it was not possible to build a robust model, since their frequencies were too low in the repertoire data. In line with previous studies, we found that numerous bNAbs accumulate improbable mutation patterns (55, 70). Our analysis illustrates that the degree of improbable mutations is correlated with the binding site and is not a common feature of bNAbs per se. Similarly, we identified a subset of antibodies with highly improbable CDRH3 recombinations. These mainly comprise V2 apex antibodies that require long CDRH3s to penetrate the envelope glycan shield (71– 73), but also some CD4bs antibodies with average CDRH3 lengths. These results support the hypothesis that bNAbs are rare, because some of their sequence features are unlikely to develop.

Previously, ongoing affinity maturation as well as the mean frequency of somatic mutations have been correlated with neutralization breadth and potency of bNAb lineages (74, 75). In line with this, we detected a weak correlation for the neutralization power of monoclonal bNAbs and SHM, looking not only at the frequency but also at the probability to accumulate a specific mutation pattern (i.e., P_SHM_). Notably, we found that a linear combination of the logarithms of P_gen_ and P_SHM_ is predictive of bNAb neutralization efficacy, suggesting that bNAbs tend to be better, the more unlikely they are. Importantly, the correlation allows for identifying bNAbs that are highly potent but on average more likely than other, similarly potent ones. These highly potent and probable antibodies include the above-mentioned V2 apex antibodies, which in contrast to their improbable CDRH3 showed more probable patterns of somatic hypermutations. Interestingly, V2 apex antibodies have been reported to occur relatively early during infection (76–78). This suggests that the selection of rare precursors outperforms the accumulation of distinct SHM in timing and that this bNAb subclass could thus be a preferential target for vaccination strategies.

In addition to healthy individuals, we also sequenced the repertoires of 34 HIV-1- and 12 HCV-infected individuals. In concordance with previous studies (49, 79), we detected marginally longer mean CDRH3 lengths in HCV-infected as well as HIV- 1-infected individuals that do not re ceive antiretroviral therapy. Similar to the findings from Roskin et al. (79), we also detected an increase of the inherently auto-reactive gene segment V_H_4-34 in our ART-naive HIV-1-infected subgroup. However, we do not see substantial differences in bNAb sequence probability scores in the IgG^+^ memory compartment when averaging over individuals in the cohorts. This suggests that chronic infection itself does not alter the probabilities to develop specific bNAbs substantially in any direction, even if it leaves traces of particular sequence features within the whole IgG repertoire. Of note, we detected high variability in the repertoire composition within each cohort. As a consequence, some individuals might still have better chances of developing certain bNAb classes because of other predispositions, e.g. by carrying specific gene segment alleles that encode for critical contact sites, as it has been previously demonstrated for VRC01-like bNAbs (53).

Taken together, we present a framework for assessing the overall probability to develop an antibody with a specific CDR3 motif and pattern of point mutations. Our modeling approach on HIV-1-neutralizing antibodies quantitatively confirms that bNAbs are unlikely outcomes of the B cell evolution within a host due to individual or a combination of improbable sequence features. Using neutralization data and probability scores, we demonstrate that the more unlikely bNAbs tend to have a higher neutralization efficacy. Moreover, this approach allows us to identify potent antibodies with higher chances to be elicited by vaccination. Finally, the data suggest that chronic infection has no impact on the generation of bNAb sequences within the IgG^+^ B cell memory compartment, fostering the hope that a potent vaccine should be able to elicit bNAbs in healthy individuals.

## Materials and Methods

### Data and Material Availability

FASTQ files for all repertoires have been deposited at the Sequence Read Archive (SRA) with accession numbers SAMN29624595-713 (BioProject Accession Number PRJNA857338). IGoR is freely available from a github repository (https://github.com/statbiophys/IGoR). Requests for materials or additional code should be directed to the corresponding author and may be subject to restrictions based on data and privacy protection regulations and/or may require a Material Transfer Agreement (MTA).

### Sample collection

Samples were obtained under a study protocol approved by the Institutional Review Boards of the University of Cologne, University of Bonn, and the respective local IRBs (study protocols 16-054 and 017/16). All participants provided written informed consent and were recruited at hospitals or as outpatients.

### Serum IgG isolation

Serum samples from HIV-1-infected individuals were heat-inactivated at 56 °C for 40 min and incubated with Protein G Sepharose (GE Life Sciences) overnight at 4 °C. IgGs were eluted from chromatography columns using 0.1 M glycine (pH = 3.0) into 0.1 M Tris (pH = 8.0). Buffer was exchanged to PBS through Amicon 30 kDa spin membranes (Millipore). Concentrations of purified IgGs were determined by UV/Vis spectroscopy (A280) on a Nanodrop 2000 and samples were stored at 4°C.

### Serum IgG neutralization test

Neutralization assays with serum IgGs against the 12-strain “global” virus panel, were performed in 96-well plates as previously described (9, 80). In brief, 12 HIV-1 pseudovirus strains were mixed with 1:2 serial dilutions of purified IgG (1 mg/ml starting concentration, 8 dilutions) and incubated for 1 h at 37°C. TZM-bl cells were added (10^4^ per well) with DEAE-dextran at a final concentration of 10 μg/ml and incubated for 2 days. Luciferin-containing lysis buffer was added and after 2 min incubation samples were resuspended and luminescence was measured with a luminometer (Berthold TriStar^2^ LB942). For IC_50_ determination, the background signal (non-infected TZM-bl cells) was subtracted and IgG concentrations resulting in a 50% RLU reduction compared to untreated virus control wells were determined by using murine leukemia virus (MuLV)-pseudotyped virus as a control for unspecific activity. All samples were tested in duplicates.

### Isolation of B cells and RNA Isolation

PBMCs were isolated by standard density gradient centrifugation using Histopaque (Sigma Aldrich) and LeucoSep tubes (Greiner Bio-one). Cells were stored at −150°C in 90% (v/v) FBS (Sigma Aldrich) and 10% (v/v) DMSO (Sigma Aldrich). Plasma was collected and stored separately at −80°C. B cells were enriched from PBMCs with CD19 microbeads (Miltenyi Biotec). CD19^+^ cells were stained with anti-human AF700-CD20 (BD), APC-IgG (BD), PE-Cy-7-IgD (BD), FITC-IgM (BD), PerCP-Cy5.5 (BD) or PE-CD27 (BD) and DAPI (Thermo Fischer). CD20^+^IgG^+^ or CD20^+^IgD^+^IgM^+^CD27^-^IgG^-^ B cells were sorted into FBS (Sigma-Aldrich) using a FACSAria Fusion cell sorter (Becton Dickinson). RNA-Isolation was performed using the RNeasy Micro Kit (Qiagen) on a QiaCube (Qiagen) instrument.

### RT-PCR and next generation sequencing

BCR repertoire sequence data was generated by template-switch RT-PCR. cDNA was generated from 10 μl RNA according to the SMARTer RACE 5’/3’ manual using SMARTScribe Reverse Transcriptase (Takara) and a self-designed template-switch oligo (AGGGCAGTCAGTCGCAGNNNNWSNNNNWSNNNNWSGCrGrGrG). cDNA was diluted with 10 μl Tricine-EDTA buffer (Takara) according to the manual. Heavy and light chain variable regions were pre-amplified from 5 μl cDNA each by PCR with 1 μM forward primer (CTGATACGATTCACGCTAGGGCAGTCAGTCGCAG) and 0.33 μM constant region-specific reverse primers (IgM: ATGGAGTCGGGAAGGAAGTC, IgG: AGGTGTGCACGCCGCTGGTC, IgK: GGTGACTTCGCAGGCGTAG, IgL: GCCGCGTACTTGTTGTTGC) in a 30 μl reaction with Q5 DNA polymerase (New England Biolabs). Cycling conditions were one cycle 98°C/30 s, four cycles 98°C/10 s and 72°C/30 s, four cycles 98°C/10 s, 62°C/30 s (IgG/IgM) or 68°C(IgK/IgL)/30 s, and 72°C/30 s, as well as a final extension cycle at 72°C/5 min. PCR products were purified with a PCR clean-up Kit (Macherey Nagel) with a 1/6 dilution of NTI binding buffer in RNAse-free water. Samples were eluted in 15 μl elution buffer (Macherey Nagel). Heavy and light chain amplicons were enriched from 5 μl purified pre-amplification product by a nested PCR with 0.33 μM forward primer (IgG: NNNNNCACGCTAGGGCAGTCAG; IgM: NNNNNNCACGCTAGGGCAGTCAG; IgK/IgL: NNNNCACGCTAGGGCAGTCAG) and 0.33 μM nested reverse primers (IgG:NNNNNSGATGGGCCCTTGGTGGARGC; IgM:NNNNGGTTGGGGCGGATGCACTCC;IgK:NNNNNNGGGAAGATGAAGACAGATGGT, IgL:NNNNNNGGGYGGGAACAGAGTGACC) with Q5 DNA Polymerase (New England Biolabs) in a 100 μl reactions. PCR conditions were one cycle 98°C/30 s, five cycles 98°C/10 s and 72°C/30 s, five cycles 98°C/10 s and 70°C/30 s, 17 cycles 98°C/10 s, 68°C/30 s (IgG/IgM) or 62°C(IgK/IgL)/30 s, and 72°C/30 s, one final extension cycle at 72°C/5 min. Amplicons were separated on a 1% Agarose gel, purified with a PCR and Gel Clean Up Kit (Macherey Nagel) and subjected to library preparation and Illumina MiSeq 2 x 300 bp sequencing (v3) at the Cologne Center for Genomics sequencing core facility.

### NGS data processing and sequence annotation

Data pre-processing was performed with a Python (v.3.6)-based pipeline, including python packages BioPython (v.1.78), pandas (v.0.23.4), Numpy (v.1.19.2), Matplotlib (v.3.3.4), and Levensthein (v.0.12.0). Raw NGS reads were filtered for a mean Phred score of 25 and read-lengths of at least 250 bp. Reads were annotated with IgBLAST (81) to identify read-orientation and CDR3 sequence. Reads were then grouped by the UMI and wrongly assigned reads (collisions) were identified by comparison of V gene calls and a second 18-nucleotide molecular identifier within the CDR3. Reads were then aligned with Clustal Omega (82, 83) and consensus sequences were generated by taking into account the quality score-weighted base-calls for each position. The 2x 300pb consensus read pairs were then aligned and combined using the AssemblePairs.py script form the pRESTO toolkit with a minimal overlap of 6 nucleotides (84). Annotation of pre-processed reads was performed by using BLAST (v.2.9) and IgBLAST(v.1.13) (81). Reference templates for all functional heavy and light chain V(D)J genes were retrieved from the IMGT database (85, 86) (265 IGHV, 30 IGHD, 13 IGHJ, 66 IGKV, 9 IGKJ, 70 IGLV, and 7 IGLJ at the time point of this study). After sequence annotation with IgBLAST, sequences were filtered to include complete V/J annotations covering at least 250 nucleotides of the V gene as well as productive sequences (i.e., no STOP codon and in-frame CDR3 recombination, as determined by IgBLAST) only. To minimize the influence of sequencing and PCR errors, annotated sequences were only used for downstream analyses, when their UMIs were initially found in at least three reads (high-quality reads). For learning P_gen_ and P_SHM_ models, sequences were additionally filtered out, when they contained gaps (i.e. insertions/deletions).

### Antibody selection

Antibody information was retrieved from the CATNAP database (59). At the time of this study, the available antibody dataset comprised 507 entries, from which non-human antibodies, polyclonal antibodies, antibody mixtures, as well as mutants/chimeras and non-antibody proteins were removed to yield a final set of 291 antibodies. The 291 antibodies were tested against different or only partially overlapping viral panels (such as the 12-strain ‘global’ panel (64) or the 118-strain multi-clade panel (87)), making it difficult to compare them in terms of breadth and potency (88). Of the 291 antibodies, 50 have been tested against the global panel and 19 against the 118 multi-clade panel. As a trade-off between number of antibodies and accuracy of breadth and potency, we selected a subset of 56 strains from the 118 multi-clade panel, which resembles its clade distribution, its tiered categorization, and yields similar values for breadth and potency (**Supplementary Figure 4**, **Supplementary Table 3**). 75 out of 291 antibodies have been tested against this 56 strain panel, are published with complete heavy chain nucleotide sequences (including 69 light chains) and neutralized at least 1 of the 56 strains. Heavy and light chains were annotated with the same IgBLAST/IMGT-based pipeline as the NGS data. Five light chains could not be annotated and were excluded.

### Determination of antibody breadth, mean potency and neutralization score

IC_50_ values for individual antibody-virus combinations were derived from the CATNAP database (59) as the geometric mean of all published IC_50_ values (i.e. from different studies). IC_50_ values below the detection limit were set to an arbitrary threshold of 100 μg/ml. The mean potency of an antibody against a viral panel was determined by the geometric mean of all geometric mean IC_50_ values that were below the arbitrary threshold of 100 μg/ml (also known as the mean potency of all neutralized strains). Antibody breadth was defined as the percentage of neutralized strains (i.e. geometric mean IC_50_ above the arbitrary threshold of 100 μg/ml) to the total number of strains tested and is given as percentage. To determine the combined neutralization score, we took all IC_50_ values that were below the threshold of 100 μg/ml for each antibody and sorted them from lowest to highest. We then divided the maximum coverage (100%) into 56 equally spaced increments (~1.786% coverage per strain), and assigned cumulative increments to the sorted IC_50_ values, i.e. the lowest IC_50_ is assigned to a coverage of 1.786%, the second lowest to 3.572%, and so on. Finally we plotted the log10 of the sorted IC_50_ values against the cumulative coverage and determined the area under the curve. Low IC_50_ values (i.e. highly potent antibodies) will increase the area by shifting the curves to the left, while higher breadth will increase the area by shifting the plateau of the curve to the top. For the ranking of bNAbs in **Figure 2B**, the neutralization score is given as percentage of the highest score among the 75 antibodies. For correlation of probabilities and neutralization (**Figures 2 and 4**), the log_10_ of the neutralization score was used.

### IGoR inference and prediction of bNAb probabilities

Models for the probability of generation (P_gen_) and somatic hypermutation (P_SHM_) were learned by using the Inference and Generation Of Repertoires (IGoR) tool (62). To reduce uncertainty in model inference and focus on comparison between cohorts, we pooled all productive sequences from all patients in a cohort. Productive sequences were chosen to explore the effect of possible selection effects on P_gen_ and P_SHM_ of a given BCR sequence. The somatic hypermutation (SHM) model is 5-mer based, taking into account the mutated position and its two neighbors on both sides. The model takes into account the full composition of these 5-mers and is context dependent. P_gen_ generation models were consistent with models inferred on previously published repertoires (58, 89) (data not shown). Indels in bNAbs were reverted to the most likely V and J templates according to the IgBLAST annotation before model building to improve alignment by IGoR. Masking of indels has no influence on P_gen_ or P_SHM_. To test correlations between the neutralization score and the overall likelihood of seeing a given bNAb, we defined a score S for each bNAb as S = c_1_ log_10_ P_gen_ + c_2_ log_10_ P_SHM_. Coefficients c_1_ and c_2_ were determined by linear regression of score S versus the log_10_ of neutralization score.

### Lineages and trees

After V, D and J annotation, unique sequences were partitioned into lineages. First, the sequences were grouped into classes of identical V and J genes and equal CDR3 length. Within each class, lineages were identified using single linkage clustering with a fixed threshold of 90% CDR3 nucleotide identity. Tree length was then estimated as the total number of unique mutations found in a given lineage (a lower bound on the true tree length). For productive lineages of more than 50 unique sequences tree topologies and branch lengths were inferred using RAxML with the GTRGAMMA model of nucleotide substitution (90). Germline V and J genes were provided as an outgroup to aid the inference. To quantify the asymmetry of the phylogenies within each cohort we examined the distributions of two indices of imbalance, following (91). We compare the weight of the common ancestor of the lineage w_anc_ with the weights of its immediate descendants, w_D_. The weight of a node is the number of leaves that stem from it and the ratio w_D_/w_anc_ quantifies the imbalance at the first branching. Additionally, we estimated the distribution of the ratio of mean terminal branch length to the mean length of all branches.

### Quantification and Statistical Analysis

Flow cytometry analysis and quantifications were done with FlowJo10 software. Statistical analyses were performed using GraphPad Prism (v8), Microsoft Excel for Mac (v14.7.3), Python (v3.6.8), and R (v4.0.0). For the identification of overlapping clonotypes in healthy individuals a maximum of one amino acid length difference and three or less differences in absolute amino acid composition of CDR3s were considered as similar. To calculate the CDRH3 sequence overlap between replicates in the control experiment (**Figure 1 C**), a random sample of n=3623 (i.e. the size of the smallest dataset) was drawn from all unique CDRH3s of each set and the overlap (i.e. identical CDRH3 sequences) between these random samples was determined. Sampling and overlap determination was repeated 100 times and the mean overlap over all iterations was reported. Testing for significant differences between CDR3 length distributions and V gene germline identity distributions (**Figure 1 D** and **E**) was performed by global Kruskal wallis tests (stats.kruskal, Scipy v.1.5.2) and Dunn post-hoc tests (posthoc_dunn, scikit_posthocs v. 0.4.0 with ‘holm’ method for p value adjustment). To estimate the fraction of spiked-in lymphoma B cells among naive B cells from a healthy donor (**Supplementary Figure 3**), the pairwise Levenshtein distance of all re-constituted CDRH3s was determined (python-Levenshtein v.0.12.2) in comparison to the most frequent cell line CDRH3 (experimentally determined by sequencing 24 single B cells for each cell line, data not shown). The observed distance frequencies revealed a bimodal distribution from which all comparisons with <4 amino acid distance (first peak) were counted as a B cell line CDRH3 and divided by all comparisons to get the fraction. For cluster analysis of the mixed cell sample (**Supplementary Figure 3**), a network analysis was performed with the networkx python package (v.2.2). Each node represents a unique CDRH3. The node size is proportional to the frequency among all identified CDRH3s and nodes are connected, if they share at least 75% of their CDRH3 amino acid sequence. Nodes are colored according to the cell lines if they share at least 75% of the CDRH3 amino acid sequence with a cell line. Phylogenetic trees of viral panels (**Supplementary Figure 4**) were generated and illustrated from aligned viral sequences from the CATNAP database (59) with Geneious Prime software (v.2020.2.4, Jukes-Cantor genetic distance model and Neighbor-Joining method with 100 bootstrap replicates). Testing for significant differences in mean CDR3 lengths and mean V gene germline identities (**Figure 3 C** and **D**, **Supplementary Figures 7** and **8**) was performed by one-way ANOVA (stats.f_oneway, Scipy v.1.5.2) followed by Tukey-HSD post hoc test (stats.multicomp.pairwise_tukeyhsd, statsmodels v.0.12.2). Differences in V gene distributions (**Figure 3 B**, **Supplementary Figures 7** and **8**) were investigated by individual Kruskal-Wallis tests (stats.kruskal, Scipy v.1.5.2) for each V gene with Bonferroni correction for multiple testing. In the case of significant differences in the global test, a Dunn post hoc test (posthoc_dunn, scikit_posthocs v.0.4.0 with ‘holm’ method for p value adjustment) was performed for subgroup analysis.

## Supporting information

SI_Kreer_Lupo_Ercanoglu_et_al

## Acknowledgments

We thank all members of the Klein Laboratory for support and helpful discussion, Viera Kovacova, Antonios Papadakis, Lukas Maas, and Milos Nikolic for advice on data evaluation and NGS pipeline programming, Peter Nürnberg, Janine Altmüller, and Christian Becker from the Cologne Center for Genomics (CCG) for sequencing support, Michael Lässig and Christa Stitz for support within the CRC1310; This work was funded by grants from the German Research Foundation (DFG; CRC 1279 to F.K.; CRC 1310 to F.K., C.K., A.M.W., A.N., J.G., A.B.), the German Center for Infection Research (DZIF to F.K.), the European Research Council (ERC-StG639961 to F.K; COG 724208 to A.M.W.), the Agence Nationale de la Recherche (ANR-19-CE45-0018 “RESP-REP” to T.M.), CAREER Award from the National Science Foundation, Grant No. 2045054 (A.N.), and the MIRA award from the National Institutes of Health, Grant No. 1R35GM142795-01 (A.N.).

## References

1. F. W. Alt, Y. Zhang, F. L. Meng, C. Guo, B. Schwer, Mechanisms of programmed DNA lesions and genomic instability in the immune system. Cell 152, 417–429 (2013).

2. G. Teng, F. N. Papavasiliou, Immunoglobulin somatic hypermutation. Annu. Rev. Genet. 41, 107–120 (2007).

3. N. Matthews, Annual Review of Immunology. J. Med. Genet. 23, 284–285 (1986).

4. M. H. Malim, M. Emerman, HIV-1 sequence variation: Drift, shift, and attenuation. Cell 104, 469–472 (2001).

5. A. Nourmohammad, J. Otwinowski, J. B. Plotkin, Host-Pathogen Coevolution and the Emergence of Broadly Neutralizing Antibodies in Chronic Infections. PLoS Genet. 12, 1–23 (2016).

6. J. F. Scheid, et al., Sequence and Structural Convergence of Broad and Potent HIV Antibodies That Mimic CD4 Binding. Science 333, 1633–1637 (2011).

7. L. M. Walker, et al., Broad and Potent Neutralizing Antibodies from an African Donor Reveal a New HIV-1 Vaccine Target. Science (80-.). 326, 285–289 (2009).

8. X. Wu, et al., Rational Design of Envelope Identifies Broadly Neutralizing Human Monoclonal Antibodies to HIV-1. Science (80-.). 329, 856–861 (2010).

9. P. Schommers, et al., Restriction of HIV-1 Escape by a Highly Broad and Potent Neutralizing Antibody. Cell 180, 471–489.e22 (2020).

10. L. Stamatatos, L. Morris, D. R. Burton, J. R. Mascola, Neutralizing antibodies generated during natural hiv-1 infection: Good news for an hiv-1 vaccine? Nat. Med. 15, 866–870 (2009).

11. F. Klein, et al., Antibodies in HIV-1 vaccine development and therapy. Science (80-.349 cl:326). 341, 1199–1204 (2013).

12. E. Landais, P. L. Moore, Development of broadly neutralizing antibodies in HIV-1 infected elite neutralizers. Retrovirology 15, 1–14 (2018).

13. I. A. Abela, C. Kadelka, A. Trkola, Correlates of broadly neutralizing antibody development. Curr. Opin. HIV AIDS 14, 279–285 (2019).

14. H. Gruell, P. Schommers, Broadly neutralizing antibodies against HIV-1 and concepts for application. Curr. Opin. Virol. 54, 101211 (2022).

15. A. J. Hessell, et al., Broadly neutralizing human anti-HIV antibody 2G12 is effective in protection against mucosal SHIV challenge even at low serum neutralizing titers. PLoS Pathog. 5 (2009).

16. F. Klein, et al., HIV therapy by a combination of broadly neutralizing antibodies in humanized mice. Nature 492, 118–122 (2012).

17. B. Moldt, et al., Highly potent HIV-specific antibody neutralization in vitro translates into effective protection against mucosal SHIV challenge in vivo. Proc. Natl. Acad. Sci. U. S. A. 109, 18921–18925 (2012).

18. M. Shingai, et al., Passive transfer of modest titers of potent and broadly neutralizing anti-HIV monoclonal antibodies block SHIV infection in macaques. J. Exp. Med. 211, 2061–2074 (2014).

19. R. Gautam, et al., A single injection of anti-HIV-1 antibodies protects against repeated SHIV challenges. Nature 533, 105–109 (2016).

20. Y. Nishimura, et al., Early antibody therapy can induce long-lasting immunity to SHIV. Nature 543, 559–563 (2017).

21. M. Caskey, et al., Viraemia suppressed in HIV-1-infected humans by broadly neutralizing antibody 3BNC117. Nature 522, 487–491 (2015).

22. T. Schoofs, et al., HIV-1 therapy with monoclonal antibody 3BNC117 elicits host immune responses against HIV-1. Science (80-.). 352, 997–1001 (2016).

23. M. Caskey, et al., Antibody 10-1074 suppresses viremia in HIV-1-infected individuals. Nat. Med. 23, 185–191 (2017).

24. J. F. Scheid, et al., HIV-1 antibody 3BNC117 suppresses viral rebound in humans during treatment interruption. Nature 535, 556–560 (2016).

25. Y. Bar-On, et al., Safety and antiviral activity of combination HIV-1 broadly neutralizing antibodies in viremic individuals. Nat. Med. 24, 1701–1707 (2018).

26. P. Mendoza, et al., Combination therapy with anti-HIV-1 antibodies maintains viral suppression. Nature 561, 479–484 (2018).

27. C. Gaebler, et al., Prolonged viral suppression with anti-HIV-1 antibody therapy. Nature (2022) https://doi.org/10.1038/s41586-022-04597-1.

28. R. M. Lynch, et al., Virologic effects of broadly neutralizing antibody VRC01 administration during chronic HIV-1 infection. Sci. Transl. Med. 7, 1–15 (2015).

29. B. Julg, et al., Safety and antiviral activity of triple combination broadly neutralizing monoclonal antibody therapy against HIV-1: a phase 1 clinical trial. Nat. Med. 27, 1718–1724 (2022).

30. K. E. Stephenson, et al., Safety, pharmacokinetics and antiviral activity of PGT121, a broadly neutralizing monoclonal antibody against HIV-1: a randomized, placebo-controlled, phase 1 clinical trial. Nat. Med. 27, 1718–1724 (2021).

31. L. Corey, et al., Two Randomized Trials of Neutralizing Antibodies to Prevent HIV-1 Acquisition. N. Engl. J. Med. 384, 1003–1014 (2021).

32. J. Huang, et al., Identification of a CD4-Binding-Site Antibody to HIV that Evolved Near-Pan Neutralization Breadth. Immunity 45, 1108–1121 (2016).

33. M. M. Sajadi, et al., Identification of Near-Pan-neutralizing Antibodies against HIV-1 by Deconvolution of Plasma Humoral Responses. Cell 173, 1783–1795.e14 (2018).

34. M. Sun, et al., Rational design and characterization of the novel, broad and potent bispecific HIV-1 neutralizing antibody iMabm36. J. Acquir. Immune Defic. Syndr. 66, 473–483 (2014).

35. J. J. Steinhardt, et al., Rational design of a trispecific antibody targeting the HIV-1 Env with elevated anti-viral activity. Nat. Commun. 9, 1–12 (2018).

36. E. Rujas, et al., Engineering pan–HIV-1 neutralization potency through multispecific antibody avidity. Proc. Natl. Acad. Sci. 119, 1–11 (2022).

37. L. Xu, et al., Trispecific broadly neutralizing HIV antibodies mediate potent SHIV protection in macaques. Science (80-.). 358, 85–90 (2017).

38. L. M. Walker, et al., Broad neutralization coverage of HIV by multiple highly potent antibodies. Nature 477, 466–470 (2011).

39. T. B. Kepler, et al., Immunoglobulin gene insertions and deletions in the affinity maturation of HIV-1 broadly reactive neutralizing antibodies. Cell Host Microbe 16, 304–313 (2014).

40. H. B. Gristick, et al., Natively glycosylated HIV-1 Env structure reveals new mode for antibody recognition of the CD4-binding site. Nat. Struct. Mol. Biol. 23, 906–915 (2016).

41. T. Zhou, et al., Structural basis for broad and potent neutralization of HIV-1 by antibody VRC01. Science (80-.). 329, 811–817 (2010).

42. D. T. MacLeod, et al., Early Antibody Lineage Diversification and Independent Limb Maturation Lead to Broad HIV-1 Neutralization Targeting the Env High-Mannose Patch. Immunity 44, 1215–1226 (2016).

43. V. Pascual, et al., Nucleotide sequence analysis of the V regions of two IgM cold agglutinins: Evidence that the V(H)4-21 gene segment is responsible for the major cross-reactive idiotype. J. Immunol. 146, 4385–43891 (1991).

44. R. F. van Vollenhoven, et al., VH4-34 encoded antibodies in systemic lupus erythematosus: a specific diagnostic marker that correlates with clinical disease characteristics. J. Rheumatol. 26, 1727–1733 (1999).

45. H. Wardemann, et al., Predominant autoantibody production by early human B cell precursors. Science (80-.). 301, 1374–1377 (2003).

46. B. F. Haynes, et al., Immunology: Cardiolipin polyspecific autoreactivity in two broadly neutralizing HIV-1 antibodies. Science (80-.). 308, 1906–1908 (2005).

47. M. Liu, et al., Polyreactivity and Autoreactivity among HIV-1 Antibodies. J. Virol. 89, 784–798 (2015).

48. B. F. Haynes, G. Kelsoe, S. C. Harrison, T. B. Kepler, B-cell-lineage immunogen design in vaccine development with HIV-1 as a case study. Nat. Biotechnol. 30, 423–433 (2012).

49. F. A. Tucci, et al., Biased IGH VDJ gene repertoire and clonal expansions in B cells of chronically hepatitis C virus–infected individuals. Blood 131, 546–557 (2018).

50. H. X. Liao, et al., Co-evolution of a broadly neutralizing HIV-1 antibody and founder virus. Nature 496, 469–476 (2013).

51. B. Briney, et al., Tailored Immunogens Direct Affinity Maturation toward HIV Neutralizing Antibodies. Cell 166, 1459–1470.e11 (2016).

52. J. M. Steichen, et al., A generalized HIV vaccine design strategy for priming of broadly neutralizing antibody responses. Science (80-.). 366 (2019).

53. C. Yacoob, et al., Differences in Allelic Frequency and CDRH3 Region Limit the Engagement of HIV Env Immunogens by Putative VRC01 Neutralizing Antibody Precursors. Cell Rep. 17, 1560–1570 (2016).

54. C. Havenar-Daughton, et al., The human naive B cell repertoire contains distinct subclasses for a germline-targeting HIV-1 vaccine immunogen. Sci. Transl. Med. 10, 1–16 (2018).

55. K. Wiehe, et al., Functional Relevance of Improbable Antibody Mutations for HIV Broadly Neutralizing Antibody Development. Cell Host Microbe 23, 759–765.e6 (2018).

56. S. D. Boyd, et al., Individual Variation in the Germline Ig Gene Repertoire Inferred from Variable Region Gene Rearrangements. J. Immunol. 184, 6986–6992 (2010).

57. L. D. Goldstein, et al., Massively parallel single-cell B-cell receptor sequencing enables rapid discovery of diverse antigen-reactive antibodies. Commun. Biol. 2, 1–10 (2019).

58. B. Briney, A. Inderbitzin, C. Joyce, D. R. Burton, Commonality despite exceptional diversity in the baseline human antibody repertoire. Nature 566, 393–397 (2019).

59. H. Yoon, et al., CATNAP: A tool to compile, analyze and tally neutralizing antibody panels. Nucleic Acids Res. 43, W213–W219 (2015).

60. D. Sok, D. R. Burton, Recent progress in broadly neutralizing antibodies to HIV. Nat. Immunol. 19, 1179–1188 (2018).

61. Y. Elhanati, et al., Inferring processes underlying B-cell repertoire diversity. Philos. Trans. R. Soc. B Biol. Sci. 370 (2015).

62. Q. Marcou, T. Mora, A. M. Walczak, High-throughput immune repertoire analysis with IGoR. Nat. Commun. 9 (2018).

63. L. Cooper, K. L. Good-Jacobson, Dysregulation of humoral immunity in chronic infection. Immunol. Cell Biol. 98, 456–466 (2020).

64. A. deCamp, et al., Global Panel of HIV-1 Env Reference Strains for Standardized Assessments of Vaccine-Elicited Neutralizing Antibodies. J. Virol. 88, 2489–2507 (2014).

65. D. H. Barouch, et al., Therapeutic efficacy of potent neutralizing HIV-1-specific monoclonal antibodies in SHIV-infected rhesus monkeys. Nature 503, 224–228 (2013).

66. L. M. Walker, D. R. Burton, Passive immunotherapy of viral infections: “super-antibodies” enter the fray. Nat. Rev. Immunol. 18, 297–308 (2018).

67. H. Gruell, F. Klein, Antibody-mediated prevention and treatment of HIV-1 infection. Retrovirology 15, 1–11 (2018).

68. P. D. Kwong, J. R. Mascola, HIV-1 Vaccines Based on Antibody Identification, B Cell Ontogeny, and Epitope Structure. Immunity 48, 855–871 (2018).

69. K. O. Saunders, et al., Targeted selection of HIV-specific antibody mutations by engineering B cell maturation. Science (80-.). 366 (2019).

70. F. Klein, et al., Somatic mutations of the immunoglobulin framework are generally required for broad and potent HIV-1 neutralization. Cell 153, 126–138 (2013).

71. J. P. Julien, et al., Asymmetric recognition of the HIV-1 trimer by broadly neutralizing antibody PG9. Proc. Natl. Acad. Sci. U. S. A. 110, 4351–4356 (2013).

72. J. S. McLellan, et al., Structure of HIV-1 gp120 V1/V2 domain with broadly neutralizing antibody PG9. Nature 480, 336–343 (2011).

73. R. Pejchal, et al., Structure and function of broadly reactive antibody PG16 reveal an H3 subdomain that mediates potent neutralization of HIV-1. Proc. Natl. Acad. Sci. U. S. A. 107, 11483–11488 (2010).

74. M. Bonsignori, et al., Staged induction of HIV-1 glycan-dependent broadly neutralizing antibodies. Sci. Transl. Med. 9 (2017).

75. D. Cizmeci, et al., Distinct clonal evolution of b-cells in hiv controllers with neutralizing antibody breadth. Elife 10, 1–15 (2021).

76. N. A. Doria-Rose, et al., Developmental pathway for potent V1V2-directed HIV-neutralizing antibodies. Nature 508, 55–62 (2014).

77. P. L. Moore, et al., Potent and Broad Neutralization of HIV-1 Subtype C by Plasma Antibodies Targeting a Quaternary Epitope Including Residues in the V2 Loop. J. Virol. 85, 3128–3141 (2011).

78. C. K. Wibmer, et al., Viral Escape from HIV-1 Neutralizing Antibodies Drives Increased Plasma Neutralization Breadth through Sequential Recognition of Multiple Epitopes and Immunotypes. PLoS Pathog. 9 (2013).

79. K. M. Roskin, et al., Aberrant B cell repertoire selection associated with HIV neutralizing antibody breadth. Nat. Immunol. 21, 199–209 (2020).

80. M. Sarzotti-Kelsoe, et al., Optimization and validation of the TZM-bl assay for standardized assessments of neutralizing antibodies against HIV-1. J. Immunol. Methods 409, 131–146 (2014).

81. J. Ye, N. Ma, T. L. Madden, J. M. Ostell, IgBLAST: an immunoglobulin variable domain sequence analysis tool. Nucleic Acids Res. (2013) https://doi.org/10.1093/nar/gkt382.

82. M. Goujon, et al., A new bioinformatics analysis tools framework at EMBL-EBI. Nucleic Acids Res. 38, 695–699 (2010).

83. F. Sievers, et al., Fast, scalable generation of high-quality protein multiple sequence alignments using Clustal Omega. Mol. Syst. Biol. (2011) https://doi.org/10.1038/msb.2011.75.

84. J. A. Vander Heiden, et al., PRESTO: A toolkit for processing high-throughput sequencing raw reads of lymphocyte receptor repertoires. Bioinformatics 30, 1930–1932 (2014).

85. M.-P. Lefranc, et al., IMGT, the international ImMunoGeneTics database. Nucleic Acids Res. 27, 209–212 (1999).

86. M.-P. Lefranc, From IMGT-ONTOLOGY IDENTIFICATION axiom to IMGT standardized keywords: for immunoglobulins (IG), T cell receptors (TR), and conventional genes. Cold Spring Harb. Protoc. 2011, 604–613 (2011).

87. M. S. Seaman, et al., Tiered Categorization of a Diverse Panel of HIV-1 Env Pseudoviruses for Assessment of Neutralizing Antibodies. J. Virol. 84, 1439–1452 (2010).

88. S. R. Walsh, M. S. Seaman, Broadly Neutralizing Antibodies for HIV-1 Prevention. Front. Immunol. 12, 1–14 (2021).

89. W. S. DeWitt, et al., A public database of memory and naive B-cell receptor sequences. PLoS One 11, 1–18 (2016).

90. A. Stamatakis, RAxML version 8: A tool for phylogenetic analysis and post-analysis of large phylogenies. Bioinformatics 30, 1312–1313 (2014).

91. A. Nourmohammad, J. Otwinowski, M. Łuksza, T. Mora, A. M. Walczak, Fierce Selection and Interference in B-Cell Repertoire Response to Chronic HIV-1. Mol. Biol. Evol. 36, 2184–2194 (2019).

