## Supplementary material for "Probabilities of HIV-1 bNAb development in healthy and chronically infected individuals": SI_Kreer_Lupo_Ercanoglu_et_al

### Supplementary Figures


| | |
|---|---|
| Supplementary Figure 1 | Gating strategy to isolate naive or antigen-experienced B cells |
| Supplementary Figure 2 | Unique molecular identifier copy number distribution of fully annotated reads |
| Supplementary Figure 3 | Spike-in experiment with different percentages of known cell line clones |
| Supplementary Figure 4 | Influence of viral panel composition on the determination of neutralization breadth and potency |
| Supplementary Figure 5 | Correlations of 75 bNAb heavy chain sequence features and probability values |
| Supplementary Figure 6 | Light chain probabilities of broadly neutralizing antibodies |
| Supplementary Figure 7 | IgG heavy and light chain repertoire characteristics, stratified by antiretroviral treatment |
| Supplementary Figure 8 | IgG heavy and light chain repertoire characteristics, stratified by neutralization activity |


### Supplementary Tables


| | |
|---|---|
| Supplementary Table 1 | Control experiment processing and sequencing statistics |
| Supplementary Table 2 | Healthy cohort demographics and and sequencing statistics |
| Supplementary Table 3 | Viral panel overview |
| Supplementary Table 4 | Features of HIV-1 broadly neutralizing antibodies |
| Supplementary Table 5 | Heavy and light chain Pgen, Pshm, and probability scores |
| Supplementary Table 6 | HIV-1 cohort demographics and and sequencing statistics |
| Supplementary Table 7 | HCV cohort demographics and and sequencing statistics |
| Supplementary Table 8 | HIV-1 cohort poly IgG neutralization on the 12-strain global panel |

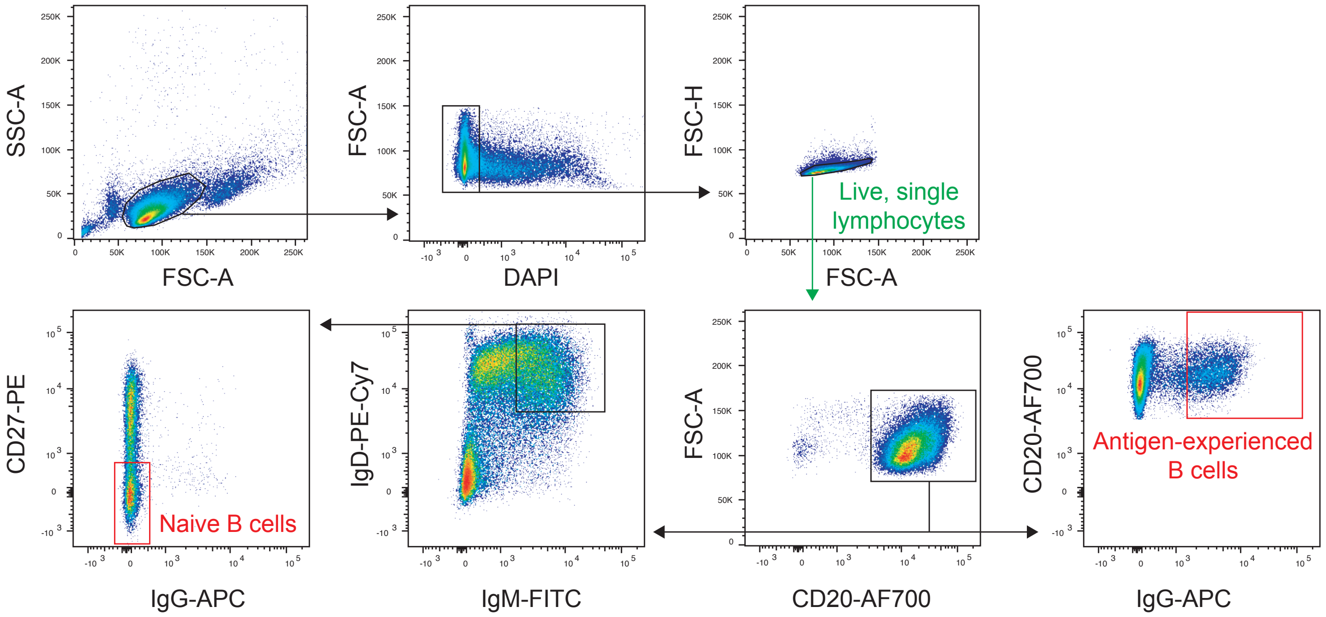


**Supplementary Figure 1: Gating strategy to isolate naive or antigen-experienced B cells.** Gating includes the pre-selection of the lymphocyte population, exclusion of dead cells and doublets, as well as selection of CD20$^{+}$ cells. From this population, antigen-experienced are defined as IgG$^{+}$ and naive B cells as IgD$^{+}$/IgM$^{+}$ and CD27$^{-}$/IgG$^{-}$.

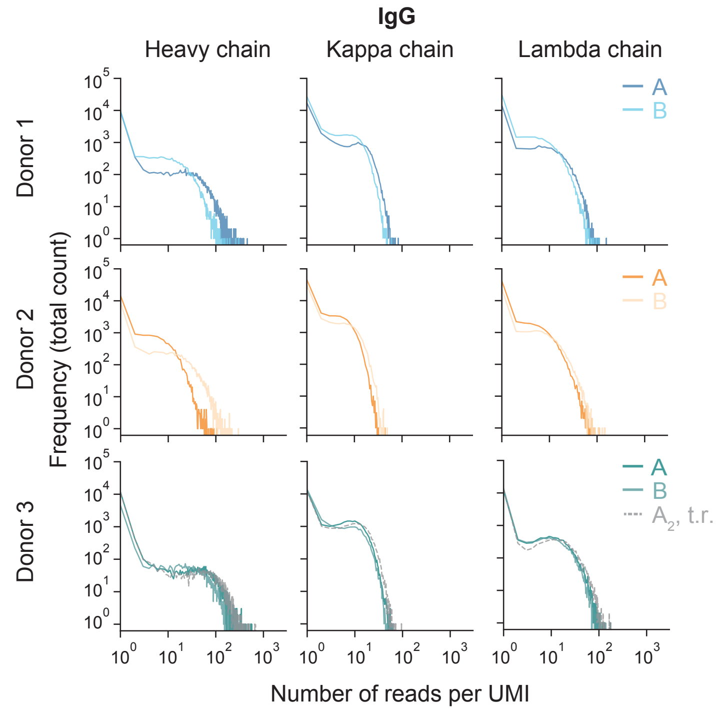


**Supplementary Figure 2: Unique molecular identifier copy number distributions of fully annotated reads.** Numbers of fully annotated reads per UMI were counted (x-axis) and the frequency of UMI groups with identical count numbers determined (y-axis). Colored solid lines represent biological replicates, dashed lines represent the technical replicate (t.r., donor 3, sample $A_2$). The technical replicate was processed independently from all other samples on a different day and sequenced in a separate NGS run.

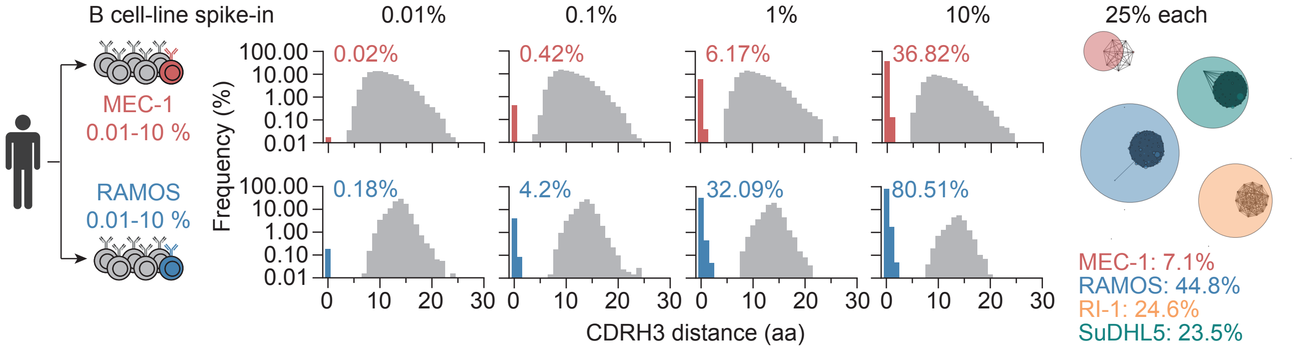


**Supplementary Figure 3: Spike-in experiment with different percentages of known cell line clones.** IgM B cells from a healthy donor were spiked with different concentrations (0.1 - 10%) of tumor B cell lines (MEC-1 and RAMOS) to analyze in total 100,000 cells. Plots show amino acid (aa) distances of all reconstituted CDRH3s from the individual spike-in experiments in comparison to the cell line CDRH3. Colored bars depict CDRH3s with <4 amino acids difference to the cell line CDRH3 and colored percentages represent their fraction of all comparisons. As an internal control, a sample comprising 25,000 cells from each of the four depicted cell lines was processed for repertoire sequencing. Each node represents a unique CDRH3. The node size is proportional to the frequency among all identified CDRH3s. Nodes are connected if they share at least 75% of their CDRH3 amino acid sequence. Nodes are colored according to the cell lines if they share at least 75% of the CDRH3 amino acid sequence with a cell line.

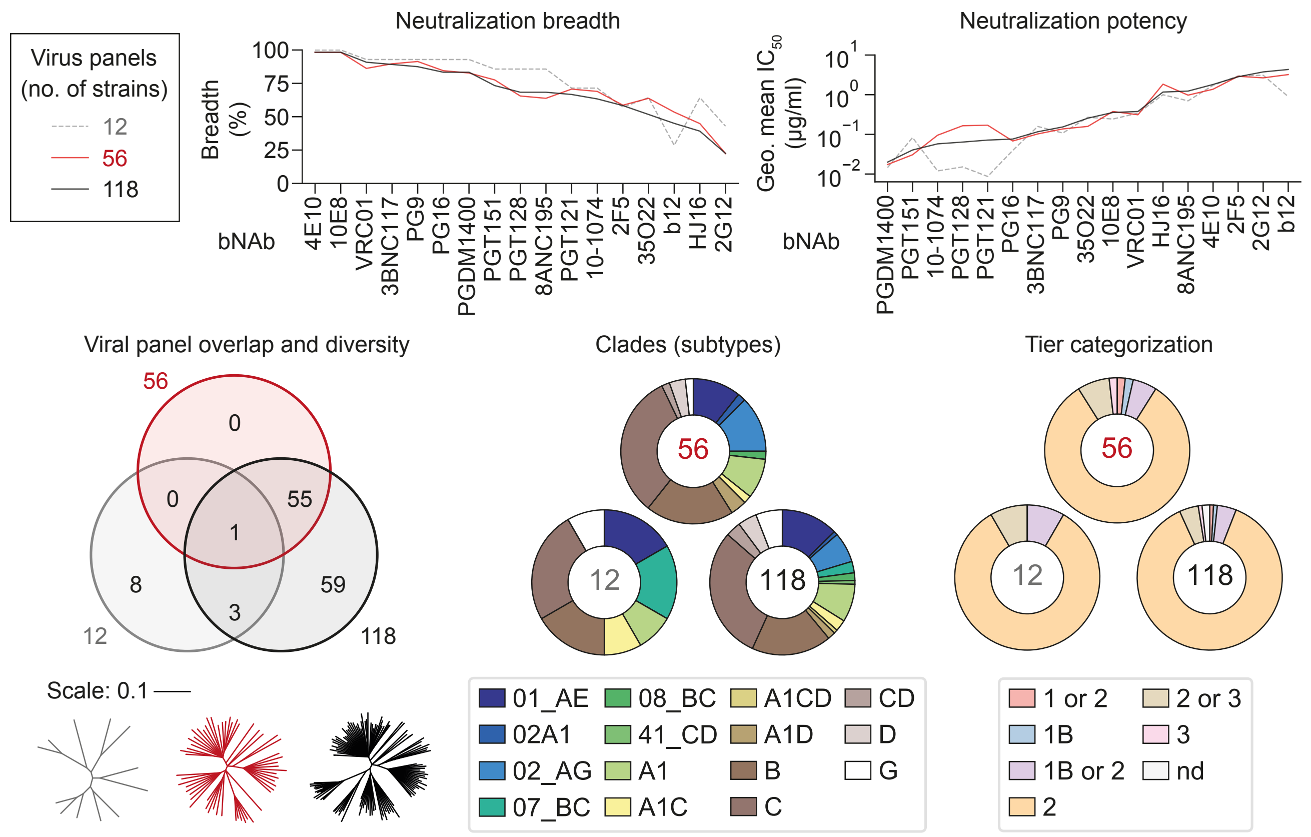


**Supplementary Figure 4: Influence of viral panel composition on the determination of neutralization breadth and potency.** Neutralization breadth and potency (geometric mean IC50 of neutralized strains) for 17 bNAbs based on three different viral panels: the 12 strain "global" panel (deCamp et al., 2014), the 118-strain multi-clade panel (Seaman et al., 2010), and a 56-strain subset panel, which was selected for the analyses in this work. The data for neutralization breadth and potency (upper part) is sorted by the values from the 118-strain panel. The lower part shows a comparison of the three panels in terms of overlap, phylogenetic trees, as well as clade composition and Tier categorization (lower part).

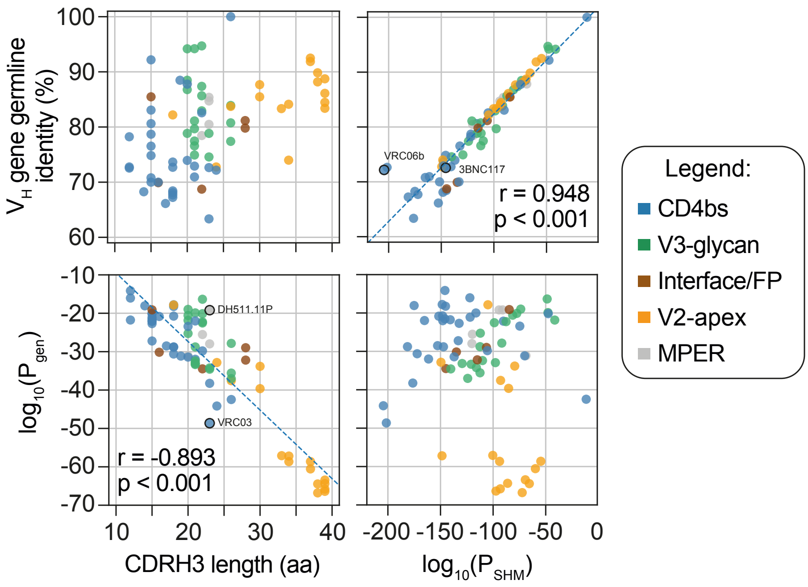


**Supplementary Figure 5: Correlations of 75 bNAb heavy chain sequence features and probability values.** Correlation coefficients (r) and p values were determined by linear regression (dashed line). Colors encode for binding sites. Labeled antibodies are highlighted by black outlines.

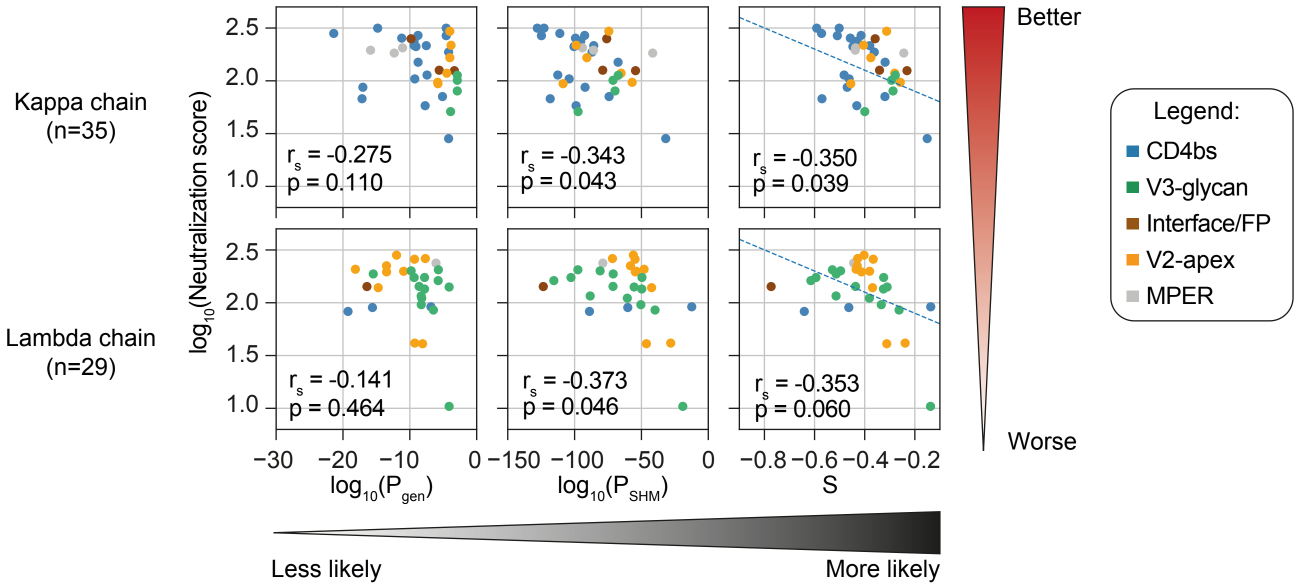


**Supplementary Figure 6: Light chain probabilities of broadly neutralizing antibodies.** Correlation plots of bNAb neutralization scores against light chain $P_{gen}$, $P_{SHM}$ and probability scores $S = c_1 log_{10}(P_{gen}) + c_2 log_{10}(P_{SHM})$, separated by light chain isotype (n=35 kappa chains, n=29 lambda chains). Probability scores were derived by a linear regression (dashed line) with $c_1$ = $6.869x10^{-03}$ and $c_2$ = $3.829x10^{-03}$ for kappa and $c_1$ = $1.134x10^{-02}$ and $c_2$ = $4.756x10^{-03}$ for lambda chains. Spearman correlation coefficients $r_s$ and p values are given in the figure. Correlation coefficients and p values from linear regressions for S are r = -0.403 and p = 0.016 for kappa and r = -0.470 and p = 0.010 for lambda chains, respectively.

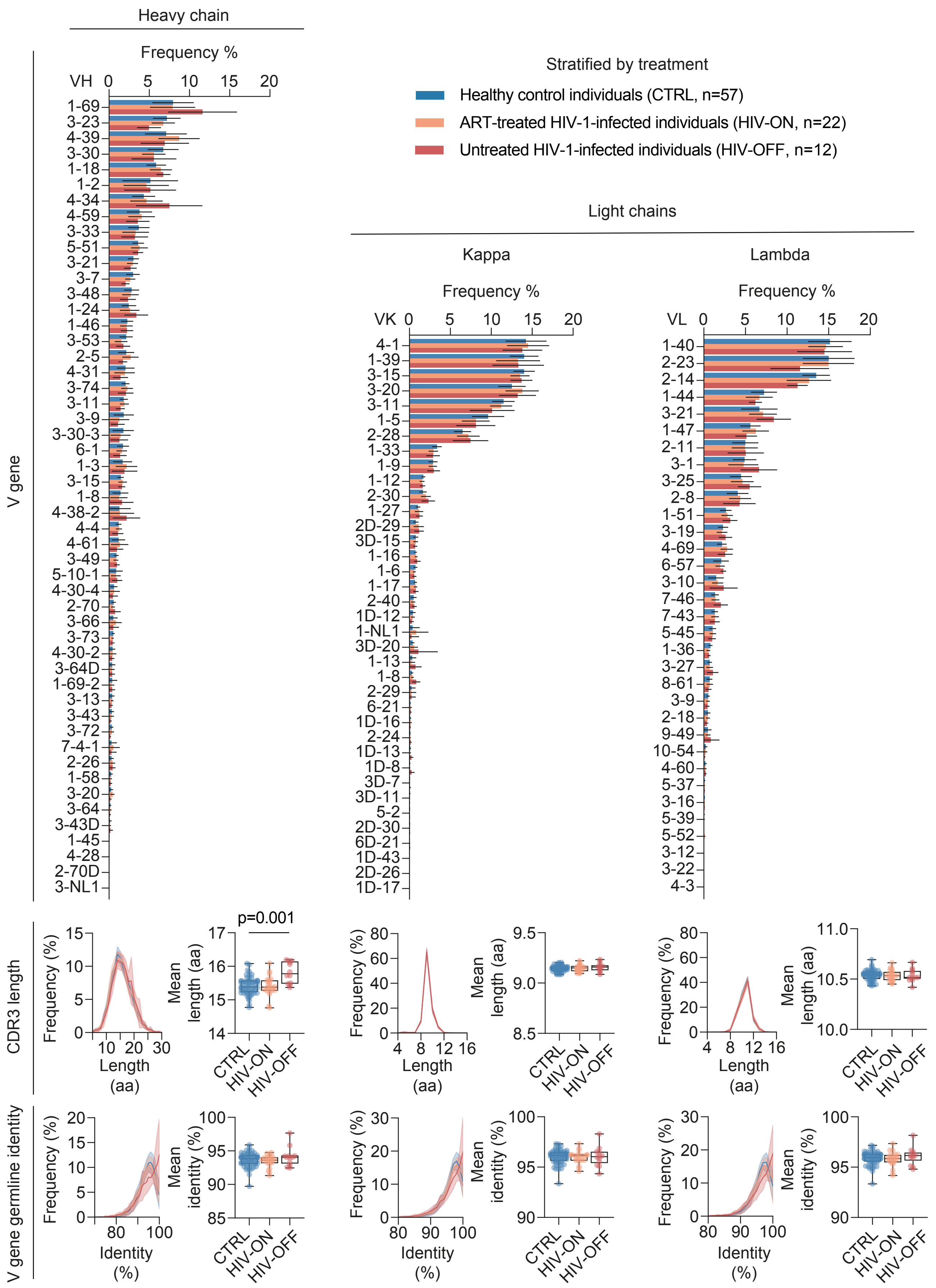


**Supplementary Figure 7: IgG heavy chain and light chain repertoire characteristics, stratified by antiretroviral treatment.** Heavy and light chain V gene, CDR3 length, and V gene germline nucleotide identity distributions for healthy (CTRL, n=57), as well as treated (HIV-ON, n=22) and untreated (HIV-OFF, n=12) HIV-1-infected individuals. Differences in mean CDR3 lengths were determined by one-way ANOVA and Tukey post hoc test. aa: amino acids.

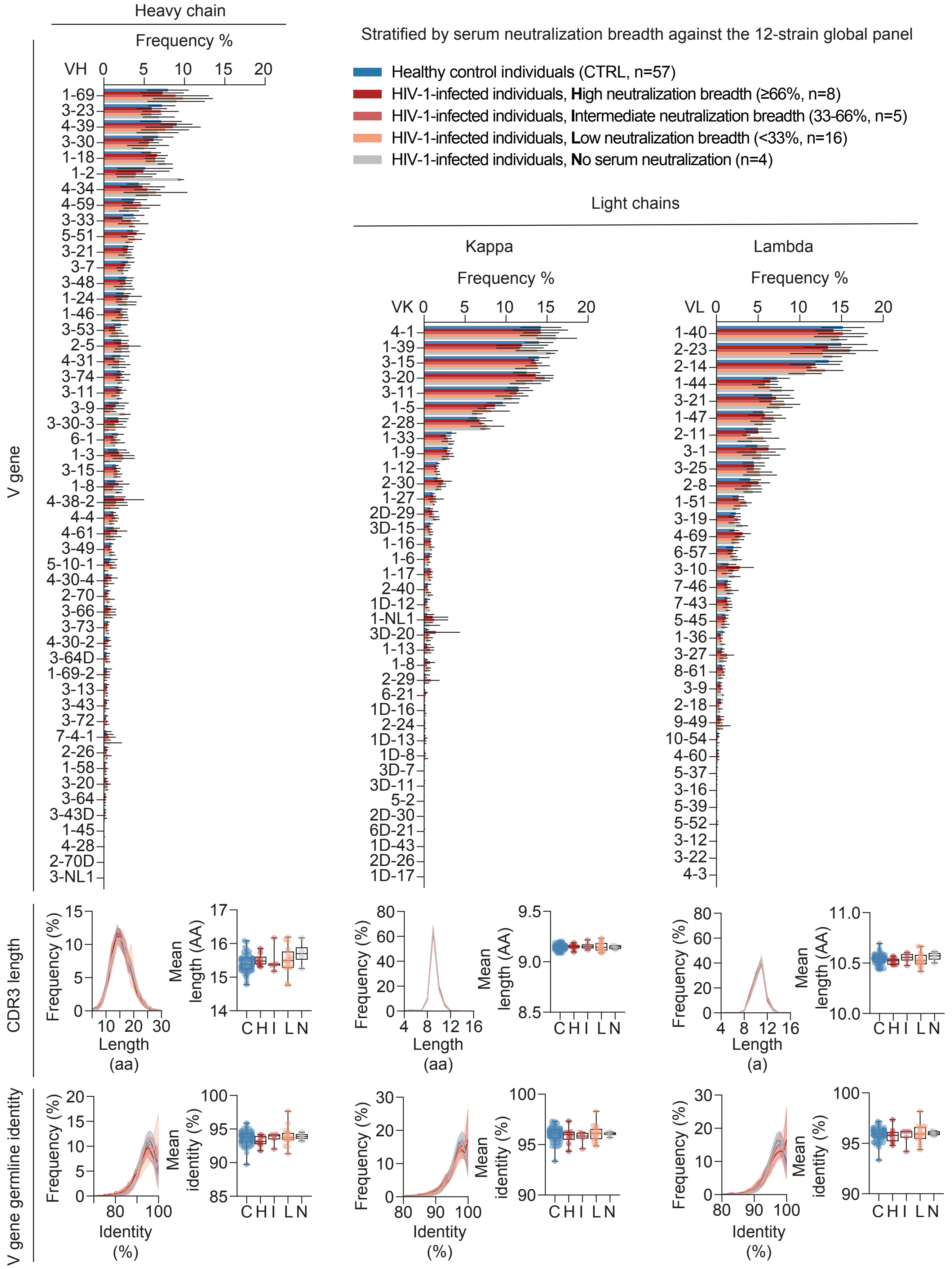


**Supplementary Figure 8: IgG heavy and light chain repertoire characteristics, stratified by neutralization activity.** Heavy and light chain V gene, CDR3 length, and V gene germline identity distributions for healthy (CTRL, n=57), as well as HIV-1-infected individuals (n=34) who have been grouped by their serum neutralization breadth against the 12-strain global panel into high (H, n=8), intermediate (I, n=5), low (L, n=16), and no neutralization breadth (N, n=4).

**Table S1: Control experiment processing and sequencing statistics**

| Donor | Set | Biological replicate | Cell number | Technical replicate | Raw reads | Assembled and V(D)J annotated reads | | | | Full V(D)J alignment | | | | Median number of reads per UMI | | |
|---|---|---|---|---|---|---|---|---|---|---|---|---|---|---|---|---|
| | | | | | Total | H | K | L | Total | H | K | L | Total | H | K | L |
| 1 | 1 | A | 100.000 | 1 | 1.630.514 | 16.156 | 37.585 | 28.781 | 82.522 | 16.156 | 37.467 | 28.781 | 82.404 | 9,5 | 96 | 25 |
| | 2 | B | 100.000 | 1 | 2.533.070 | 17.816 | 53.449 | 50.092 | 121.357 | 17.816 | 53.228 | 50.092 | 121.136 | 7 | 104 | 23 |
| 2 | 3 | A | 100.000 | 1 | 2.136.806 | 23.208 | 73.083 | 60.599 | 156.890 | 23.208 | 72.833 | 60.599 | 156.640 | 7 | 123,5 | 18 |
| | 4 | B | 100.000 | 1 | 2.552.903 | 15.824 | 53.782 | 38.905 | 108.511 | 15.824 | 53.566 | 38.905 | 108.295 | 10 | 92 | 20 |
| 3 | 5 | A | 100.000 | 1 | 1.354.097 | 17.272 | 34.760 | 22.254 | 74.286 | 17.272 | 34.709 | 22.254 | 74.235 | 10 | 109 | 27 |
| | 6 | | | 2 | 1.695.595 | 16.739 | 34.723 | 23.919 | 75.381 | 16.739 | 34.664 | 23.919 | 75.322 | 10 | 110,5 | 19 |
| | 7 | B | 100.000 | 1 | 1.476.766 | 8.945 | 30.116 | 22.628 | 61.689 | 8.945 | 29.976 | 22.628 | 61.549 | 8 | 97 | 21,5 |
| | Minimum | | | | 1.354.097 | 8.945 | 30.116 | 22.254 | 61.689 | 8.945 | 29.976 | 22.254 | 61.549 | 7 | 92 | 18 |
| | Maximum | | | | 2.552.903 | 23.208 | 73.083 | 60.599 | 156.890 | 23.208 | 72.833 | 60.599 | 156.640 | 10 | 124 | 27 |
| | Mean | | | | 1.911.393 | 16.566 | 45.357 | 35.311 | 97.234 | 16.566 | 45.206 | 35.311 | 97.083 | 9 | 105 | 22 |
| | Median | | | | 1.695.595 | 16.739 | 37.585 | 28.781 | 82.522 | 16.739 | 37.467 | 28.781 | 82.404 | 10 | 104 | 22 |
| | Std. deviation | | | | 458.845 | 3.873 | 14.259 | 14.013 | 31.058 | 3.873 | 14.196 | 14.013 | 30.995 | 1 | 10 | 3 |
| | Sum | | | | 13.379.751 | 115.960 | 317.498 | 247.178 | 680.636 | 115.960 | 316.443 | 247.178 | 679.581 | - | - | - |

**Table S1: Control experiment processing and sequencing statistics - continued**

| Donor | Set | Biological replicate | Cell number | Technical replicate | Raw reads | Sequences with ≥3 reads per UMI | | | | Productive sequences | | | | Unique CDR3s | | | |
|---|---|---|---|---|---|---|---|---|---|---|---|---|---|---|---|---|---|
| | | | | | Total | H | K | L | Total | H | K | L | Total | H | K | L | Total |
| 1 | 1 | A | 100.000 | 1 | 1.630.514 | 6.829 | 18.220 | 13.647 | 38.696 | 6.581 | 17.083 | 12.426 | 36.090 | 5.616 | 6.561 | 5.271 | 17.448 |
| | 2 | B | 100.000 | 1 | 2.533.070 | 8.054 | 23.020 | 17.085 | 48.159 | 7.779 | 21.620 | 15.570 | 44.969 | 6.530 | 7.606 | 5.833 | 19.969 |
| 2 | 3 | A | 100.000 | 1 | 2.136.806 | 8.089 | 22.785 | 16.707 | 47.581 | 7.865 | 21.493 | 15.481 | 44.839 | 6.705 | 8.763 | 6.941 | 22.409 |
| | 4 | B | 100.000 | 1 | 2.552.903 | 7.105 | 20.289 | 14.647 | 42.041 | 6.861 | 19.185 | 13.626 | 39.672 | 5.906 | 8.234 | 6.498 | 20.638 |
| 3 | 5 | A | 100.000 | 1 | 1.354.097 | 5.074 | 21.241 | 10.255 | 36.570 | 4.860 | 19.421 | 9.398 | 33.679 | 4.279 | 8.693 | 4.941 | 17.913 |
| | 6 | | | 2 | 1.695.595 | 5.174 | 21.849 | 10.442 | 37.465 | 4.927 | 19.988 | 9.565 | 34.480 | 4.329 | 8.818 | 4.994 | 18.141 |
| | 7 | B | 100.000 | 1 | 1.476.766 | 4.133 | 14.727 | 8.752 | 27.612 | 3.974 | 13.283 | 7.936 | 25.193 | 3.623 | 6.735 | 4.394 | 14.752 |
| | Minimum | | | | 1.354.097 | 4.133 | 14.727 | 8.752 | 27.612 | 3.974 | 13.283 | 7.936 | 25.193 | 3.623 | 6.561 | 4.394 | 14.752 |
| | Maximum | | | | 2.552.903 | 8.089 | 23.020 | 17.085 | 48.159 | 7.865 | 21.620 | 15.570 | 44.969 | 6.705 | 8.818 | 6.941 | 22.409 |
| | Mean | | | | 1.911.393 | 6.351 | 20.304 | 13.076 | 39.732 | 6.121 | 18.868 | 12.000 | 36.989 | 5.284 | 7.916 | 5.553 | 18.753 |
| | Median | | | | 1.695.595 | 6.829 | 21.241 | 13.647 | 38.696 | 6.581 | 19.421 | 12.426 | 36.090 | 5.616 | 8.234 | 5.271 | 18.141 |
| | Std. deviation | | | | 458.845 | 1.447 | 2.732 | 3.062 | 6.558 | 1.423 | 2.686 | 2.851 | 6.433 | 1.118 | 891 | 846 | 2.305 |
| | Sum | | | | 13.379.751 | 44.458 | 142.131 | 91.535 | 278.124 | 42.847 | 132.073 | 84.002 | 258.922 | 36.988 | 55.410 | 38.872 | 131.270 |

**Table S2: Healthy cohort demographics and and sequencing statistics**

| # | Internal ID | Sex | Age | Cell Number | Raw reads | Assembled and V(D)J annotated reads | | | | Full V(D)J alignment | | | | Median number of reads per UMI | | |
|---|---|---|---|---|---|---|---|---|---|---|---|---|---|---|---|---|
| | | | | | Total | H | K | L | Total | H | K | L | Total | H | K | L |
| **1** | 75 | **f** | 47 | 100.000 | 1.470.930 | 14.835 | 24.555 | 15.382 | 54.772 | 14.835 | 24.468 | 15.382 | 54.685 | 5 | 114 | 28 |
| **2** | 76 | **f** | 51 | 100.000 | 1.749.311 | 15.072 | 21.954 | 17.035 | 54.061 | 15.072 | 21.884 | 17.035 | 53.991 | 6 | 99 | 20 |
| **3** | 84 | **m** | 47 | 100.000 | 1.655.132 | 28.255 | 18.368 | 12.212 | 58.835 | 28.255 | 18.298 | 12.212 | 58.765 | 9 | 60 | 14 |
| **4** | 85 | **m** | 23 | 100.000 | 1.710.794 | 19.455 | 18.727 | 9.859 | 48.041 | 19.455 | 18.647 | 9.859 | 47.961 | 4 | 54,5 | 13 |
| **5** | 86 | **m** | 47 | 100.000 | 1.472.918 | 12.588 | 11.399 | 5.374 | 29.361 | 12.588 | 11.333 | 5.374 | 29.295 | 2 | 24,5 | 6 |
| **6** | 90 | **f** | 25 | 100.000 | 1.728.880 | 65.143 | 32.552 | 24.898 | 122.593 | 65.143 | 32.447 | 24.898 | 122.488 | 10 | 122 | 22,5 |
| **7** | 91 | **m** | 55 | 100.000 | 1.710.077 | 85.034 | 42.767 | 15.249 | 143.050 | 85.034 | 42.663 | 15.249 | 142.946 | 13 | 157 | 20 |
| **8** | 92 | **f** | 54 | 100.000 | 1.694.506 | 80.483 | 34.285 | 22.624 | 137.392 | 80.483 | 34.174 | 22.624 | 137.281 | 9 | 121,5 | 27 |
| **9** | 93 | **m** | 34 | 100.000 | 2.004.033 | 86.172 | 34.020 | 23.449 | 143.641 | 86.172 | 33.922 | 23.449 | 143.543 | 12 | 116 | 27 |
| **10** | 94 | **m** | 31 | 100.000 | 2.045.940 | 85.687 | 44.386 | 18.641 | 148.714 | 85.687 | 44.269 | 18.641 | 148.597 | 8 | 134,5 | 13 |
| **11** | 95 | **m** | 19 | 100.000 | 1.739.870 | 136.867 | 36.639 | 27.464 | 200.970 | 136.867 | 36.573 | 27.464 | 200.904 | 22 | 92 | 27 |
| **12** | 96 | **m** | 55 | 100.000 | 1.965.848 | 22.551 | 31.927 | 20.348 | 74.826 | 22.551 | 31.801 | 20.348 | 74.700 | 6 | 47,5 | 13 |
| **13** | 97 | **f** | 22 | 100.000 | 1.677.943 | 33.863 | 25.917 | 19.348 | 79.128 | 33.863 | 25.769 | 19.348 | 78.980 | 7 | 57,5 | 22 |
| **14** | 98 | **m** | 34 | 100.000 | 1.672.543 | 22.499 | 20.346 | 29.674 | 72.519 | 22.499 | 20.231 | 29.674 | 72.404 | 8 | 20 | 23,5 |
| **15** | 99 | **m** | 51 | 100.000 | 1.746.925 | 21.001 | 32.530 | 19.843 | 73.374 | 21.001 | 32.428 | 19.843 | 73.272 | 9 | 96 | 16 |
| **16** | 100 | **m** | 46 | 100.000 | 1.953.231 | 27.899 | 34.030 | 34.429 | 96.358 | 27.899 | 33.942 | 34.429 | 96.270 | 9 | 56,5 | 23,5 |
| **17** | 101 | **m** | 46 | 100.000 | 1.895.250 | 22.644 | 29.372 | 18.289 | 70.305 | 22.644 | 29.266 | 18.289 | 70.199 | 6 | 86,5 | 14 |
| **18** | 102 | **m** | 31 | 100.000 | 2.030.669 | 58.679 | 16.164 | 15.344 | 90.187 | 58.679 | 16.112 | 15.344 | 90.135 | 9 | 29 | 16 |
| **19** | 103 | **m** | 25 | 100.000 | 1.692.152 | 57.343 | 34.949 | 30.533 | 122.825 | 57.343 | 34.836 | 30.533 | 122.712 | 15 | 93 | 46 |
| **20** | 104 | **m** | 21 | 100.000 | 1.601.433 | 18.396 | 19.277 | 18.681 | 56.354 | 18.396 | 19.246 | 18.681 | 56.323 | 7 | 74 | 15 |
| **21** | 105 | **m** | 20 | 100.000 | 1.321.306 | 25.608 | 22.611 | 23.285 | 71.504 | 25.608 | 22.547 | 23.285 | 71.440 | 8 | 40,5 | 26 |
| **22** | 106 | **f** | 48 | 100.000 | 1.685.813 | 30.697 | 34.825 | 44.435 | 109.957 | 30.697 | 34.729 | 44.435 | 109.861 | 8 | 72 | 33,5 |
| **23** | 117 | **f** | 24 | 100.000 | 1.483.778 | 27.524 | 26.689 | 18.480 | 72.693 | 27.524 | 26.575 | 18.480 | 72.579 | 10 | 64 | 22 |
| **24** | 118 | **m** | 30 | 100.000 | 1.632.990 | 26.998 | 30.432 | 21.781 | 79.211 | 26.998 | 30.358 | 21.781 | 79.137 | 8 | 60 | 18,5 |
| **25** | 119 | **f** | 28 | 100.000 | 1.625.973 | 70.372 | 40.516 | 28.243 | 139.131 | 70.372 | 40.418 | 28.243 | 139.033 | 10 | 132,5 | 38 |
| **26** | 120 | **m** | 21 | 100.000 | 1.749.428 | 76.782 | 32.330 | 21.125 | 130.237 | 76.782 | 32.250 | 21.125 | 130.157 | 13 | 64 | 36 |
| **27** | 121 | **f** | 21 | 100.000 | 1.576.639 | 33.368 | 37.170 | 16.154 | 86.692 | 33.368 | 37.072 | 16.154 | 86.594 | 11 | 103 | 12 |
| **28** | 126 | **f** | 24 | 100.000 | 1.932.765 | 129.234 | 37.960 | 35.871 | 203.065 | 129.234 | 37.825 | 35.871 | 202.930 | 14 | 107,5 | 28,5 |
| **29** | 127 | **m** | 42 | 100.000 | 1.661.332 | 35.794 | 18.782 | 13.216 | 67.792 | 35.794 | 18.698 | 13.216 | 67.708 | 7 | 31 | 15 |
| **30** | 128 | **f** | 67 | 100.000 | 2.026.801 | 16.556 | 25.980 | 15.505 | 58.041 | 16.556 | 25.844 | 15.505 | 57.905 | 5 | 42 | 10,5 |
| **31** | 130 | **m** | 37 | 100.000 | 1.769.184 | 12.976 | 26.633 | 12.391 | 52.000 | 12.976 | 26.523 | 12.391 | 51.890 | 6 | 82,5 | 9,5 |
| **32** | 131 | **f** | 67 | 100.000 | 1.544.258 | 47.392 | 22.646 | 19.924 | 89.962 | 47.392 | 22.588 | 19.924 | 89.904 | 21 | 64 | 24,5 |
| **33** | 132 | **m** | 39 | 100.000 | 1.863.306 | 50.688 | 36.775 | 26.900 | 114.363 | 50.688 | 36.630 | 26.900 | 114.218 | 10 | 122 | 32 |
| **34** | 133 | **m** | 35 | 100.000 | 1.478.049 | 86.334 | 39.367 | 17.942 | 143.643 | 86.334 | 39.263 | 17.942 | 143.539 | 18 | 120 | 19 |
| **35** | 134 | **m** | 36 | 100.000 | 1.566.603 | 70.086 | 55.822 | 28.134 | 154.042 | 70.086 | 55.674 | 28.134 | 153.894 | 17,5 | 58 | 33 |
| **36** | 135 | **f** | 22 | 100.000 | 1.372.889 | 45.457 | 25.575 | 14.964 | 85.996 | 45.457 | 25.454 | 14.964 | 85.875 | 15 | 56,5 | 20 |
| **37** | 136 | **f** | 20 | 100.000 | 1.657.997 | 52.815 | 71.130 | 29.485 | 153.430 | 52.815 | 70.970 | 29.485 | 153.270 | 15 | 62,5 | 15 |
| **38** | 137 | **f** | 21 | 100.000 | 1.436.791 | 28.447 | 43.420 | 58.015 | 129.882 | 28.447 | 43.352 | 58.015 | 129.814 | 14 | 57 | 37,5 |
| **39** | 138 | **f** | 26 | 100.000 | 1.465.807 | 96.795 | 30.010 | 50.530 | 177.335 | 96.795 | 29.925 | 50.530 | 177.250 | 14 | 100,5 | 22 |
| **40** | 139 | **f** | 36 | 100.000 | 1.746.660 | 66.221 | 36.302 | 30.517 | 133.040 | 66.221 | 36.142 | 30.517 | 132.880 | 11 | 129 | 22,5 |
| **41** | 140 | **m** | 23 | 100.000 | 1.993.175 | 78.771 | 30.236 | 17.109 | 126.116 | 78.771 | 30.140 | 17.109 | 126.020 | 10 | 113,5 | 15 |
| **42** | 141 | **m** | 21 | 100.000 | 1.552.022 | 39.184 | 29.321 | 13.192 | 81.697 | 39.184 | 29.218 | 13.192 | 81.594 | 19 | 87 | 15 |
| **43** | 145 | **f** | 21 | 100.000 | 1.943.870 | 114.297 | 35.303 | 42.349 | 191.949 | 114.297 | 35.140 | 42.349 | 191.786 | 11 | 56 | 16 |
| **44** | 146 | **f** | 21 | 100.000 | 1.982.753 | 175.684 | 91.396 | 61.945 | 329.025 | 175.684 | 91.203 | 61.945 | 328.832 | 28 | 169 | 14,5 |
| **45** | 147 | **f** | 50 | 100.000 | 1.697.764 | 155.454 | 77.983 | 85.572 | 319.009 | 155.454 | 77.782 | 85.572 | 318.808 | 15 | 181 | 23 |
| **46** | 148 | **f** | 24 | 100.000 | 1.694.446 | 151.025 | 68.659 | 80.809 | 300.493 | 151.025 | 68.538 | 80.809 | 300.372 | 18 | 142,5 | 23 |
| **47** | 151 | **f** | 49 | 100.000 | 1.996.229 | 118.478 | 32.698 | 30.188 | 181.364 | 118.478 | 32.629 | 30.188 | 181.295 | 9 | 69 | 18 |
| **48** | 176 | **f** | 20 | 100.000 | 1.762.958 | 8.929 | 31.978 | 29.837 | 70.744 | 8.929 | 31.824 | 29.837 | 70.590 | 8 | 64 | 24 |
| **49** | 188 | **f** | 38 | 100.000 | 1.922.921 | 122.305 | 52.948 | 27.320 | 202.573 | 122.305 | 52.846 | 27.320 | 202.471 | 13 | 77 | 24 |
| **50** | 189 | **m** | 18 | 100.000 | 1.588.345 | 46.417 | 25.598 | 18.358 | 90.373 | 46.417 | 25.563 | 18.358 | 90.338 | 9 | 117 | 21 |
| **51** | 190 | **m** | 41 | 100.000 | 1.504.210 | 107.109 | 31.250 | 32.478 | 170.837 | 107.109 | 31.161 | 32.478 | 170.748 | 20 | 118 | 36,5 |
| **52** | 191 | **f** | 20 | 100.000 | 1.333.144 | 44.701 | 20.089 | 25.287 | 90.077 | 44.701 | 20.044 | 25.287 | 90.032 | 6 | 64 | 17 |
| **53** | 192 | **m** | 35 | 100.000 | 1.843.561 | 84.469 | 29.716 | 26.142 | 140.327 | 84.469 | 29.680 | 26.142 | 140.291 | 12 | 84 | 16,5 |
| **54** | 193 | **m** | 24 | 100.000 | 1.861.370 | 39.356 | 21.967 | 22.415 | 83.738 | 39.356 | 21.921 | 22.415 | 83.692 | 4 | 45 | 17 |
| **55** | 194 | **f** | 24 | 100.000 | 1.498.888 | 55.079 | 16.517 | 33.282 | 104.878 | 55.079 | 16.483 | 33.282 | 104.844 | 7 | 61,5 | 20 |
| **56** | 195 | **m** | 20 | 100.000 | 1.373.436 | 29.816 | 15.571 | 16.743 | 62.130 | 29.816 | 15.544 | 16.743 | 62.103 | 6 | 42 | 14 |
| **57** | 196 | **m** | 28 | 100.000 | 1.459.168 | 31.937 | 21.143 | 16.355 | 69.435 | 31.937 | 21.120 | 16.355 | 69.412 | 4 | 70 | 16 |
| **Minimum** | | | 18 | 100.000 | 1.321.306 | 8.929 | 11.399 | 5.374 | 29.361 | 8.929 | 11.333 | 5.374 | 29.295 | 2 | 20 | 6 |
| **Maximum** | | | 67 | 100.000 | 2.045.940 | 175.684 | 91.396 | 85.572 | 329.025 | 175.684 | 91.203 | 85.572 | 328.832 | 28 | 181 | 46 |
| **Mean** | | | 33 | 100.000 | **1.698.684** | 58.730 | 33.184 | 26.403 | 118.317 | 58.730 | 33.087 | 26.403 | 118.220 | 11 | 84 | 21 |
| **Median** | | | 30 | 100.000 | 1.694.446 | 46.417 | 31.250 | 22.415 | 96.358 | 46.417 | 31.161 | 22.415 | 96.270 | 9 | 74 | 20 |
| **Std. deviation** | | | 13 | 0 | 195.690 | 40.746 | 15.155 | 15.346 | 63.902 | 40.746 | 15.128 | 15.346 | 63.883 | 5 | 37 | 8 |
| **Sum** | | | - | 5.700.000 | **96.825.014** | 3.347.621 | 1.891.512 | 1.504.954 | 6.744.087 | 3.347.621 | 1.885.982 | 1.504.954 | 6.738.557 | - | - | - |

**Table S2: Healthy cohort demographics and and sequencing statistics - continued**

| # | Internal ID | Sex | Age | Cell Number | Raw reads | Sequences with ≥3 reads per UMI | | | | Productive sequences | | | | Unique productive CDR3s | | | |
|---|---|---|---|---|---|---|---|---|---|---|---|---|---|---|---|---|---|
| | | | | | Total | H | K | L | Total | H | K | L | Total | H | K | L | Total |
| **1** | 75 | **f** | 47 | 100.000 | 1.470.930 | 3.966 | 12.257 | 6.266 | 22.489 | 3.750 | 11.394 | 5.749 | 20.893 | 3.223 | 5.783 | 3.440 | 12.446 |
| **2** | 76 | **f** | 51 | 100.000 | 1.749.311 | 3.550 | 11.682 | 6.376 | 21.608 | 3.345 | 10.769 | 5.874 | 19.988 | 2.891 | 5.119 | 3.173 | 11.183 |
| **3** | 84 | **m** | 47 | 100.000 | 1.655.132 | 3.883 | 9.825 | 5.006 | 18.714 | 3.653 | 9.207 | 4.547 | 17.407 | 2.859 | 4.958 | 2.893 | 10.710 |
| **4** | 85 | **m** | 23 | 100.000 | 1.710.794 | 2.474 | 9.372 | 3.181 | 15.027 | 2.316 | 8.758 | 2.871 | 13.945 | 1.921 | 4.927 | 1.969 | 8.817 |
| **5** | 86 | **m** | 47 | 100.000 | 1.472.918 | 1.811 | 5.144 | 1.851 | 8.806 | 1.655 | 4.514 | 1.668 | 7.837 | 1.389 | 2.986 | 1.254 | 5.629 |
| **6** | 90 | **f** | 25 | 100.000 | 1.728.880 | 5.617 | 17.204 | 8.924 | 31.745 | 5.379 | 16.376 | 8.308 | 30.063 | 3.403 | 7.626 | 4.085 | 15.114 |
| **7** | 91 | **m** | 55 | 100.000 | 1.710.077 | 6.662 | 22.231 | 7.194 | 36.087 | 6.418 | 21.156 | 6.767 | 34.341 | 3.668 | 7.594 | 2.607 | 13.869 |
| **8** | 92 | **f** | 54 | 100.000 | 1.694.506 | 5.390 | 17.716 | 9.350 | 32.456 | 5.119 | 16.968 | 8.643 | 30.730 | 4.152 | 7.342 | 4.627 | 16.121 |
| **9** | 93 | **m** | 34 | 100.000 | 2.004.033 | 5.571 | 16.883 | 10.358 | 32.812 | 5.357 | 16.202 | 9.627 | 31.186 | 4.678 | 8.036 | 5.449 | 18.163 |
| **10** | 94 | **m** | 31 | 100.000 | 2.045.940 | 4.392 | 18.570 | 5.588 | 28.550 | 4.200 | 17.409 | 5.198 | 26.807 | 3.454 | 9.205 | 2.960 | 15.619 |
| **11** | 95 | **m** | 19 | 100.000 | 1.739.870 | 8.095 | 18.704 | 13.575 | 40.374 | 7.817 | 17.815 | 12.431 | 38.063 | 6.588 | 7.458 | 5.832 | 19.878 |
| **12** | 96 | **m** | 55 | 100.000 | 1.965.848 | 3.771 | 18.747 | 7.383 | 29.901 | 3.568 | 17.927 | 6.787 | 28.282 | 2.992 | 8.221 | 4.059 | 15.272 |
| **13** | 97 | **f** | 22 | 100.000 | 1.677.943 | 4.931 | 13.070 | 7.834 | 25.835 | 4.689 | 12.482 | 7.222 | 24.393 | 3.833 | 5.922 | 4.052 | 13.807 |
| **14** | 98 | **m** | 34 | 100.000 | 1.672.543 | 4.869 | 12.148 | 8.117 | 25.134 | 4.645 | 11.635 | 7.529 | 23.809 | 3.983 | 6.131 | 4.329 | 14.443 |
| **15** | 99 | **m** | 51 | 100.000 | 1.746.925 | 5.710 | 19.462 | 9.340 | 34.512 | 5.478 | 18.725 | 8.770 | 32.973 | 4.626 | 7.986 | 4.416 | 17.028 |
| **16** | 100 | **m** | 46 | 100.000 | 1.953.231 | 5.691 | 17.919 | 8.011 | 31.621 | 5.422 | 17.304 | 7.513 | 30.239 | 4.042 | 7.344 | 3.936 | 15.322 |
| **17** | 101 | **m** | 46 | 100.000 | 1.895.250 | 5.074 | 16.421 | 6.958 | 28.453 | 4.811 | 15.278 | 6.346 | 26.435 | 3.759 | 7.463 | 3.476 | 14.698 |
| **18** | 102 | **m** | 31 | 100.000 | 2.030.669 | 3.171 | 8.735 | 7.930 | 19.836 | 2.953 | 8.311 | 7.345 | 18.609 | 2.426 | 4.501 | 4.441 | 11.368 |
| **19** | 103 | **m** | 25 | 100.000 | 1.692.152 | 6.489 | 20.505 | 14.981 | 41.975 | 6.248 | 19.546 | 13.915 | 39.709 | 4.282 | 8.049 | 5.245 | 17.576 |
| **20** | 104 | **m** | 21 | 100.000 | 1.601.433 | 3.427 | 8.987 | 7.373 | 19.787 | 3.246 | 8.575 | 6.753 | 18.574 | 2.770 | 4.890 | 3.954 | 11.614 |
| **21** | 105 | **m** | 20 | 100.000 | 1.321.306 | 4.675 | 13.007 | 10.731 | 28.413 | 4.480 | 12.436 | 9.864 | 26.780 | 3.945 | 6.120 | 5.360 | 15.425 |
| **22** | 106 | **f** | 48 | 100.000 | 1.685.813 | 6.162 | 18.771 | 14.737 | 39.670 | 5.927 | 18.086 | 13.894 | 37.907 | 4.875 | 7.850 | 6.729 | 19.454 |
| **23** | 117 | **f** | 24 | 100.000 | 1.483.778 | 5.530 | 16.165 | 10.587 | 32.282 | 5.306 | 15.442 | 9.622 | 30.370 | 3.812 | 6.189 | 4.510 | 14.511 |
| **24** | 118 | **m** | 30 | 100.000 | 1.632.990 | 6.693 | 19.169 | 11.510 | 37.372 | 6.426 | 18.244 | 10.622 | 35.292 | 5.389 | 8.192 | 5.255 | 18.836 |
| **25** | 119 | **f** | 28 | 100.000 | 1.625.973 | 5.793 | 22.314 | 11.529 | 39.636 | 5.539 | 21.402 | 10.708 | 37.649 | 4.267 | 8.846 | 5.107 | 18.220 |
| **26** | 120 | **m** | 21 | 100.000 | 1.749.428 | 7.341 | 18.681 | 11.177 | 37.199 | 7.001 | 17.841 | 10.316 | 35.158 | 5.922 | 7.938 | 5.348 | 19.208 |
| **27** | 121 | **f** | 21 | 100.000 | 1.576.639 | 5.341 | 20.127 | 8.256 | 33.724 | 5.101 | 19.368 | 7.635 | 32.104 | 4.556 | 9.260 | 4.528 | 18.344 |
| **28** | 126 | **f** | 24 | 100.000 | 1.932.765 | 6.243 | 16.884 | 11.844 | 34.971 | 5.892 | 15.943 | 11.000 | 32.835 | 4.076 | 5.912 | 4.749 | 14.737 |
| **29** | 127 | **m** | 42 | 100.000 | 1.661.332 | 3.398 | 11.643 | 4.697 | 19.738 | 3.201 | 11.102 | 4.399 | 18.702 | 2.683 | 5.372 | 2.526 | 10.581 |
| **30** | 128 | **f** | 67 | 100.000 | 2.026.801 | 4.163 | 14.141 | 7.414 | 25.718 | 3.936 | 13.516 | 6.809 | 24.261 | 3.388 | 7.725 | 4.175 | 15.288 |
| **31** | 130 | **m** | 37 | 100.000 | 1.769.184 | 4.763 | 13.580 | 6.754 | 25.097 | 4.558 | 13.045 | 6.261 | 23.864 | 3.917 | 6.160 | 3.490 | 13.567 |
| **32** | 131 | **f** | 67 | 100.000 | 1.544.258 | 4.529 | 12.403 | 8.832 | 25.764 | 4.345 | 11.930 | 8.171 | 24.446 | 3.852 | 5.989 | 4.723 | 14.564 |
| **33** | 132 | **m** | 39 | 100.000 | 1.863.306 | 6.606 | 17.563 | 12.208 | 36.377 | 6.325 | 16.898 | 11.344 | 34.567 | 4.946 | 7.496 | 5.666 | 18.108 |
| **34** | 133 | **m** | 35 | 100.000 | 1.478.049 | 6.863 | 20.318 | 10.992 | 38.173 | 6.571 | 19.396 | 10.051 | 36.018 | 5.680 | 8.216 | 5.259 | 19.155 |
| **35** | 134 | **m** | 36 | 100.000 | 1.566.603 | 6.859 | 20.330 | 11.899 | 39.088 | 6.538 | 19.430 | 10.931 | 36.899 | 5.762 | 9.062 | 5.768 | 20.592 |
| **36** | 135 | **f** | 22 | 100.000 | 1.372.889 | 6.020 | 16.761 | 8.674 | 31.455 | 5.728 | 16.126 | 7.893 | 29.747 | 4.778 | 6.958 | 3.975 | 15.711 |
| **37** | 136 | **f** | 20 | 100.000 | 1.657.997 | 10.400 | 17.747 | 18.633 | 46.780 | 9.996 | 16.930 | 17.206 | 44.132 | 7.369 | 7.159 | 7.274 | 21.802 |
| **38** | 137 | **f** | 21 | 100.000 | 1.436.791 | 10.171 | 28.724 | 15.420 | 54.315 | 9.823 | 27.669 | 14.422 | 51.914 | 6.944 | 9.467 | 5.435 | 21.846 |
| **39** | 138 | **f** | 26 | 100.000 | 1.465.807 | 6.183 | 15.278 | 10.417 | 31.878 | 5.917 | 14.523 | 9.600 | 30.040 | 4.132 | 6.324 | 4.500 | 14.956 |
| **40** | 139 | **f** | 36 | 100.000 | 1.746.660 | 4.911 | 16.701 | 11.381 | 32.993 | 4.646 | 15.363 | 10.458 | 30.467 | 3.748 | 6.772 | 5.160 | 15.680 |
| **41** | 140 | **m** | 23 | 100.000 | 1.993.175 | 5.077 | 16.561 | 8.408 | 30.046 | 4.810 | 15.870 | 7.704 | 28.384 | 4.065 | 7.496 | 4.237 | 15.798 |
| **42** | 141 | **m** | 21 | 100.000 | 1.552.022 | 6.432 | 15.721 | 6.830 | 28.983 | 6.036 | 13.853 | 5.976 | 25.865 | 5.252 | 6.905 | 3.257 | 15.414 |
| **43** | 145 | **f** | 21 | 100.000 | 1.943.870 | 4.984 | 13.038 | 8.308 | 26.330 | 4.701 | 12.020 | 7.343 | 24.064 | 3.836 | 5.990 | 4.018 | 13.844 |
| **44** | 146 | **f** | 21 | 100.000 | 1.982.753 | 10.115 | 31.029 | 11.232 | 52.376 | 9.671 | 29.074 | 10.152 | 48.897 | 7.387 | 10.249 | 4.489 | 22.125 |
| **45** | 147 | **f** | 50 | 100.000 | 1.697.764 | 8.055 | 22.325 | 14.875 | 45.255 | 7.715 | 21.479 | 13.923 | 43.117 | 5.379 | 8.872 | 6.025 | 20.276 |
| **46** | 148 | **f** | 24 | 100.000 | 1.694.446 | 7.893 | 21.115 | 16.756 | 45.764 | 7.570 | 19.976 | 15.154 | 42.700 | 6.112 | 8.198 | 7.591 | 21.901 |
| **47** | 151 | **f** | 49 | 100.000 | 1.996.229 | 5.112 | 13.465 | 8.671 | 27.248 | 4.822 | 12.803 | 7.979 | 25.604 | 3.507 | 6.624 | 4.654 | 14.785 |
| **48** | 176 | **f** | 20 | 100.000 | 1.762.958 | 4.173 | 12.887 | 10.088 | 27.148 | 4.021 | 12.048 | 9.180 | 25.249 | 3.748 | 5.741 | 5.050 | 14.539 |
| **49** | 188 | **f** | 38 | 100.000 | 1.922.921 | 4.595 | 14.268 | 9.125 | 27.988 | 4.342 | 13.643 | 8.551 | 26.536 | 3.203 | 6.741 | 4.898 | 14.842 |
| **50** | 189 | **m** | 18 | 100.000 | 1.588.345 | 3.673 | 11.223 | 5.162 | 20.058 | 3.447 | 10.271 | 4.683 | 18.401 | 2.811 | 5.513 | 2.887 | 11.211 |
| **51** | 190 | **m** | 41 | 100.000 | 1.504.210 | 5.460 | 15.065 | 9.192 | 29.717 | 5.165 | 14.509 | 8.599 | 28.273 | 3.636 | 6.306 | 4.074 | 14.016 |
| **52** | 191 | **f** | 20 | 100.000 | 1.333.144 | 3.013 | 10.127 | 9.065 | 22.205 | 2.821 | 9.617 | 8.378 | 20.816 | 2.212 | 5.162 | 4.512 | 11.886 |
| **53** | 192 | **m** | 35 | 100.000 | 1.843.561 | 4.609 | 11.502 | 7.314 | 23.425 | 4.306 | 10.936 | 6.689 | 21.931 | 3.640 | 5.859 | 4.145 | 13.644 |
| **54** | 193 | **m** | 24 | 100.000 | 1.861.370 | 3.448 | 7.356 | 5.612 | 16.416 | 3.176 | 6.983 | 5.137 | 15.296 | 2.323 | 4.304 | 3.464 | 10.091 |
| **55** | 194 | **f** | 24 | 100.000 | 1.498.888 | 3.169 | 7.618 | 5.517 | 16.304 | 2.980 | 7.202 | 5.057 | 15.239 | 2.574 | 3.953 | 3.135 | 9.662 |
| **56** | 195 | **m** | 20 | 100.000 | 1.373.436 | 2.502 | 6.531 | 3.946 | 12.979 | 2.341 | 6.190 | 3.588 | 12.119 | 1.830 | 3.071 | 2.008 | 6.909 |
| **57** | 196 | **m** | 28 | 100.000 | 1.459.168 | 2.427 | 8.641 | 4.012 | 15.080 | 2.256 | 8.261 | 3.673 | 14.190 | 1.784 | 4.078 | 2.260 | 8.122 |
| **Minimum** | | | 18 | 100.000 | 1.321.306 | 1.811 | 5.144 | 1.851 | 8.806 | 1.655 | 4.514 | 1.668 | 7.837 | 1.389 | 2.986 | 1.254 | 5.629 |
| **Maximum** | | | 67 | 100.000 | 2.045.940 | 10.400 | 31.029 | 18.633 | 54.315 | 9.996 | 29.074 | 17.206 | 51.914 | 7.387 | 10.249 | 7.591 | 22.125 |
| **Mean** | | | 33 | 100.000 | **1.698.684** | 5.297 | 15.515 | 9.077 | 29.889 | 5.044 | 14.733 | 8.366 | 28.142 | 4.005 | 6.730 | 4.323 | 15.058 |
| **Median** | | | 30 | 100.000 | 1.694.446 | 5.077 | 16.165 | 8.674 | 29.717 | 4.811 | 15.278 | 7.979 | 28.273 | 3.836 | 6.772 | 4.416 | 14.956 |
| **Std. deviation** | | | 13 | 0 | 195.690 | 1.851 | 5.083 | 3.391 | 9.545 | 1.799 | 4.889 | 3.152 | 9.121 | 1.351 | 1.606 | 1.248 | 3.767 |
| **Sum** | | | - | 5.700.000 | **96.825.014** | 301.925 | 884.361 | 517.401 | 1.703.687 | 287.505 | 839.776 | 476.835 | 1.604.116 | 228.279 | 383.610 | 246.438 | 858.327 |

Table S3: Viral panel overview

| | | | Panel | | |
|---|---|---|---|---|---|
| **Virus name** | **Subtype** | **Tier** | **118** | **56** | **12** |
| **0260_V5_C36** | A1 | nd | | | |
| **0330_V4_C3** | A1 | 2 | | | |
| **1006_11_C3_1601** | B | 2 | | | |
| **1012_11_TC21_3257** | B | 1B or 2 | | | |
| **1054_07_TC4_1499** | B | 2 | | | |
| **1056_10_TA11_1826** | B | 1B or 2 | | | |
| **1394_C9G1** | C | 2 | | | |
| **16055_2_3** | C | 2 | | | |
| **191084_B7_19** | A1 | 2 | | | |
| **191821_E6_1** | D | 2 | | | |
| **191955_A11** | A1 | 2 | | | |
| **211_9** | 02_AG | 2 | | | |
| **231966_C2** | D | 2 | | | |
| **249M_B10** | C | 2 | | | |
| **3103_V3_C10** | A1CD | 2 | | | |
| **6041_V3_C23** | A1C | 2 | | | |
| **62357_14_D3_4589** | B | 2 | | | |
| **6240_08_TA5_4622** | B | 2 | | | |
| **6244_13_B5_4567** | B | 2 | | | |
| **6480_V4_C25** | CD | 2 | | | |
| **6545_V4_C1** | A1C | nd | | | |
| **6811_V7_C18** | CD | 2 | | | |
| **6952_V1_C20** | 41_CD | 2 | | | |
| **703010054_2A2** | C | 2 | | | |
| **703010200_1E5** | C | 2 | | | |
| **704809221_1B3** | C | 2 | | | |
| **89_F1_2_25** | CD | 2 | | | |
| **9004SS_A3_4** | A1 | 2 | | | |
| **A07412M1_VRC12** | D | 2 | | | |
| **BF1266_431A** | C | 2 | | | |
| **BJOX009000_02_4** | 01_AE | 2 | | | |
| **BJOX010000_06_2** | 01_AE | 2 | | | |
| **BJOX015000_11_5** | 01_AE | 2 | | | |
| **BJOX025000_01_1** | 01_AE | 2 | | | |
| **BJOX028000_10_3** | 01_AE | 2 | | | |
| **C2101_C1** | 01_AE | 2 | | | |
| **C3347_C11** | 01_AE | 2 | | | |
| **CE0393_C3** | C | 2 | | | |
| **CE0682_E4** | C | 2 | | | |
| **CE1086_B2** | C | 2 | | | |
| **CE1172_H1** | C | 2 | | | |
| **CE2010_F5** | C | 2 | | | |
| **CE2060_G9** | C | 2 | | | |
| **CNE17** | C | 2 | | | |
| **CNE19** | 07_BC | 2 | | | |
| **CNE20** | 07_BC | 2 | | | |
| **CNE21** | 07_BC | 2 | | | |
| **CNE52** | 08_BC | 2 | | | |
| **P0402_C2_11** | G | 2 | | | |
| **P1981_C5_3** | G | 2 | | | |
| **R3265_C6** | 01_AE | 2 | | | |
| **SC05_8C11_2344** | B | 2 | | | |
| **THRO4156** | B | 2 | | | |
| **WEAU_D15_410** | B | 2 | | | |
| **X1193_C1** | G | 2 | | | |
| **X1254_C3** | G | 2 | | | |
| **X2131_C1_B5** | G | 2 | | | |
| **ZM246F** | C | 2 | | | |
| **ZM247_V1** | C | 2 | | | |

Table S3: Viral panel overview - continued

| | | | | | |
|---|---|---|---|---|---|
| **0013095_2_11** | C | 2 | Included | Included | Not included |
| **001428_2_42** | C | 2 | Included | Included | Not included |
| **0815_V3_C3** | A1D | 2 | Included | Included | Not included |
| **16845_2_22** | C | 2 | Included | Included | Not included |
| **231965_C1** | D | 2 | Included | Included | Not included |
| **235_47** | 02_AG | 2 | Included | Included | Not included |
| **263_8** | 02_AG | 2 | Included | Included | Not included |
| **3016_V5_C45** | D | 2 | Included | Included | Not included |
| **3301_V1_C24** | C | 2 | Included | Included | Not included |
| **3817_V2_C59** | CD | 2 | Included | Included | Not included |
| **620345_C1** | 01_AE | 2 | Included | Included | Not included |
| **6535_3** | B | 1B or 2 | Included | Included | Not included |
| **6540_V4_C1** | A1C | 2 | Included | Included | Not included |
| **928_28** | 02_AG | 2 | Included | Included | Not included |
| **AC10_29** | B | 2 | Included | Included | Not included |
| **C1080_C3** | 01_AE | 2 | Included | Included | Not included |
| **C4118_9** | 01_AE | 2 | Included | Included | Not included |
| **CAAN5342** | B | 2 | Included | Included | Not included |
| **CAP210_E8** | C | 2 | Included | Included | Not included |
| **CAP45_G3** | C | 2 | Included | Included | Not included |
| **CNE30** | C | 2 | Included | Included | Not included |
| **CNE5** | 01_AE | 2 | Included | Included | Not included |
| **CNE53** | 08_BC | 2 | Included | Included | Not included |
| **CNE58** | C | 2 | Included | Included | Not included |
| **DU156_12** | C | 2 | Included | Included | Not included |
| **DU172_17** | C | 2 | Included | Included | Not included |
| **DU422_1** | C | 2 | Included | Included | Not included |
| **MS208_A1** | A1D | 1 or 2 | Included | Included | Not included |
| **PVO_4** | B | 2 or 3 | Included | Included | Not included |
| **Q23_17** | A1 | 1B | Included | Included | Not included |
| **Q259_17** | A1 | 2 | Included | Included | Not included |
| **Q461_E2** | A1 | 2 | Included | Included | Not included |
| **Q769_D22** | A1 | 2 | Included | Included | Not included |
| **Q842_D12** | A1 | 2 | Included | Included | Not included |
| **QH0692_42** | B | 2 | Included | Included | Not included |
| **R1166_C1** | 01_AE | 2 | Included | Included | Not included |
| **R2184_C4** | 01_AE | 2 | Included | Included | Not included |
| **REJO4541_67** | B | 2 | Included | Included | Not included |
| **RHPA4259_7** | B | 2 | Included | Included | Not included |
| **SC422_8** | B | 2 | Included | Included | Not included |
| **T250_4** | 02_AG | 2 | Included | Included | Not included |
| **T251_18** | 02_AG | 2 or 3 | Included | Included | Not included |
| **T255_34** | 02_AG | 2 | Included | Included | Not included |
| **T257_31** | 02A1 | 2 or 3 | Included | Included | Not included |
| **T278_50** | 02_AG | 2 or 3 | Included | Included | Not included |
| **TRJO4551_58** | B | 3 | Included | Included | Not included |
| **WITO4160_33** | B | 2 | Included | Included | Not included |
| **X2088_C9** | G | 2 | Included | Included | Not included |
| **ZM109_4** | C | 1B or 2 | Included | Included | Not included |
| **ZM135_10A** | C | 2 | Included | Included | Not included |
| **ZM197_7** | C | 1B or 2 | Included | Included | Not included |
| **ZM214_15** | C | 2 | Included | Included | Not included |
| **ZM233_6** | C | 2 | Included | Included | Not included |
| **ZM249_1** | C | 2 | Included | Included | Not included |
| **ZM53_12** | C | 2 | Included | Included | Not included |
| **TRO_11** | B | 2 | Included | Included | Included |
| **CE1176_A3** | C | 2 | Included | Not included | Included |
| **CNE8** | 01_AE | 2 or 3 | Included | Not included | Included |
| **X1632_S2_B10** | G | 2 | Included | Not included | Included |
| **CNE55** | 01_AE | 2 | Not included | Not included | Included |
| **246_F3_C10_2** | A1C | 2 | Not included | Not included | Included |
| **CH119_10** | 07_BC | 2 | Not included | Not included | Included |
| **X2278_C2_B6** | B | 2 | Not included | Not included | Included |
| **703010217_B6** | C | 2 | Not included | Not included | Included |
| **BJOX002000_03_2** | 07_BC | 2 | Not included | Not included | Included |
| **25710_2_43** | C | 1B or 2 | Not included | Not included | Included |
| **398_F1_F6_20** | A1 | 2 | Not included | Not included | Included |

**Legend:** Incldued (blue) / Not included (grey)

**Table S4: Features of HIV-1 broadly neutralizing antibodies**

| Antibody information | | | | | Neutralization 56-strain panel (CATNAP) | | | Heavy chain | | | | | |
|---|---|---|---|---|---|---|---|---|---|---|---|---|---|
| # | Name | Binding site | Donor | Reference | Coverage (%) | geo. mean IC50 (ng/µl) | normed AUC | V gene | D gene | J gene | V gene germline identity (%) | CDR3 amino acid sequence | CDR3 length (aa) |
| 1 | **10-1074** | V3-glycan | Donor 17 | Mouquet et al., 2012 | 67,9 | 0,126 | 63,6 | IGHV4-59 | IGHD3-3 | IGHJ6 | 83,9 | ATARRGQRIYGVVSFGEFFYYYSMDV | 26 |
| 2 | **10E8** | MPER | Donor N152 | Huang et al., 2012 | 98,2 | 0,392 | 75,3 | IGHV3-15 | IGHD3-3 | IGHJ1 | 78,5 | ARTGKYYDFWSGYPPGEEYFQD | 22 |
| 3 | **12A12** | CD4bs | Patient 12 | Scheid et al., 2011 | 92,9 | 0,195 | 80,4 | IGHV1-2 | IGHD1-26 | IGHJ5 | 78,7 | ARDGSGDDTSWHLDP | 15 |
| 4 | **179NC75** | CD4bs | EB179 | Freund et al., 2015 | 30,4 | 0,135 | 29 | IGHV3-21 | IGHD3-9 | IGHJ5 | 100,0 | ARVHAWRYFDWVNRRSPVEKVLSIDP | 26 |
| 5 | **1B2530** | CD4bs | Patient 1 | Scheid et al., 2011 | 67,9 | 6,331 | 26,2 | IGHV1-46 | IGHD6-13 | IGHJ5 | 72,7 | ARAEAASDSHSRPIMFDH | 18 |
| 6 | **3074** | V3-glycan | n/a | Gorny et al., 2006 | 16,1 | 13,193 | 5 | IGHV4-59 | IGHD3-22 | IGHJ3 | 94,2 | ARDFGEYHYDGRGFQCEGFDL | 21 |
| 7 | **35O22** | Interface/FP | Donor N152 | Huang et al., 2014 | 62,5 | 0,594 | 45,2 | IGHV1-18 | IGHD3-10 | IGHJ1 | 69,9 | AKGLLRDGSSTWLPYL | 16 |
| 8 | **3BNC117** | CD4bs | Patient 3 | Scheid et al., 2011 | 89,3 | 0,103 | 85,3 | IGHV1-2 | IGHD6-25 | IGHJ2 | 72,5 | ARQRSDYWDFDV | 12 |
| 9 | **447-52D** | V3-glycan | n/a | Buchbinder et al., 1992 | 10,7 | 16,991 | 3,3 | IGHV3-15 | IGHD3-10 | IGHJ6 | 94,7 | TTDGFIMIRGVSEDYYYYYMDV | 22 |
| 10 | **4E10** | MPER | n/a | Buchacher et al., 1994 | 98,2 | 1,389 | 58,1 | IGHV1-69 | IGHD6-19 | IGHJ4 | 87,8 | AREGTTGWGWLGKPIGAFAH | 20 |
| 11 | **561_01_18** | CD4bs | IDC561 | Schommers et al., 2020 | 94,6 | 0,049 | 100 | IGHV1-46 | IGHD3-10 | IGHJ6 | 68,1 | ARDPFGDRAPHYNYHMDV | 18 |
| 12 | **561_01_55** | CD4bs | IDC561 | Schommers et al., 2020 | 89,3 | 0,105 | 84,9 | IGHV1-46 | IGHD3-10 | IGHJ6 | 68,4 | ARDPFGDMYPHYNYHMDV | 18 |
| 13 | **561_02_12** | CD4bs | IDC561 | Schommers et al., 2020 | 83,9 | 0,047 | 89,2 | IGHV1-46 | IGHD3-16 | IGHJ6 | 73,9 | ARDPFGETFRGHDQPYRMDV | 20 |
| 14 | **8ANC131** | CD4bs | Patient 8 | Scheid et al., 2011 | 69,6 | 1,659 | 40 | IGHV1-46 | IGHD3-16 | IGHJ6 | 73,6 | ARDGLGEVAPDYRYGIDV | 18 |
| 15 | **8ANC195** | Interface/FP | Patient 8 | Scheid et al., 2011 | 62,5 | 1,037 | 40 | IGHV1-69 | IGHD3-3 | IGHJ4 | 68,7 | TTTSTYDKWSGLHHDGVMAFSS | 22 |
| 16 | **BG1** | V2-apex | EB354 | Freund et al., 2017 | 41,1 | 0,615 | 29,8 | IGHV3-49 | IGHD2-21 | IGHJ6 | 72,8 | AREQRNKDYRYGQEGFGYSYGMDV | 24 |
| 17 | **BG18** | V3-glycan | EB354 | Freund et al., 2017 | 62,5 | 0,058 | 65,1 | IGHV4-4 | IGHD3-3 | IGHJ6 | 78,8 | ARNAIRIYGVVALGEWFHYGMDV | 23 |
| 18 | **BG8** | V3-glycan | EB354 | Freund et al., 2017 | 58,9 | 0,195 | 51,4 | IGHV4-4 | IGHD3-3 | IGHJ6 | 74,6 | ARNVIRVFGVISLGEWFHYGMDV | 23 |
| 19 | **CH01** | V2-apex | CH0219 | Bonsignori et al., 2011 | 58,9 | 1,096 | 37,4 | IGHV3-20 | IGHD3-10 | IGHJ2 | 83,7 | ARGTDYTIDDAGIHYQGSGTFWYFDL | 26 |
| 20 | **CH103** | CD4bs | Donor CH505 | Liao et al., 2013 | 87,5 | 9,579 | 28,6 | IGHV4-59 | IGHD3-3 | IGHJ4 | 83,1 | ASLPRGQLVNAYFRN | 15 |
| 21 | **CH235** | CD4bs | Donor CH505 | Gao et al., 2014 | 23,2 | 6,474 | 9 | IGHV1-46 | IGHD1-26 | IGHJ4 | 92,2 | ARNVATEGSLLHYDY | 15 |
| 22 | **CH235.12** | CD4bs | Donor CH505 | Bonsignori et al., 2016 | 87,5 | 0,754 | 59,3 | IGHV1-46 | IGHD6-13 | IGHJ4 | 74,8 | VRNVGTAGSLLHYDH | 15 |
| 23 | **CH235.9** | CD4bs | Donor CH505 | Bonsignori et al., 2016 | 76,8 | 3,326 | 36,3 | IGHV1-46 | IGHD1-1 | IGHJ4 | 80,6 | VRNVGTAGSLLHYDH | 15 |
| 24 | **DH270.1** | V3-glycan | CH848 | Bonsignori et al., 2017 | 39,3 | 0,761 | 27,1 | IGHV1-2 | IGHD3-22 | IGHJ4 | 94,2 | TTGGWIGLYSDTSGYPNFDY | 20 |
| 25 | **DH270.5** | V3-glycan | CH848 | Bonsignori et al., 2017 | 44,6 | 0,367 | 35,1 | IGHV1-2 | IGHD3-22 | IGHJ4 | 88,5 | VTGAWISDYYDSSYYPNFDH | 20 |
| 26 | **DH270.6** | V3-glycan** | CH848** | Bonsignori et al., 2017 | 51,8 | 0,207 | 44,7 | IGHV1-2 | IGHD3-22 | IGHJ4 | 86,8 | TTGGWISLYYDSSYYPNFDH | 20 |
| 27 | **DH511.11P** | MPER | CH0210 | Williams et al., 2017 | 98,2 | 0,841 | 65,1 | IGHV3-15 | IGHD3-3 | IGHJ6 | 85,4 | TADEGAPILRFFEWGYYNYYMDV | 23 |
| 28 | **DH511.12P** | MPER | CH0210 | Williams et al., 2017 | 98,2 | 0,751 | 66,6 | IGHV3-15 | IGHD3-3 | IGHJ6 | 84,7 | TADEGAPILRFFEWGYYNYYMDV | 23 |
| 29 | **DH511.2** | MPER | CH0210 | Williams et al., 2017 | 98,2 | 1,073 | 61,8 | IGHV3-15 | IGHD4-4 | IGHJ6 | 80,5 | TMDEGTPVTRFLEWGYFYYYMAV | 23 |
| 30 | **HJ16** | CD4bs | VI3208 | Corti et al., 2010 | 42,9 | 3,078 | 21,4 | IGHV3-30 | IGHD3-3 | IGHJ2 | 70,9 | VKDQTIFHKNGAVDFFSYFDL | 21 |
| 31 | **IOMA** | CD4bs | R1 | Gristick et al., 2016 | 42,9 | 2,664 | 21,9 | IGHV1-2 | IGHD6-19 | IGHJ5 | 88,5 | AREMFDSSADWSPWRGMVA | 19 |
| 32 | **N6** | CD4bs | Z258 | Huang et al., 2016 | 98,2 | 0,064 | 100 | IGHV1-2 | IGHD3-10 | IGHJ5 | 70,8 | ARDRSYGDSSWALDA | 15 |
| 33 | **NIH45-46** | CD4bs | NIH45 | Scheid et al., 2011 | 80,4 | 0,116 | 80,5 | IGHV1-2 | IGHD2-8 | IGHJ1 | 67,2 | TRGKYCTARDYYNWDFEH | 18 |
| 34 | **PCDN-33A** | V3-glycan | PC76 | MacLeod et al., 2016 | 42,9 | 0,261 | 38,2 | IGHV4-34 | IGHD3-3 | IGHJ5 | 87,4 | ARGGRKICYDYWCGYVNNCFDT | 22 |
| 35 | **PCDN-38A** | V3-glycan | PC76 | MacLeod et al., 2016 | 35,7 | 0,681 | 27,2 | IGHV4-34 | IGHD2-8 | IGHJ5 | 85,7 | ARGGRKVYHPYWTGYVNNCFDP | 22 |
| 36 | **PCDN-38B** | V3-glycan | PC76 | MacLeod et al., 2016 | 42,9 | 0,493 | 34,3 | IGHV4-34 | IGHD3-3 | IGHJ5 | 82,9 | ARGGRKVYHAYWTGYVNNCFDP | 22 |
| 37 | **PG16** | V2-apex | Donor 24 | Walker et al., 2009 | 83,9 | 0,081 | 88,9 | IGHV3-33 | IGHD3-3 | IGHJ6 | 85,5 | AREAGGPIWHDDVKYYDFNDGYYNYHYMDV | 30 |
| 38 | **PG9** | V2-apex | Donor 24 | Walker et al., 2009 | 91,1 | 0,157 | 87,6 | IGHV3-33 | IGHD3-3 | IGHJ6 | 87,7 | VREAGGPDYRNGYNYYDFYDGYYNYHYMDV | 30 |
| 39 | **PGDM1400** | V2-apex | Donor 84 | Sok et al., 2014 | 82,1 | 0,028 | 100 | IGHV1-8 | IGHD3-16 | IGHJ6 | 74,0 | AKGSKHRLRDYALYDDDGALNWAVDVDYLSNLEF | 34 |
| 40 | **PGT121** | V3-glycan | Donor 17 | Walker et al., 2011 | 69,6 | 0,228 | 66,4 | IGHV4-59 | IGHD3-3 | IGHJ6 | 80,8 | ARTLHGRRIYGIVAFNEWFTYFYMDV | 26 |
| 41 | **PGT123** | V3-glycan | Donor 17 | Walker et al., 2011 | 53,6 | 0,067 | 61,4 | IGHV4-59 | IGHD3-10 | IGHJ6 | 77,4 | ARALHGKRIYGIVALGELFTYFYMDV | 26 |
| 42 | **PGT125** | V3-glycan | Donor 36 | Walker et al., 2011 | 35,7 | 0,073 | 41,1 | IGHV4-39 | IGHD1-20 | IGHJ5 | 74,9 | ARFDGEVLVYNHWPKPAWVDL | 21 |
| 43 | **PGT126** | V3-glycan | Donor 36 | Walker et al., 2011 | 48,2 | 0,181 | 47,8 | IGHV4-39 | IGHD2-8 | IGHJ5 | 77,5 | ARFDGEVLVYHDWPKPAWVDL | 21 |
| 44 | **PGT127** | V3-glycan | Donor 36 | Walker et al., 2011 | 33,9 | 0,186 | 33,9 | IGHV4-39 | IGHD1-1 | IGHJ5 | 79,7 | ARFGGEVLVYRDWPKPAWVDL | 21 |
| 45 | **PGT128** | V3-glycan | Donor 36 | Walker et al., 2011 | 64,3 | 0,225 | 61,5 | IGHV4-39 | IGHD1-26 | IGHJ5 | 76,6 | ARFGGEVLRYTDWPKPAWVDL | 21 |
| 46 | **PGT130** | V3-glycan | Donor 36 | Walker et al., 2011 | 57,1 | 0,353 | 50,6 | IGHV4-39 | IGHD3-22 | IGHJ5 | 81,1 | VRSGGDILYYYEWQKPHWFSP | 21 |
| 47 | **PGT135** | V3-glycan | Donor 39 | Walker et al., 2011 | 25,0 | 1,192 | 18,1 | IGHV4-39 | IGHD2-15 | IGHJ5 | 78,8 | ARHRHHDVFMLVPIAGWFDV | 20 |
| 48 | **PGT143** | V2-apex | Donor 84 | Walker et al., 2011 | 58,9 | 0,177 | 58,7 | IGHV1-8 | IGHD4-17 | IGHJ6 | 84,1 | TRGSKHRLRDYVLYDDYGLINYQEWNDYLEFLDV | 34 |
| 49 | **PGT145** | V2-apex | Donor 84 | Walker et al., 2011 | 80,4 | 0,220 | 77 | IGHV1-8 | IGHD4-17 | IGHJ6 | 83,3 | LTGSKHRLRDYFLYNEYGPNYEEWGDYLATLDV | 33 |
| 50 | **PGT151** | Interface/FP | Donor 31 | Falkowska et al., 2014 | 76,8 | 0,063 | 88,3 | IGHV3-30 | IGHD3-16 | IGHJ6 | 79,8 | ARMFQESGPPRLDRWSGRNYYYYSGMDV | 28 |
| 51 | **PGT152** | Interface/FP | Donor 31 | Falkowska et al., 2014 | 67,9 | 0,022 | 88,6 | IGHV3-30 | IGHD3-3 | IGHJ6 | 81,2 | ARMFQESGPPRFDSWSGRNYYYYSGMDV | 28 |
| 52 | **VRC-CH31** | CD4bs | CH0219 | Wu et al., 2011 | 80,4 | 0,225 | 76,3 | IGHV1-2 | IGHD1-26 | IGHJ1 | 70,0 | ARAQKRGRSEWAYAH | 15 |
| 53 | **VRC-PG04** | CD4bs | Donor 74 | Wu et al., 2011 | 85,7 | 0,367 | 74,8 | IGHV1-2 | IGHD3-16 | IGHJ2 | 70,0 | ARQKFYTGGQGWYFDL | 16 |
| 54 | **VRC-PG20** | CD4bs | IAVI 23 | Zhou et al., 2013 | 76,8 | 0,246 | 71,8 | IGHV1-2 | IGHD1-26 | IGHJ1 | 76,5 | ARRMRSQDREWDFQH | 15 |
| 55 | **VRC01** | CD4bs | NIH45 | Wu et al., 2010 | 85,7 | 0,318 | 76,6 | IGHV1-2 | IGHD3-16 | IGHJ1 | 68,3 | TRGKNCDYNWDFEH | 14 |
| 56 | **VRC03** | CD4bs | NIH45 | Wu et al., 2010 | 51,8 | 0,730 | 40,2 | IGHV1-2 | IGHD3-3 | IGHJ1 | 72,6 | DTAEYFCVRRGSCDYCGDFPWQY | 23 |
| 57 | **VRC06** | CD4bs | NIH45 | Li et al., 2012 | 35,7 | 2,696 | 20,6 | IGHV1-2 | IGHD2-21 | IGHJ1 | 66,1 | VRRGSSCPHCGDFHFEH | 17 |
| 58 | **VRC06b** | CD4bs | NIH45 | Li et al., 2012 | 44,6 | 0,528 | 36,9 | IGHV1-2 | IGHD3-3 | IGHJ1 | 72,2 | DTGEYFCVRKGPSCPHCGDFHWQH | 24 |
| 59 | **VRC07** | CD4bs | NIH45 | Rudicelli et al., 2014 | 87,5 | 0,119 | 91,2 | IGHV1-2 | IGHD2-8 | IGHJ1 | 67,7 | TRGKYCTARDYYNWDFEH | 18 |
| 60 | **VRC13** | CD4bs | 44 | Zhou et al., 2015 | 85,7 | 0,167 | 85,3 | IGHV1-69 | IGHD3-9 | IGHJ2 | 63,3 | ARERGRHFEPKNRDNLEGKFFDL | 23 |
| 61 | **VRC16** | CD4bs | C38 | Zhou et al., 2015 | 58,9 | 2,592 | 33,6 | IGHV3-23 | IGHD5-12 | IGHJ4 | 82,6 | VRVTFYHEGSGYYYRAGNYFDS | 22 |
| 62 | **VRC18** | CD4bs | C38 | Zhou et al., 2015 | 69,6 | 1,244 | 47,6 | IGHV1-2 | IGHD3-10 | IGHJ4 | 72,8 | ARGFAGYEWSFI | 12 |
| 63 | **VRC26.01** | V2-apex | CAP256 | Doria-Rose et al., 2014 | 23,2 | 2,081 | 14,7 | IGHV3-30 | IGHD3-9 | IGHJ3 | 92,5 | AKDVGDYKSDEWGTEYYDISISYPIQDPRAMVGAFDL | 37 |
| 64 | **VRC26.03** | V2-apex | CAP256 | Doria-Rose et al., 2014 | 42,9 | 0,068 | 49,2 | IGHV3-30 | IGHD3-3 | IGHJ3 | 91,9 | AKDLREDECEEWWSDYYDFGKQLPCRKSRGVAGIFDG | 37 |
| 65 | **VRC26.06** | V2-apex | CAP256 | Doria-Rose et al., 2014 | 16,1 | 0,442 | 14,5 | IGHV3-30 | IGHD3-16 | IGHJ3 | 89,8 | ARDLRELECEEWTLYNYYDFGSRGPCVDPRGVAGSFDV | 38 |
| 66 | **VRC26.08** | V2-apex | CAP256 | Doria-Rose et al., 2014 | 62,5 | 0,055 | 73,5 | IGHV3-30 | IGHD3-3 | IGHJ3 | 88,7 | VRDQREDECEEWWSDYYDFGRELPCRKFRGLGLAGIFDI | 39 |
| 67 | **VRC26.09** | V2-apex | CAP256 | Doria-Rose et al., 2014 | 57,1 | 0,040 | 70,4 | IGHV3-30 | IGHD3-3 | IGHJ3 | 86,1 | VKDQREDECEEWWSDYYDFGRELPCRKSRGLGLAGIFDM | 39 |
| 68 | **VRC26.25** | V2-apex | CAP256 | Doria-Rose et al., 2016 | 64,3 | 0,005 | 100 | IGHV3-30 | IGHD3-3 | IGHJ3 | 88,2 | AKDLREDECEEWWSDYYDFGKQLPCAKSRGGLVGIADN | 38 |
| 69 | **VRC26.26** | V2-apex | CAP256 | Doria-Rose et al., 2016 | 62,5 | 0,082 | 87,6 | IGHV3-30 | IGHD3-3 | IGHJ3 | 83,4 | VRDQREDECEEWWSDYYDFGKELPCRKFRGLGLAGIFDV | 39 |
| 70 | **VRC26.27** | V2-apex | CAP256 | Doria-Rose et al., 2016 | 53,6 | 0,007 | 100 | IGHV3-30 | IGHD3-3 | IGHJ3 | 84,5 | VRDQREDECEEWWSDYYDFGKELPCRKFRGLGLAGIFDI | 39 |
| 71 | **VRC34.01** | Interface/FP | N123 | Kong et al., 2016 | 46,4 | 0,222 | 83,2 | IGHV1-2 | IGHD3-10 | IGHJ6 | 85,5 | ARDKYYGNEAVGMDV | 15 |
| 72 | **VRC38.01** | V2-apex | N90 | Cale et al., 2017 | 35,7 | 0,258 | 64,5 | IGHV3-13 | IGHD6-19 | IGHJ6 | 82,2 | VRGPESGWFYHYYWGLGV | 18 |
| 73 | **b12** | CD4bs | Donor b | Burton et al., 1991 | 51,8 | 4,648 | 47,3 | IGHV1-3 | IGHD5-12 | IGHJ6 | 87,8 | ARVGPYSWDDSPQDNYYMDV | 20 |
| 74 | **3BNC55** | CD4bs | Patient 3 | Scheid et al., 2011 | 67,9 | 0,658 | 100 | IGHV1-2 | IGHD2-15 | IGHJ2 | 78,2 | ARRHSDYCDFDI | 12 |
| 75 | **NC37** | CD4bs | EB354 | Freund et al., 2017 | 35,7 | 0,426 | 100 | IGHV1-2 | IGHD6-19 | IGHJ6 | 72,9 | SRDNFGTRPVPGRGYYYGMDV | 21 |

Note: V(D)J information, V gene germline identity and CDR3 amino acid sequence were derived by annotation with IgBLAST according to the IMGT nomenclature and numbering.
n/a: not available
*For light chains, where no nucleotide sequence was available or IgBLAST annotation failed, data has been filled up with information deposited in the CATNAP database
** Epitope and Donor Information for DH270.6 was not deposited in the CATNAP database and therefore derived from the reference

**Table S4: Features of HIV-1 broadly neutralizing antibodies - continued**

| Antibody information | | | | | Light chain | | | | | |
|---|---|---|---|---|---|---|---|---|---|---|
| # | Name | Binding site | Donor | Reference | Isotype | V gene | J gene | V gene germline identity (%) | CDR3 amino acid sequence | CDR3 length (aa) |
| 1 | **10-1074** | V3-glycan | Donor 17 | Mouquet et al., 2012 | Lambda | IGLV3-21 | IGLJ2 | 80,5 | HMWDSRSGFSWS | 12 |
| 2 | **10E8** | MPER | Donor N152 | Huang et al., 2012 | Lambda | IGLV3-19 | IGLJ2 | 85,2 | SSRDKSGSRLSV | 12 |
| 3 | **12A12** | CD4bs | Patient 12 | Scheid et al., 2011 | Kappa | IGKV1-33 | IGKJ1 | 84,5 | AVLEF | 5 |
| 4 | **179NC75** | CD4bs | EB179 | Freund et al., 2015 | Lambda | IGLV3-1 | IGLJ2 | 100 | QAWDSKNYVT | 10 |
| 5 | **1B2530** | CD4bs | Patient 1 | Scheid et al., 2011 | Lambda | IGLV1-47 | IGLJ3 | 83,9 | ATYDSDGSIRL | 11 |
| 6 | **3074** | V3-glycan | n/a | Gorny et al., 2006 | Lambda* | IGLV1-51* | IGLV2 or 3* | n/a | ATWDGSLRTV* | 10 |
| 7 | **35O22** | Interface/FP | Donor N152 | Huang et al., 2014 | Lambda | IGLV2-23 | IGLJ1 | 77,3 | CSYTHNSGCV | 10 |
| 8 | **3BNC117** | CD4bs | Patient 3 | Scheid et al., 2011 | Kappa | IGKV1-33 | IGKJ3 | 80,7 | QVYEF | 5 |
| 9 | **447-52D** | V3-glycan | n/a | Buchbinder et al., 1992 | Lambda | IGLV1-51 | IGLJ3 | 97,6 | ATWDSGLSADWV | 12 |
| 10 | **4E10** | MPER | n/a | Buchacher et al., 1994 | Kappa | IGKV3-20 | IGKJ1 | 94,3 | QQYGQSLST | 9 |
| 11 | **561_01_18** | CD4bs | IDC561 | Schommers et al., 2020 | Kappa | IGKV3-20 | IGKJ4 | 79,1 | QRYGGTPIT | 9 |
| 12 | **561_01_55** | CD4bs | IDC561 | Schommers et al., 2020 | Kappa | IGKV3-20 | IGKJ4 | 78,4 | QRYGGTPIT | 9 |
| 13 | **561_02_12** | CD4bs | IDC561 | Schommers et al., 2020 | Kappa | IGKV3-20 | IGKJ4 | 80,6 | QSYGSITPLV | 10 |
| 14 | **8ANC131** | CD4bs | Patient 8 | Scheid et al., 2011 | Kappa | IGKV3-20 | IGKJ3 | 79,9 | QEYSSTPYN* | 9 |
| 15 | **8ANC195** | Interface/FP | Patient 8 | Scheid et al., 2011 | Kappa | IGKV1-5 | IGKJ5 | 84,9 | QQYDTYPGT | 9 |
| 16 | **BG1** | V2-apex | EB354 | Freund et al., 2017 | Kappa | IGKV1-39 | IGKJ2 | 80,4 | QQSHSPPVT | 9 |
| 17 | **BG18** | V3-glycan | EB354 | Freund et al., 2017 | Lambda | IGLV3-25 | IGLJ3 | 82,3 | QSSDTSDSYKM | 11 |
| 18 | **BG8** | V3-glycan | EB354 | Freund et al., 2017 | Lambda | IGLV3-25 | IGLJ3 | 79,5 | QSSDTSDSYKM | 11 |
| 19 | **CH01** | V2-apex | CH0219 | Bonsignori et al., 2011 | Kappa | IGKV3-20 | IGKJ1 | 88,9 | QQYGGSPYT | 9 |
| 20 | **CH103** | CD4bs | Donor CH505 | Liao et al., 2013 | Lambda | IGLV3-1 | IGLJ1 | 85,5 | QVWDSFSTFV | 10 |
| 21 | **CH235** | CD4bs | Donor CH505 | Gao et al., 2014 | Kappa | IGKV3-15 | IGKJ1 | 95 | LQYNNWWT | 8 |
| 22 | **CH235.12** | CD4bs | Donor CH505 | Bonsignori et al., 2016 | Kappa | IGKV3-15 | IGKJ1 | 85,8 | LQYNNWWT | 8 |
| 23 | **CH235.9** | CD4bs | Donor CH505 | Bonsignori et al., 2016 | Kappa* | IGKV3-15* | IGKJ1* | n/a | n/a | n/a |
| 24 | **DH270.1** | V3-glycan | CH848 | Bonsignori et al., 2017 | Lambda | IGLV2-23 | IGLJ2 | 94,9 | CSYAGSSIIF | 10 |
| 25 | **DH270.5** | V3-glycan | CH848 | Bonsignori et al., 2017 | Lambda | IGLV2-23 | IGLJ2 | 89,1 | CSFGGSAAVV | 10 |
| 26 | **DH270.6** | V3-glycan** | CH848** | Bonsignori et al., 2017 | Lambda | IGLV2-23 | IGLJ2 | 91,8 | CSFGGSATVV | 10 |
| 27 | **DH511.11P** | MPER | CH0210 | Williams et al., 2017 | Kappa | IGKV1-39 | IGKJ2 | 83,6 | QESYSSVPMYI | 11 |
| 28 | **DH511.12P** | MPER | CH0210 | Williams et al., 2017 | n/a | n/a | n/a | n/a | n/a | n/a |
| 29 | **DH511.2** | MPER | CH0210 | Williams et al., 2017 | Kappa | IGKV1-39 | IGKJ2 | 85,4 | QENYNTIPSLS | 11 |
| 30 | **HJ16** | CD4bs | VI3208 | Corti et al., 2010 | Kappa | IGKV4-1 | IGKJ4 | 79,7 | QQTRWTPPT | 9 |
| 31 | **IOMA** | CD4bs | R1 | Gristick et al., 2016 | Lambda | IGLV2-23 | IGLJ2 | 92 | YSYADGVAF* | 9 |
| 32 | **N6** | CD4bs | Z258 | Huang et al., 2016 | Kappa | IGKV1-33 | IGKJ5 | 77,8 | QVLQF | 5 |
| 33 | **NIH45-46** | CD4bs | NIH45 | Scheid et al., 2011 | Kappa | IGKV3-11 | IGKJ2 | 81,5 | QQYEF | 5 |
| 34 | **PCDN-33A** | V3-glycan | PC76 | MacLeod et al., 2016 | Kappa | IGKV3-20 | IGKJ1 | 88,9 | QQCGSSPT | 8 |
| 35 | **PCDN-38A** | V3-glycan | PC76 | MacLeod et al., 2016 | Kappa | IGKV3-20 | IGKJ1 | 88,9 | QQYGSSPT | 8 |
| 36 | **PCDN-38B** | V3-glycan | PC76 | MacLeod et al., 2016 | Kappa | IGKV3-20 | IGKJ1 | 88,2 | QQYGSSPT | 8 |
| 37 | **PG16** | V2-apex | Donor 24 | Walker et al., 2009 | Lambda | IGLV2-14 | IGLJ2 | 88,3 | SSLTDRSHRI | 10 |
| 38 | **PG9** | V2-apex | Donor 24 | Walker et al., 2009 | Lambda | IGLV2-14 | IGLJ3 | 92 | KSLTSTRRRV | 10 |
| 39 | **PGDM1400** | V2-apex | Donor 84 | Sok et al., 2014 | Kappa | IGKV2-28 | IGKJ1 | 87,6 | MQGRESPWT | 9 |
| 40 | **PGT121** | V3-glycan | Donor 17 | Walker et al., 2011 | Lambda | IGLV3-21 | IGLJ3 | 81,3 | HIWDSRVPTKWV | 12 |
| 41 | **PGT123** | V3-glycan | Donor 17 | Walker et al., 2011 | Lambda | IGLV3-21 | IGLJ3 | 75,6 | HIYDARGGTNWV | 12 |
| 42 | **PGT125** | V3-glycan | Donor 36 | Walker et al., 2011 | Lambda | IGLV2-8 | IGLJ2 | 80,4 | GSLVGNWDVI | 10 |
| 43 | **PGT126** | V3-glycan | Donor 36 | Walker et al., 2011 | Lambda | IGLV2-8 | IGLJ2 | 86,7 | SSLVGNWDVI | 10 |
| 44 | **PGT127** | V3-glycan | Donor 36 | Walker et al., 2011 | Lambda | IGLV2-8 | IGLJ2 | 86,7 | SSLVGNWDVI | 10 |
| 45 | **PGT128** | V3-glycan | Donor 36 | Walker et al., 2011 | Lambda | IGLV2-8 | IGLJ2 | 86,7 | GSLVGNWDVI | 10 |
| 46 | **PGT130** | V3-glycan | Donor 36 | Walker et al., 2011 | Lambda | IGLV2-8 | IGLJ2 | 88,5 | SSLFGRWDVV | 10 |
| 47 | **PGT135** | V3-glycan | Donor 39 | Walker et al., 2011 | Kappa | IGKV3-15 | IGKJ1 | 82,9 | QQYEEWPRT | 9 |
| 48 | **PGT143** | V2-apex | Donor 84 | Walker et al., 2011 | Kappa | IGKV2-28 | IGKJ1 | 86,9 | MQGLNRPWT | 9 |
| 49 | **PGT145** | V2-apex | Donor 84 | Walker et al., 2011 | Kappa | IGKV2-28 | IGKJ1 | 83,4 | MQGLHSPWT | 9 |
| 50 | **PGT151** | Interface/FP | Donor 31 | Falkowska et al., 2014 | Kappa | IGKV2-29 | IGKJ4 | 88 | MQSKDFPLT* | 9 |
| 51 | **PGT152** | Interface/FP | Donor 31 | Falkowska et al., 2014 | Kappa | IGKV2-29 | IGKJ4 | 88 | MQSKDFPLT | 9 |
| 52 | **VRC-CH31** | CD4bs | CH0219 | Wu et al., 2011 | Kappa | IGKV1-33 | IGKJ2 | 84,7 | QQYET | 5 |
| 53 | **VRC-PG04** | CD4bs | Donor 74 | Wu et al., 2011 | Kappa | IGKV3-20 | IGKJ5 | 78,8 | QQLEF | 5 |
| 54 | **VRC-PG20** | CD4bs | IAVI 23 | Zhou et al., 2013 | Lambda | IGLV2-14 | IGLJ2 | 78,8 | NAYEF* | 5 |
| 55 | **VRC01** | CD4bs | NIH45 | Wu et al., 2010 | Kappa | IGKV3-11 | IGKJ2 | 81,1 | QQYEF | 5 |
| 56 | **VRC03** | CD4bs | NIH45 | Wu et al., 2010 | Kappa | IGKV3-20 | IGKJ2 | 78,7 | QQFEF | 5 |
| 57 | **VRC06** | CD4bs | NIH45 | Li et al., 2012 | Kappa | IGKV3-20 | IGKJ2 | 82,2 | QQFEF | 5 |
| 58 | **VRC06b** | CD4bs | NIH45 | Li et al., 2012 | Kappa | IGKV3-20 | IGKJ2 | 79,8 | QQFEF | 5 |
| 59 | **VRC07** | CD4bs | NIH45 | Rudicelli et al., 2014 | Kappa* | IGKV3-20* | n/a | n/a | n/a | n/a |
| 60 | **VRC13** | CD4bs | 44 | Zhou et al., 2015 | Lambda* | IGLV2-14* | n/a | n/a | n/a | n/a |
| 61 | **VRC16** | CD4bs | C38 | Zhou et al., 2015 | Kappa* | IGKV1-39* | n/a | n/a | n/a | n/a |
| 62 | **VRC18** | CD4bs | C38 | Zhou et al., 2015 | Kappa* | IGKV3-20* | n/a | n/a | n/a | n/a |
| 63 | **VRC26.01** | V2-apex | CAP256 | Doria-Rose et al., 2014 | Lambda | IGLV1-51 | IGLJ1 | 95,9 | GTWDTSLSGGGV | 12 |
| 64 | **VRC26.03** | V2-apex | CAP256 | Doria-Rose et al., 2014 | Lambda | IGLV1-51 | IGLJ1 | 92,8 | ATWSASLSSARV | 12 |
| 65 | **VRC26.06** | V2-apex | CAP256 | Doria-Rose et al., 2014 | Lambda | IGLV1-51 | IGLJ1 | 91,8 | ETWDGSGGV | 9 |
| 66 | **VRC26.08** | V2-apex | CAP256 | Doria-Rose et al., 2014 | Lambda | IGLV1-51 | IGLJ1 | 90,5 | TVWGVRKGVGAV | 12 |
| 67 | **VRC26.09** | V2-apex | CAP256 | Doria-Rose et al., 2014 | Lambda | IGLV1-51 | IGLJ1 | 89,5 | AVWGVRRGAGAV | 12 |
| 68 | **VRC26.25** | V2-apex | CAP256 | Doria-Rose et al., 2016 | Lambda | IGLV1-51 | IGLJ1 | 90,4 | ATWAASLSSARV | 12 |
| 69 | **VRC26.26** | V2-apex | CAP256 | Doria-Rose et al., 2016 | Lambda | IGLV1-51 | IGLJ1 | 89,7 | ATWGVRQGVGAL | 12 |
| 70 | **VRC26.27** | V2-apex | CAP256 | Doria-Rose et al., 2016 | Lambda | IGLV1-51 | IGLJ1 | 90,1 | ATWGVRQGVGAL | 12 |
| 71 | **VRC34.01** | Interface/FP | N123 | Kong et al., 2016 | Kappa | IGKV1-9 | IGKJ4 | 90,2 | QHMSSYPLT | 9 |
| 72 | **VRC38.01** | V2-apex | N90 | Cale et al., 2017 | Kappa | IGKV2-28 | IGKJ4 | 91,6 | MEARQTPRLT | 10 |
| 73 | **b12** | CD4bs | Donor b | Burton et al., 1991 | Kappa | IGKV3-20 | IGKJ2 | 87,7 | QVYGASSYT | 9 |
| 74 | **3BNC55** | CD4bs | Patient 3 | Scheid et al., 2011 | Kappa | IGKV1-33 | IGKJ3 | 84,4 | QVYEF | 5 |
| 75 | **NC37** | CD4bs | EB354 | Freund et al., 2017 | Kappa | IGKV3-20 | IGKJ3 | 83,8 | QNSGGGTPLI | 10 |

Note: V(D)J information, V gene germline identity and CDR3 amino acid sequence were derived by annotation with IgBLAST according to the IMGT nomenclature and numbering.
n/a: not available
*For light chains, where no nucleotide sequence was available or IgBLAST annotation failed, data has been filled up with information deposited in the CATNAP database
** Epitope and Donor Information for DH270.6 was not deposited in the CATNAP database and therefore derived from the reference

**Table S5: Heavy and light chain Pgen, Pshm, and probability scores**

| | Heavy chains | | | | | | | | | Light chains | | | | | | | | |
|---|---|---|---|---|---|---|---|---|---|---|---|---|---|---|---|---|---|---|
| | Healthy cohort (n=57) | | | HIV-1 infected cohort (n=34) | | | HCV infected cohort (n=12) | | | Healthy cohort (n=57) | | | HIV-1 infected cohort (n=34) | | | HCV infected cohort (n=12) | | |
| | log(Pgen) | log(Pshm) | Score S | log(Pgen) | log(Pshm) | Score S | log(Pgen) | log(Pshm) | Score S | log(Pgen) | log(Pshm) | Score S | log(Pgen) | log(Pshm) | Score S | log(Pgen) | log(Pshm) | Score S |
| **10-1074** | -27,848 | -91,758 | -0,401 | -27,896 | -92,085 | -0,402 | -27,247 | -93,076 | -0,403 | -9,802 | -80,913 | -0,377 | -15,761 | -75,610 | -0,398 | -15,964 | -76,198 | -0,401 |
| **10E8** | -25,535 | -119,687 | -0,480 | -25,787 | -120,942 | -0,485 | -39,264 | -118,166 | -0,530 | -6,085 | -78,874 | -0,344 | -5,906 | -79,145 | -0,344 | -5,881 | -79,765 | -0,346 |
| **12A12** | -21,099 | -121,869 | -0,469 | -21,867 | -122,974 | -0,476 | -21,634 | -123,764 | -0,477 | -11,213 | -99,336 | -0,457 | -11,720 | -99,497 | -0,461 | -12,138 | -100,785 | -0,469 |
| **179NC75** | -42,471 | -11,550 | -0,206 | -46,929 | -11,361 | -0,223 | -41,892 | -10,674 | -0,201 | -6,843 | -12,385 | -0,094 | -6,967 | -12,087 | -0,094 | -6,904 | -12,320 | -0,095 |
| **1B2530** | -28,802 | -147,175 | -0,580 | -29,668 | -148,449 | -0,587 | -28,465 | -150,153 | -0,588 | -19,244 | -88,876 | -0,472 | -18,266 | -89,671 | -0,469 | -19,564 | -89,799 | -0,478 |
| **3074** | -19,897 | -47,837 | -0,231 | -19,750 | -48,077 | -0,231 | -20,033 | -47,994 | -0,232 | n/a | n/a | n/a | n/a | n/a | n/a | n/a | n/a | n/a |
| **35O22** | -30,155 | -134,878 | -0,547 | -30,286 | -135,963 | -0,550 | -30,431 | -137,090 | -0,555 | -16,416 | -123,430 | -0,585 | -16,962 | -123,690 | -0,590 | -18,777 | -123,571 | -0,602 |
| **3BNC117** | -14,140 | -145,771 | -0,517 | -14,495 | -146,935 | -0,522 | -14,047 | -148,780 | -0,526 | -8,779 | -92,468 | -0,414 | -8,761 | -92,776 | -0,415 | -8,868 | -94,064 | -0,421 |
| **447-52D** | -16,350 | -48,637 | -0,219 | -16,787 | -48,804 | -0,221 | -23,356 | -44,518 | -0,234 | -4,123 | -19,071 | -0,101 | -4,103 | -19,000 | -0,101 | -3,985 | -19,028 | -0,100 |
| **4E10** | -31,111 | -68,769 | -0,341 | -30,382 | -69,416 | -0,341 | -29,527 | -70,794 | -0,342 | -12,332 | -41,604 | -0,244 | -12,268 | -41,580 | -0,243 | -11,998 | -42,337 | -0,245 |
| **561_01_18** | -17,790 | -148,382 | -0,540 | -17,133 | -150,266 | -0,543 | -18,490 | -151,646 | -0,553 | -4,557 | -122,876 | -0,502 | -4,526 | -122,884 | -0,502 | -4,519 | -124,949 | -0,509 |
| **561_01_55** | -18,238 | -145,237 | -0,532 | -16,957 | -147,462 | -0,534 | -18,574 | -148,475 | -0,543 | -4,557 | -124,813 | -0,509 | -4,526 | -124,717 | -0,509 | -4,519 | -126,871 | -0,517 |
| **561_02_12** | -31,416 | -137,037 | -0,558 | -31,094 | -138,671 | -0,562 | -31,361 | -139,943 | -0,567 | -21,384 | -111,046 | -0,572 | -21,178 | -111,080 | -0,571 | -21,431 | -112,938 | -0,580 |
| **8ANC131** | -20,750 | -148,029 | -0,550 | -20,496 | -149,783 | -0,555 | -20,027 | -151,603 | -0,559 | n/a | n/a | n/a | n/a | n/a | n/a | n/a | n/a | n/a |
| **8ANC195** | -34,478 | -144,879 | -0,595 | -35,996 | -144,936 | -0,602 | -35,579 | -146,987 | -0,606 | -5,606 | -78,960 | -0,341 | -5,612 | -79,095 | -0,341 | -5,601 | -80,119 | -0,345 |
| **BG1** | -32,831 | -149,572 | -0,604 | -33,019 | -151,099 | -0,609 | -30,632 | -154,640 | -0,611 | -5,794 | -108,598 | -0,456 | -6,010 | -108,519 | -0,457 | -6,340 | -109,858 | -0,464 |
| **BG18** | -34,234 | -115,902 | -0,503 | -32,527 | -118,904 | -0,506 | -33,529 | -118,585 | -0,509 | -5,734 | -97,537 | -0,413 | -5,887 | -97,760 | -0,415 | -5,760 | -98,753 | -0,418 |
| **BG8** | -34,570 | -140,245 | -0,581 | -34,234 | -142,406 | -0,587 | -34,697 | -143,010 | -0,590 | -5,762 | -115,529 | -0,482 | -6,145 | -115,522 | -0,485 | -5,914 | -116,876 | -0,488 |
| **CH01** | -37,563 | -92,905 | -0,444 | -37,712 | -92,724 | -0,444 | -36,582 | -95,161 | -0,447 | -4,463 | -65,231 | -0,280 | -4,622 | -65,156 | -0,281 | -4,512 | -66,052 | -0,284 |
| **CH103** | -22,133 | -90,239 | -0,373 | -22,553 | -91,244 | -0,378 | -21,258 | -92,536 | -0,377 | -15,595 | -60,226 | -0,338 | -15,725 | -60,519 | -0,340 | -16,781 | -60,837 | -0,348 |
| **CH235** | -20,179 | -47,628 | -0,231 | -19,995 | -47,865 | -0,231 | -20,362 | -47,617 | -0,232 | -4,197 | -31,837 | -0,151 | -4,338 | -31,937 | -0,152 | -4,239 | -32,026 | -0,152 |
| **CH235.12** | -21,518 | -145,845 | -0,547 | -21,581 | -147,391 | -0,552 | -21,743 | -148,659 | -0,556 | -4,197 | -86,966 | -0,362 | -4,338 | -87,088 | -0,363 | -4,239 | -88,248 | -0,367 |
| **CH235.9** | -19,996 | -116,331 | -0,447 | -20,121 | -117,341 | -0,451 | -20,082 | -118,492 | -0,455 | n/a | n/a | n/a | n/a | n/a | n/a | n/a | n/a | n/a |
| **DH270.1** | -21,844 | -41,412 | -0,218 | -21,904 | -41,046 | -0,217 | -22,104 | -41,218 | -0,218 | -6,474 | -39,824 | -0,197 | -6,652 | -39,718 | -0,198 | -6,872 | -39,651 | -0,199 |
| **DH270.5** | -18,989 | -73,736 | -0,309 | -19,051 | -74,077 | -0,310 | -18,846 | -74,485 | -0,311 | -8,222 | -60,574 | -0,288 | -8,420 | -60,777 | -0,291 | -8,794 | -61,132 | -0,294 |
| **DH270.6** | -20,576 | -81,396 | -0,339 | -20,460 | -81,976 | -0,341 | -21,053 | -81,912 | -0,343 | -4,122 | -55,466 | -0,241 | -4,505 | -55,403 | -0,243 | -4,526 | -55,683 | -0,244 |
| **DH511.11P** | -19,196 | -90,953 | -0,364 | -19,060 | -91,624 | -0,366 | -19,141 | -92,146 | -0,368 | -11,083 | -94,098 | -0,436 | -11,126 | -94,181 | -0,437 | -10,870 | -95,767 | -0,441 |
| **DH511.12P** | -19,197 | -94,380 | -0,375 | -19,059 | -95,090 | -0,377 | -19,145 | -95,652 | -0,379 | n/a | n/a | n/a | n/a | n/a | n/a | n/a | n/a | n/a |
| **DH511.2** | -27,974 | -120,153 | -0,491 | -27,708 | -121,616 | -0,495 | -27,837 | -122,060 | -0,497 | -15,878 | -85,811 | -0,438 | -15,577 | -85,793 | -0,435 | -15,634 | -87,046 | -0,441 |
| **HJ16** | -31,987 | -161,349 | -0,637 | -32,626 | -162,868 | -0,645 | -31,885 | -165,029 | -0,649 | -17,148 | -118,229 | -0,570 | -16,997 | -118,242 | -0,569 | -17,248 | -120,152 | -0,579 |
| **IOMA** | -31,158 | -69,972 | -0,345 | -32,576 | -69,321 | -0,349 | -32,392 | -69,866 | -0,350 | n/a | n/a | n/a | n/a | n/a | n/a | n/a | n/a | n/a |
| **N6** | -21,967 | -165,217 | -0,610 | -21,918 | -167,300 | -0,616 | -23,537 | -167,168 | -0,622 | -14,811 | -127,895 | -0,591 | -14,612 | -128,116 | -0,591 | -14,675 | -130,146 | -0,599 |
| **NIH45-46** | -28,772 | -181,382 | -0,688 | -28,111 | -183,912 | -0,693 | -29,603 | -185,458 | -0,704 | -9,361 | -95,416 | -0,430 | -9,374 | -95,334 | -0,429 | -9,503 | -96,684 | -0,435 |
| **PCDN-33A** | -20,112 | -77,519 | -0,325 | -20,090 | -77,921 | -0,326 | -21,013 | -78,054 | -0,330 | -2,897 | -67,386 | -0,278 | -2,893 | -67,432 | -0,278 | -2,864 | -68,256 | -0,281 |
| **PCDN-38A** | -22,254 | -87,678 | -0,366 | -22,335 | -88,385 | -0,368 | -23,666 | -88,165 | -0,373 | -2,899 | -69,765 | -0,287 | -2,895 | -69,788 | -0,287 | -2,866 | -70,667 | -0,290 |
| **PCDN-38B** | -22,533 | -98,851 | -0,402 | -22,632 | -99,584 | -0,405 | -23,941 | -99,728 | -0,411 | -2,897 | -71,278 | -0,293 | -2,893 | -71,305 | -0,293 | -2,864 | -72,198 | -0,296 |
| **PG16** | -39,636 | -85,276 | -0,428 | -38,750 | -86,726 | -0,429 | -39,010 | -86,432 | -0,429 | -7,680 | -71,690 | -0,327 | -7,887 | -71,655 | -0,329 | -7,810 | -72,504 | -0,331 |
| **PG9** | -33,826 | -79,465 | -0,386 | -33,277 | -79,983 | -0,386 | -33,478 | -80,498 | -0,388 | -9,296 | -54,768 | -0,274 | -9,361 | -54,718 | -0,274 | -10,155 | -54,260 | -0,277 |
| **PGDM1400** | -57,196 | -148,564 | -0,698 | -59,855 | -147,810 | -0,706 | -58,694 | -150,064 | -0,708 | -4,017 | -74,397 | -0,312 | -3,980 | -74,442 | -0,312 | -3,974 | -75,346 | -0,316 |
| **PGT121** | -35,580 | -112,444 | -0,497 | -34,727 | -114,249 | -0,500 | -35,368 | -114,501 | -0,503 | -15,471 | -71,120 | -0,379 | -16,261 | -71,324 | -0,385 | -16,613 | -71,822 | -0,389 |
| **PGT123** | -37,037 | -124,145 | -0,540 | -36,053 | -126,142 | -0,542 | -35,861 | -127,540 | -0,546 | -9,346 | -102,697 | -0,457 | -9,227 | -103,008 | -0,458 | -8,794 | -103,498 | -0,457 |
| **PGT125** | -33,162 | -129,441 | -0,541 | -33,208 | -130,852 | -0,546 | -33,494 | -131,750 | -0,550 | -8,299 | -88,355 | -0,395 | -9,280 | -87,802 | -0,400 | -8,142 | -89,339 | -0,398 |
| **PGT126** | -32,392 | -110,086 | -0,477 | -32,258 | -111,514 | -0,481 | -32,311 | -112,193 | -0,483 | -7,799 | -49,783 | -0,244 | -10,057 | -48,408 | -0,254 | -7,604 | -50,197 | -0,244 |
| **PGT127** | -32,936 | -97,821 | -0,441 | -32,312 | -99,467 | -0,443 | -33,079 | -99,658 | -0,447 | -8,290 | -50,440 | -0,250 | -9,294 | -49,647 | -0,254 | -9,465 | -49,953 | -0,256 |
| **PGT128** | -28,770 | -112,443 | -0,470 | -29,859 | -112,532 | -0,475 | -30,169 | -113,132 | -0,478 | -7,798 | -49,600 | -0,243 | -10,587 | -48,265 | -0,258 | -10,495 | -48,500 | -0,258 |
| **PGT130** | -33,356 | -120,897 | -0,515 | -33,807 | -121,671 | -0,519 | -32,967 | -123,573 | -0,522 | -8,613 | -71,255 | -0,332 | -8,546 | -71,744 | -0,333 | -8,494 | -72,037 | -0,334 |
| **PGT135** | -25,264 | -112,140 | -0,455 | -26,979 | -111,293 | -0,459 | -27,477 | -112,130 | -0,464 | -3,900 | -97,388 | -0,400 | -3,905 | -97,353 | -0,400 | -3,935 | -98,778 | -0,405 |
| **PGT143** | -58,645 | -93,856 | -0,531 | -58,868 | -94,649 | -0,534 | -58,801 | -95,426 | -0,536 | -4,018 | -90,863 | -0,375 | -4,320 | -90,601 | -0,377 | -4,063 | -92,069 | -0,380 |
| **PGT145** | -57,094 | -100,313 | -0,545 | -58,552 | -99,884 | -0,549 | -57,421 | -101,679 | -0,551 | -3,835 | -98,825 | -0,405 | -3,834 | -98,890 | -0,405 | -3,798 | -100,262 | -0,410 |
| **PGT151** | -32,202 | -115,048 | -0,492 | -33,237 | -115,376 | -0,497 | -30,995 | -117,615 | -0,495 | n/a | n/a | n/a | n/a | n/a | n/a | n/a | n/a | n/a |
| **PGT152** | -28,975 | -106,309 | -0,452 | -29,225 | -107,423 | -0,456 | -26,749 | -109,391 | -0,452 | -9,818 | -75,983 | -0,358 | -12,015 | -73,386 | -0,364 | -7,428 | -79,590 | -0,356 |
| **VRC-CH31** | -27,223 | -133,147 | -0,529 | -26,932 | -134,049 | -0,531 | -28,146 | -135,208 | -0,540 | -7,536 | -85,683 | -0,380 | -7,417 | -85,879 | -0,380 | -7,854 | -86,610 | -0,386 |
| **VRC-PG04** | -20,837 | -151,642 | -0,562 | -21,468 | -153,142 | -0,570 | -21,249 | -154,570 | -0,573 | -9,081 | -100,448 | -0,447 | -9,109 | -100,327 | -0,447 | -9,436 | -101,689 | -0,454 |
| **VRC-PG20** | -22,697 | -132,308 | -0,509 | -22,692 | -133,354 | -0,512 | -23,645 | -134,506 | -0,519 | n/a | n/a | n/a | n/a | n/a | n/a | n/a | n/a | n/a |
| **VRC01** | -18,061 | -175,075 | -0,625 | -20,419 | -175,692 | -0,637 | -19,444 | -177,944 | -0,640 | -9,361 | -97,709 | -0,438 | -9,374 | -97,627 | -0,438 | -9,503 | -99,018 | -0,444 |
| **VRC03** | -48,638 | -201,480 | -0,831 | -47,764 | -204,557 | -0,837 | -48,108 | -207,876 | -0,849 | -7,429 | -112,580 | -0,482 | -7,378 | -112,573 | -0,482 | -7,674 | -114,170 | -0,490 |
| **VRC06** | -28,513 | -152,764 | -0,596 | -28,986 | -153,966 | -0,602 | -29,099 | -155,535 | -0,608 | -7,710 | -98,885 | -0,432 | -7,676 | -98,881 | -0,431 | -8,028 | -100,138 | -0,439 |
| **VRC06b** | -44,162 | -204,318 | -0,822 | -45,077 | -205,144 | -0,828 | -46,572 | -206,857 | -0,839 | -9,257 | -104,078 | -0,462 | -9,595 | -104,025 | -0,464 | -9,792 | -105,567 | -0,471 |
| **VRC07** | -30,618 | -172,181 | -0,666 | -32,965 | -172,397 | -0,676 | -32,828 | -175,208 | -0,685 | n/a | n/a | n/a | n/a | n/a | n/a | n/a | n/a | n/a |
| **VRC13** | -38,253 | -176,327 | -0,710 | -39,183 | -177,604 | -0,718 | -39,499 | -178,520 | -0,722 | n/a | n/a | n/a | n/a | n/a | n/a | n/a | n/a | n/a |
| **VRC16** | -29,807 | -105,468 | -0,452 | -31,436 | -104,999 | -0,457 | -30,473 | -106,735 | -0,459 | n/a | n/a | n/a | n/a | n/a | n/a | n/a | n/a | n/a |
| **VRC18** | -21,759 | -140,746 | -0,532 | -21,997 | -142,281 | -0,537 | -21,509 | -143,988 | -0,541 | n/a | n/a | n/a | n/a | n/a | n/a | n/a | n/a | n/a |
| **VRC26.01** | -58,628 | -54,685 | -0,407 | -57,428 | -56,012 | -0,406 | -58,501 | -54,897 | -0,407 | -9,260 | -28,124 | -0,171 | n/a | n/a | n/a | -9,417 | -28,102 | -0,172 |
| **VRC26.03** | -60,548 | -59,823 | -0,431 | -61,038 | -59,716 | -0,432 | -59,724 | -60,684 | -0,430 | -14,727 | -42,401 | -0,264 | n/a | n/a | n/a | -15,470 | -42,036 | -0,267 |
| **VRC26.06** | -64,457 | -65,577 | -0,465 | -64,081 | -66,331 | -0,465 | -63,857 | -66,337 | -0,465 | -8,091 | -46,289 | -0,233 | n/a | n/a | n/a | -9,352 | -45,636 | -0,239 |
| **VRC26.08** | -63,448 | -69,392 | -0,473 | -64,859 | -68,983 | -0,477 | -62,675 | -70,586 | -0,473 | -18,145 | -48,030 | -0,309 | n/a | n/a | n/a | -21,858 | -44,422 | -0,320 |
| **VRC26.09** | -64,366 | -86,235 | -0,529 | -65,097 | -86,221 | -0,532 | -63,350 | -87,593 | -0,530 | -10,929 | -53,901 | -0,281 | n/a | n/a | n/a | -11,001 | -54,562 | -0,284 |
| **VRC26.25** | -66,791 | -72,363 | -0,495 | -67,900 | -72,534 | -0,500 | -67,398 | -72,520 | -0,498 | -11,987 | -56,064 | -0,297 | n/a | n/a | n/a | -13,048 | -55,837 | -0,303 |
| **VRC26.26** | -66,389 | -97,297 | -0,573 | -65,287 | -100,038 | -0,577 | -64,650 | -100,367 | -0,575 | -13,480 | -54,567 | -0,302 | n/a | n/a | n/a | -13,412 | -55,547 | -0,305 |
| **VRC26.27** | -65,804 | -93,181 | -0,557 | -65,533 | -94,315 | -0,560 | -64,295 | -95,456 | -0,558 | -13,486 | -58,206 | -0,315 | n/a | n/a | n/a | -13,280 | -59,404 | -0,319 |
| **VRC34.01** | -19,109 | -84,664 | -0,344 | -18,801 | -85,716 | -0,346 | -20,182 | -84,874 | -0,349 | -3,337 | -54,422 | -0,231 | -3,322 | -54,497 | -0,231 | -3,340 | -55,000 | -0,234 |
| **VRC38.01** | -17,860 | -104,928 | -0,403 | -17,473 | -106,138 | -0,405 | -19,167 | -105,621 | -0,410 | -5,830 | -56,922 | -0,258 | -5,783 | -56,945 | -0,258 | -5,798 | -57,490 | -0,260 |
| **b12** | -23,425 | -74,632 | -0,329 | -26,009 | -72,340 | -0,332 | -27,372 | -71,553 | -0,335 | -5,114 | -74,206 | -0,319 | -5,097 | -74,172 | -0,319 | -5,112 | -75,076 | -0,323 |
| **3BNC55** | -16,042 | -122,093 | -0,450 | -16,468 | -122,997 | -0,454 | -17,069 | -123,293 | -0,458 | -8,765 | -67,481 | -0,319 | -8,698 | -67,719 | -0,319 | -8,796 | -68,575 | -0,323 |
| **NC37** | -22,010 | -147,844 | -0,555 | -22,224 | -149,336 | -0,561 | -22,141 | -150,734 | -0,565 | -17,025 | -92,163 | -0,470 | -16,757 | -92,364 | -0,469 | -17,107 | -93,664 | -0,476 |

***Probabilty scores are derived from a linear regression on the heavy chain sequences with S = c1*log(Pgen)+c2*log(Pshm),**
**and c1 = 0.0040 and c2 = 0.0032 for heavy, c1 = 0.0069 and c2 = 0.0038 for kappa, and c1 = 0.0113 and c2 = 0.0048 for lambda chains**

n/a: not available

**Table S6: HIV-1 cohort demographics and sequencing statistics**

| # | Internal ID | Sex | Age | ART | Cell Number | Raw reads | Assembled and V(D)J annotated reads | | | | Full V(D)J alignment | | | | Median number of reads per UMI | | |
|---|---|---|---|---|---|---|---|---|---|---|---|---|---|---|---|---|---|
| | | | | | | Total | H | K | L | Total | H | K | L | Total | H | K | L |
| **1** | IDC0016 | **m** | 57 | Yes | 100.000 | 1.573.779 | 97.533 | 31.292 | 39.641 | 168.466 | 97.533 | 31.215 | 39.641 | 168.389 | 16,5 | 82 | 22 |
| **2** | IDC0042 | **m** | 27 | Yes | 100.000 | 1.447.715 | 113.598 | 74.111 | 81.064 | 268.773 | 113.598 | 73.936 | 81.064 | 268.598 | 24 | 206 | 37 |
| **3** | IDC0067 | **m** | 66 | Yes | 100.000 | 1.665.259 | 47.282 | 32.578 | 24.508 | 104.368 | 47.282 | 32.454 | 24.508 | 104.244 | 8 | 122 | 20 |
| **4** | IDC0094 | **m** | 68 | Yes | 100.000 | 1.851.080 | 61.522 | 25.563 | 23.422 | 110.507 | 61.522 | 25.491 | 23.422 | 110.435 | 5 | 73 | 16 |
| **5** | IDC0345 | **m** | 48 | Yes | 100.000 | 1.794.943 | 96.984 | 33.913 | 27.631 | 158.528 | 96.984 | 33.848 | 27.631 | 158.463 | 10 | 91,5 | 19 |
| **6** | IDC0465 | **m** | 39 | Yes | 100.000 | 1.735.781 | 71.299 | 22.040 | 28.888 | 122.227 | 71.299 | 21.950 | 28.888 | 122.137 | 11 | 72 | 17,5 |
| **7** | IDC0527 | **m** | 48 | Yes | 100.000 | 1.607.190 | 12.184 | 11.791 | 17.736 | 41.711 | 12.184 | 11.760 | 17.736 | 41.680 | 2 | 66,5 | 16 |
| **8** | IDC0562 | **m** | 39 | Yes | 100.000 | 1.411.971 | 83.739 | 34.552 | 29.257 | 147.548 | 83.739 | 34.480 | 29.257 | 147.476 | 20 | 166,5 | 48 |
| **9** | IDC0577 | **m** | 57 | Yes | 100.000 | 1.437.589 | 81.271 | 25.361 | 17.035 | 123.667 | 81.271 | 25.281 | 17.035 | 123.587 | 13 | 79 | 28 |
| **10** | IDC0764 | **m** | 25 | Yes | 100.000 | 1.877.908 | 160.794 | 105.397 | 94.863 | 361.054 | 160.794 | 105.181 | 94.863 | 360.838 | 14 | 83 | 17 |
| **11** | IDC0788 | **m** | 43 | Yes | 100.000 | 1.495.237 | 84.260 | 64.490 | 24.970 | 173.720 | 84.260 | 64.327 | 24.970 | 173.557 | 25 | 181 | 45 |
| **12** | IDC0809 | **m** | 38 | Yes | 100.000 | 1.647.349 | 42.409 | 29.504 | 20.584 | 92.497 | 42.409 | 29.389 | 20.584 | 92.382 | 8 | 105 | 34,5 |
| **13** | IDC0810 | **m** | 28 | Yes | 100.000 | 1.536.682 | 82.313 | 29.385 | 17.618 | 129.316 | 82.313 | 29.190 | 17.618 | 129.121 | 12 | 74 | 30 |
| **14** | IDC0811 | **n/a** | n/a | Yes | 100.000 | 1.788.544 | 76.851 | 32.087 | 17.134 | 126.072 | 76.851 | 31.979 | 17.134 | 125.964 | 9 | 60 | 16 |
| **15** | IDC0813 | **m** | 57 | Yes | 100.000 | 1.540.879 | 115.529 | 39.750 | 44.903 | 200.182 | 115.529 | 39.684 | 44.903 | 200.116 | 14 | 80 | 22 |
| **16** | IDC0816 | **m** | 44 | Yes | 100.000 | 1.547.132 | 18.880 | 8.139 | 6.152 | 33.171 | 18.880 | 8.112 | 6.152 | 33.144 | 1 | 10 | 5 |
| **17** | IDC0817 | **m** | 50 | Yes | 100.000 | 1.551.317 | 84.615 | 40.040 | 40.354 | 165.009 | 84.615 | 39.896 | 40.354 | 164.865 | 16 | 135,5 | 35 |
| **18** | IDC0828 | **f** | 33 | Yes | 100.000 | 1.689.866 | 156.855 | 96.551 | 88.966 | 342.372 | 156.855 | 96.313 | 88.966 | 342.134 | 33 | 160 | 16 |
| **19** | IDC0831 | **m** | 55 | Yes | 100.000 | 1.326.643 | 126.587 | 86.332 | 63.497 | 276.416 | 126.587 | 86.065 | 63.497 | 276.149 | 18 | 150 | 20 |
| **20** | IDC0832 | **f** | 49 | Yes | 100.000 | 1.556.857 | 135.322 | 95.833 | 75.998 | 307.153 | 135.322 | 95.559 | 75.998 | 306.879 | 20 | 270 | 21 |
| **21** | IDC0836 | **m** | 37 | Yes | 100.000 | 1.413.559 | 104.332 | 60.579 | 63.181 | 228.092 | 104.332 | 60.442 | 63.181 | 227.955 | 19 | 134 | 39 |
| **22** | IDF0033 | **f** | 53 | Yes | 100.000 | 2.020.219 | 73.686 | 36.307 | 31.045 | 141.038 | 73.686 | 36.230 | 31.045 | 140.961 | 6 | 88 | 24,5 |
| **23** | IDC0167 | **m** | 28 | No | 100.000 | 1.846.966 | 142.813 | 131.407 | 85.210 | 359.430 | 142.813 | 130.931 | 85.210 | 358.954 | 24 | 110 | 34 |
| **24** | IDC0359 | **m** | 49 | No | 100.000 | 1.252.176 | 121.479 | 111.309 | 99.026 | 331.814 | 121.479 | 111.094 | 99.026 | 331.599 | 30 | 215,5 | 15 |
| **25** | IDC0456 | **m** | 32 | No | 100.000 | 1.553.823 | 129.205 | 90.033 | 78.051 | 297.289 | 129.205 | 89.738 | 78.051 | 296.994 | 28 | 132 | 41,5 |
| **26** | IDC0466 | **m** | 53 | No | 100.000 | 1.512.996 | 150.032 | 91.768 | 69.730 | 311.530 | 150.032 | 91.561 | 69.730 | 311.323 | 35 | 189 | 19 |
| **27** | IDC0648 | **m** | 41 | No | 100.000 | 1.503.977 | 154.025 | 141.541 | 94.471 | 390.037 | 154.025 | 141.266 | 94.471 | 389.762 | 20,5 | 122 | 19 |
| **28** | IDC0790 | **m** | 34 | No | 100.000 | 1.471.242 | 142.636 | 120.342 | 116.815 | 379.793 | 142.636 | 120.092 | 116.815 | 379.543 | 29 | 129,5 | 5 |
| **29** | IDC0792 | **f** | 41 | No | 100.000 | 1.860.496 | 51.727 | 39.771 | 37.798 | 129.296 | 51.727 | 39.632 | 37.798 | 129.157 | 12 | 67,5 | 33 |
| **30** | IDC0795 | **m** | 33 | No | 100.000 | 1.997.426 | 35.968 | 5.257 | 10.188 | 51.413 | 35.968 | 5.119 | 10.188 | 51.275 | 2 | 49,5 | 3 |
| **31** | IDC0821 | **m** | 56 | No | 100.000 | 1.464.432 | 175.416 | 150.737 | 122.385 | 448.538 | 175.416 | 150.509 | 122.385 | 448.310 | 49 | 225 | 30,5 |
| **32** | IDC0833 | **m** | 50 | No | 100.000 | 1.608.243 | 155.144 | 107.663 | 119.924 | 382.731 | 155.144 | 107.458 | 119.924 | 382.526 | 22,5 | 129 | 44,5 |
| **33** | IDC0837 | **f** | 27 | No | 100.000 | 1.308.500 | 132.230 | 104.661 | 87.054 | 323.945 | 132.230 | 104.530 | 87.054 | 323.814 | 14 | 110 | 31 |
| **34** | IDC0846 | **m** | 39 | No | 100.000 | 1.730.801 | 178.317 | 116.609 | 99.074 | 394.000 | 178.317 | 116.264 | 99.074 | 393.655 | 23 | 132,5 | 29,5 |
| **Minimum** | | | 25 | - | 100.000 | 1.252.176 | 12.184 | 5.257 | 6.152 | 33.171 | 12.184 | 5.119 | 6.152 | 33.144 | 1 | 10 | 3 |
| **Maximum** | | | 68 | - | 100.000 | 2.020.219 | 178.317 | 150.737 | 122.385 | 448.538 | 178.317 | 150.509 | 122.385 | 448.310 | 49 | 270 | 48 |
| **Mean** | | | 44 | - | 100.000 | **1.606.723** | 102.848 | 65.491 | 55.829 | 224.168 | 102.848 | 65.323 | 55.829 | 224.000 | 17 | 121 | 25 |
| **Median** | | | 43 | - | 100.000 | 1.555.340 | 100.933 | 50.310 | 42.629 | 186.951 | 100.933 | 50.169 | 42.629 | 186.837 | 16 | 116 | 22 |
| **Std. deviation** | | | 11 | - | 0 | 188.363 | 44.430 | 41.788 | 35.258 | 117.868 | 44.430 | 41.710 | 35.258 | 117.793 | 10 | 56 | 11 |
| **Sum** | | | 1.445 | - | 3.400.000 | **54.628.577** | 3.496.837 | 2.226.693 | 1.898.173 | 7.621.703 | 3.496.837 | 2.220.976 | 1.898.173 | 7.615.986 | - | - | - |

**Table S6: HIV-1 cohort demographics and sequencing statistics - continued**

| # | Internal ID | Sex | Age | ART | Cell Number | Raw reads | Sequences with ≥3 reads per UMI | | | | Productive sequences | | | | Unique productive CDR3s | | | |
|---|---|---|---|---|---|---|---|---|---|---|---|---|---|---|---|---|---|---|
| | | | | | | Total | H | K | L | Total | H | K | L | Total | H | K | L | Total |
| **1** | IDC0016 | **m** | 57 | Yes | 100.000 | 1.573.779 | 5.447 | 14.948 | 10.940 | 31.335 | 5.190 | 14.276 | 10.247 | 29.713 | 4.298 | 7.012 | 5.414 | 16.724 |
| **2** | IDC0042 | **m** | 27 | Yes | 100.000 | 1.447.715 | 12.619 | 32.142 | 19.695 | 64.456 | 12.151 | 30.605 | 18.054 | 60.810 | 8.743 | 9.187 | 7.033 | 24.963 |
| **3** | IDC0067 | **m** | 66 | Yes | 100.000 | 1.665.259 | 3.077 | 16.482 | 5.869 | 25.428 | 2.790 | 15.524 | 5.330 | 23.644 | 2.073 | 6.741 | 2.680 | 11.494 |
| **4** | IDC0094 | **m** | 68 | Yes | 100.000 | 1.851.080 | 3.623 | 9.417 | 6.000 | 19.040 | 3.362 | 9.032 | 5.503 | 17.897 | 2.001 | 4.293 | 3.170 | 9.464 |
| **5** | IDC0345 | **m** | 48 | Yes | 100.000 | 1.794.943 | 5.329 | 14.459 | 7.220 | 27.008 | 5.032 | 13.978 | 6.779 | 25.789 | 3.384 | 6.725 | 3.764 | 13.873 |
| **6** | IDC0465 | **m** | 39 | Yes | 100.000 | 1.735.781 | 3.550 | 11.108 | 7.033 | 21.691 | 3.369 | 10.644 | 6.504 | 20.517 | 2.759 | 5.361 | 3.628 | 11.748 |
| **7** | IDC0527 | **m** | 48 | Yes | 100.000 | 1.607.190 | 1.443 | 6.208 | 4.011 | 11.662 | 1.317 | 5.937 | 3.650 | 10.904 | 857 | 2.514 | 1.755 | 5.126 |
| **8** | IDC0562 | **m** | 39 | Yes | 100.000 | 1.411.971 | 6.264 | 18.673 | 9.940 | 34.877 | 5.991 | 18.007 | 9.354 | 33.352 | 4.595 | 7.368 | 4.355 | 16.318 |
| **9** | IDC0577 | **m** | 57 | Yes | 100.000 | 1.437.589 | 4.169 | 13.024 | 7.654 | 24.847 | 3.975 | 12.552 | 7.249 | 23.776 | 2.995 | 6.641 | 4.303 | 13.939 |
| **10** | IDC0764 | **m** | 25 | Yes | 100.000 | 1.877.908 | 7.192 | 24.358 | 16.167 | 47.717 | 6.826 | 23.524 | 15.144 | 45.494 | 5.546 | 9.780 | 7.316 | 22.642 |
| **11** | IDC0788 | **m** | 43 | Yes | 100.000 | 1.495.237 | 9.765 | 32.902 | 13.851 | 56.518 | 9.447 | 30.935 | 12.665 | 53.047 | 7.935 | 11.470 | 5.819 | 25.224 |
| **12** | IDC0809 | **m** | 38 | Yes | 100.000 | 1.647.349 | 5.595 | 16.723 | 11.049 | 33.367 | 5.349 | 15.926 | 10.229 | 31.504 | 4.443 | 6.988 | 5.399 | 16.830 |
| **13** | IDC0810 | **m** | 28 | Yes | 100.000 | 1.536.682 | 6.020 | 15.763 | 7.898 | 29.681 | 5.761 | 14.547 | 7.276 | 27.584 | 4.789 | 6.368 | 3.566 | 14.723 |
| **14** | IDC0811 | **n/a** | n/a | Yes | 100.000 | 1.788.544 | 5.436 | 16.408 | 7.846 | 29.690 | 5.212 | 15.196 | 7.154 | 27.562 | 4.415 | 7.444 | 4.045 | 15.904 |
| **15** | IDC0813 | **m** | 57 | Yes | 100.000 | 1.540.879 | 6.129 | 14.646 | 7.229 | 28.004 | 5.842 | 14.139 | 6.778 | 26.759 | 3.441 | 5.880 | 3.186 | 12.507 |
| **16** | IDC0816 | **m** | 44 | Yes | 100.000 | 1.547.132 | 1.137 | 2.865 | 1.403 | 5.405 | 1.023 | 2.691 | 1.242 | 4.956 | 598 | 1.605 | 819 | 3.022 |
| **17** | IDC0817 | **m** | 50 | Yes | 100.000 | 1.551.317 | 7.193 | 24.083 | 15.349 | 46.625 | 6.895 | 23.243 | 14.156 | 44.294 | 5.982 | 9.321 | 7.027 | 22.330 |
| **18** | IDC0828 | **f** | 33 | Yes | 100.000 | 1.689.866 | 10.044 | 22.782 | 16.127 | 48.953 | 9.644 | 21.752 | 14.949 | 46.345 | 7.668 | 8.507 | 6.618 | 22.793 |
| **19** | IDC0831 | **m** | 55 | Yes | 100.000 | 1.326.643 | 10.254 | 27.390 | 13.018 | 50.662 | 9.967 | 25.555 | 12.066 | 47.588 | 7.919 | 10.147 | 5.422 | 23.488 |
| **20** | IDC0832 | **f** | 49 | Yes | 100.000 | 1.556.857 | 11.359 | 25.296 | 16.115 | 52.770 | 10.965 | 23.363 | 14.848 | 49.176 | 8.368 | 9.319 | 6.179 | 23.866 |
| **21** | IDC0836 | **m** | 37 | Yes | 100.000 | 1.413.559 | 5.196 | 15.861 | 10.371 | 31.428 | 4.905 | 15.186 | 9.509 | 29.600 | 3.473 | 6.247 | 4.430 | 14.150 |
| **22** | IDF0033 | **f** | 53 | Yes | 100.000 | 2.020.219 | 2.993 | 15.217 | 9.182 | 27.392 | 2.746 | 14.546 | 8.456 | 25.748 | 2.074 | 6.110 | 3.910 | 12.094 |
| **23** | IDC0167 | **m** | 28 | No | 100.000 | 1.846.966 | 11.486 | 31.412 | 17.161 | 60.059 | 11.077 | 29.491 | 15.940 | 56.508 | 7.267 | 7.742 | 4.953 | 19.962 |
| **24** | IDC0359 | **m** | 49 | No | 100.000 | 1.252.176 | 13.273 | 31.717 | 20.821 | 65.811 | 12.888 | 30.430 | 19.496 | 62.814 | 8.371 | 8.251 | 6.245 | 22.867 |
| **25** | IDC0456 | **m** | 32 | No | 100.000 | 1.553.823 | 11.797 | 26.313 | 17.209 | 55.319 | 11.432 | 25.239 | 15.632 | 52.303 | 7.314 | 8.202 | 5.966 | 21.482 |
| **26** | IDC0466 | **m** | 53 | No | 100.000 | 1.512.996 | 12.398 | 35.088 | 15.068 | 62.554 | 12.053 | 33.556 | 14.033 | 59.642 | 8.448 | 9.135 | 4.726 | 22.309 |
| **27** | IDC0648 | **m** | 41 | No | 100.000 | 1.503.977 | 13.619 | 33.459 | 18.357 | 65.435 | 13.213 | 32.304 | 17.209 | 62.726 | 8.634 | 9.184 | 5.408 | 23.226 |
| **28** | IDC0790 | **m** | 34 | No | 100.000 | 1.471.242 | 15.256 | 27.847 | 21.104 | 64.207 | 14.883 | 26.548 | 19.522 | 60.953 | 8.621 | 6.177 | 5.457 | 20.255 |
| **29** | IDC0792 | **f** | 41 | No | 100.000 | 1.860.496 | 6.479 | 24.381 | 16.763 | 47.623 | 6.225 | 23.573 | 15.851 | 45.649 | 3.916 | 6.030 | 4.529 | 14.475 |
| **30** | IDC0795 | **m** | 33 | No | 100.000 | 1.997.426 | 1.972 | 3.139 | 1.732 | 6.843 | 1.821 | 3.007 | 1.591 | 6.419 | 900 | 1.684 | 829 | 3.413 |
| **31** | IDC0821 | **m** | 56 | No | 100.000 | 1.464.432 | 17.723 | 32.194 | 24.025 | 73.942 | 17.263 | 30.854 | 22.177 | 70.294 | 10.525 | 7.450 | 5.971 | 23.946 |
| **32** | IDC0833 | **m** | 50 | No | 100.000 | 1.608.243 | 13.550 | 31.397 | 22.000 | 66.947 | 13.197 | 30.242 | 20.948 | 64.387 | 8.718 | 8.054 | 7.337 | 24.109 |
| **33** | IDC0837 | **f** | 27 | No | 100.000 | 1.308.500 | 16.701 | 35.841 | 21.052 | 73.594 | 16.299 | 34.343 | 19.620 | 70.262 | 3.383 | 6.086 | 3.669 | 13.138 |
| **34** | IDC0846 | **m** | 39 | No | 100.000 | 1.730.801 | 13.092 | 35.047 | 19.907 | 68.046 | 12.685 | 33.251 | 18.349 | 64.285 | 8.098 | 6.515 | 4.112 | 18.725 |
| **Minimum** | | | 25 | - | 100.000 | 1.252.176 | 1.137 | 2.865 | 1.403 | 5.405 | 1.023 | 2.691 | 1.242 | 4.956 | 598 | 1.605 | 819 | 3.022 |
| **Maximum** | | | 68 | - | 100.000 | 2.020.219 | 17.723 | 35.841 | 24.025 | 73.942 | 17.263 | 34.343 | 22.177 | 70.294 | 10.525 | 11.470 | 7.337 | 25.224 |
| **Mean** | | | 44 | - | 100.000 | **1.606.723** | 8.270 | 21.723 | 12.917 | 42.910 | 7.965 | 20.706 | 11.986 | 40.656 | 5.369 | 7.045 | 4.678 | 17.092 |
| **Median** | | | 43 | - | 100.000 | 1.555.340 | 6.836 | 23.433 | 13.435 | 47.124 | 6.526 | 22.498 | 12.366 | 44.894 | 4.692 | 7.000 | 4.628 | 16.777 |
| **Std. deviation** | | | 11 | - | 0 | 188.363 | 4.553 | 9.518 | 6.109 | 19.673 | 4.473 | 9.081 | 5.697 | 18.768 | 2.772 | 2.192 | 1.659 | 6.164 |
| **Sum** | | | 1.445 | - | 3.400.000 | **54.628.577** | 281.180 | 738.590 | 439.166 | 1.458.936 | 270.795 | 703.996 | 407.510 | 1.382.301 | 182.551 | 239.538 | 159.040 | 581.129 |

**Table S7: HCV cohort demographics and sequencing statistics**

| # | Internal ID | Sex | Age | Cell Number | Raw reads | Assembled and V(D)J annotated reads | | | | Full V(D)J alignment | | | | Median number of reads per UMI | | |
|---|---|---|---|---|---|---|---|---|---|---|---|---|---|---|---|---|
| | | | | | Total | H | K | L | Total | H | K | L | Total | H | K | L |
| **1** | IDC0814 | **m** | 58 | 100.000 | 1.802.643 | 10.579 | 28.938 | 5.653 | 45.170 | 10.579 | 28.739 | 5.653 | 44.971 | 2 | 28,5 | 9 |
| **2** | IDC0815 | **m** | 30 | 100.000 | 1.585.244 | 8.967 | 15.315 | 19.389 | 43.671 | 8.967 | 15.260 | 19.389 | 43.616 | 4 | 34 | 10 |
| **3** | IDC0820 | **m** | 30 | 100.000 | 1.595.574 | 109.742 | 54.707 | 74.597 | 239.046 | 109.742 | 54.528 | 74.597 | 238.867 | 16 | 64 | 29 |
| **4** | IDC0822 | **m** | 60 | 100.000 | 1.574.480 | 101.119 | 52.161 | 43.934 | 197.214 | 101.119 | 52.014 | 43.934 | 197.067 | 11 | 95 | 39 |
| **5** | IDC0823 | **m** | 57 | 100.000 | 1.484.911 | 48.654 | 31.049 | 16.233 | 95.936 | 48.654 | 30.959 | 16.233 | 95.846 | 12 | 72 | 20 |
| **6** | IDC0826 | **m** | 58 | 100.000 | 1.752.793 | 14.654 | 61.315 | 27.947 | 103.916 | 14.654 | 61.228 | 27.947 | 103.829 | 14 | 172 | 27 |
| **7** | IDC0830 | **m** | 31 | 100.000 | 2.422.696 | 179.070 | 137.978 | 101.170 | 418.218 | 179.070 | 137.547 | 101.170 | 417.787 | 26 | 272 | 40 |
| **8** | IDC0834 | **m** | 61 | 100.000 | 1.562.511 | 126.892 | 84.218 | 63.159 | 274.269 | 126.892 | 84.082 | 63.159 | 274.133 | 10 | 120 | 18 |
| **9** | IDC0840 | **m** | 39 | 100.000 | 1.421.762 | 101.096 | 84.173 | 55.363 | 240.632 | 101.096 | 83.956 | 55.363 | 240.415 | 15 | 158 | 25 |
| **10** | IDC0841 | **m** | 45 | 100.000 | 1.511.192 | 127.965 | 90.892 | 65.970 | 284.827 | 127.965 | 90.717 | 65.970 | 284.652 | 16 | 255 | 34 |
| **11** | IDC0842 | **m** | 32 | 100.000 | 1.418.444 | 143.038 | 83.699 | 91.495 | 318.232 | 143.038 | 83.509 | 91.495 | 318.042 | 19 | 98 | 21 |
| **12** | IDC0843 | **m** | 42 | 100.000 | 1.635.455 | 110.544 | 68.076 | 78.433 | 257.053 | 110.544 | 67.933 | 78.433 | 256.910 | 11 | 177 | 37 |
| **Minimum** | | | 30 | 100.000 | 1.418.444 | 8.967 | 15.315 | 5.653 | 43.671 | 8.967 | 15.260 | 5.653 | 43.616 | 2 | 29 | 9 |
| **Maximum** | | | 61 | 100.000 | 2.422.696 | 179.070 | 137.978 | 101.170 | 418.218 | 179.070 | 137.547 | 101.170 | 417.787 | 26 | 272 | 40 |
| **Mean** | | | 45 | 100.000 | **1.647.309** | 90.193 | 66.043 | 53.612 | 209.849 | 90.193 | 65.873 | 53.612 | 209.678 | 13 | 129 | 26 |
| **Median** | | | 44 | 100.000 | 1.579.862 | 105.431 | 64.696 | 59.261 | 239.839 | 105.431 | 64.581 | 59.261 | 239.641 | 13 | 109 | 26 |
| **Std. deviation** | | | 12 | 0 | 258.811 | 53.904 | 31.996 | 29.750 | 111.077 | 53.904 | 31.925 | 29.750 | 111.011 | 6 | 76 | 10 |
| **Sum** | | | - | 1.200.000 | **19.767.705** | 1.082.320 | 792.521 | 643.343 | 2.518.184 | 1.082.320 | 790.472 | 643.343 | 2.516.135 | - | - | - |

**Table S7: HCV cohort demographics and sequencing statistics - continued**

| # | Internal ID | Sex | Age | Cell Number | Raw reads | Sequences with ≥3 reads per UMI | | | | Productive sequences | | | | Unique productive CDR3s | | | |
|---|---|---|---|---|---|---|---|---|---|---|---|---|---|---|---|---|---|
| | | | | | Total | H | K | L | Total | H | K | L | Total | H | K | L | Total |
| **1** | IDC0814 | **m** | 58 | 100.000 | 1.802.643 | 1.450 | 6.272 | 2.490 | 10.212 | 1.287 | 6.028 | 2.314 | 9.629 | 1.093 | 3.187 | 1.618 | 5.898 |
| **2** | IDC0815 | **m** | 30 | 100.000 | 1.585.244 | 1.876 | 6.256 | 6.594 | 14.726 | 1.765 | 5.936 | 6.011 | 13.712 | 1.620 | 3.609 | 3.730 | 8.959 |
| **3** | IDC0820 | **m** | 30 | 100.000 | 1.595.574 | 6.015 | 15.751 | 14.171 | 35.937 | 5.699 | 14.971 | 13.153 | 33.823 | 4.657 | 5.612 | 5.603 | 15.872 |
| **4** | IDC0822 | **m** | 60 | 100.000 | 1.574.480 | 8.125 | 23.890 | 14.919 | 46.934 | 7.852 | 23.074 | 13.769 | 44.695 | 3.713 | 6.869 | 4.582 | 15.164 |
| **5** | IDC0823 | **m** | 57 | 100.000 | 1.484.911 | 4.293 | 15.924 | 9.901 | 30.118 | 4.033 | 15.260 | 9.027 | 28.320 | 3.353 | 6.425 | 4.586 | 14.364 |
| **6** | IDC0826 | **m** | 58 | 100.000 | 1.752.793 | 6.116 | 29.319 | 11.796 | 47.231 | 5.857 | 28.363 | 10.936 | 45.156 | 4.976 | 9.567 | 4.754 | 19.297 |
| **7** | IDC0830 | **m** | 31 | 100.000 | 2.422.696 | 15.200 | 41.733 | 24.968 | 81.901 | 14.673 | 39.800 | 23.316 | 77.789 | 9.286 | 9.800 | 6.570 | 25.656 |
| **8** | IDC0834 | **m** | 61 | 100.000 | 1.562.511 | 7.793 | 29.962 | 11.618 | 49.373 | 7.418 | 27.927 | 10.585 | 45.930 | 5.509 | 9.474 | 4.677 | 19.660 |
| **9** | IDC0840 | **m** | 39 | 100.000 | 1.421.762 | 7.451 | 25.036 | 10.235 | 42.722 | 7.141 | 24.287 | 9.456 | 40.884 | 4.534 | 6.740 | 4.472 | 15.746 |
| **10** | IDC0841 | **m** | 45 | 100.000 | 1.511.192 | 8.242 | 27.343 | 13.099 | 48.684 | 7.948 | 26.544 | 12.242 | 46.734 | 6.126 | 10.004 | 5.957 | 22.087 |
| **11** | IDC0842 | **m** | 32 | 100.000 | 1.418.444 | 10.175 | 28.220 | 18.977 | 57.372 | 9.714 | 27.004 | 17.605 | 54.323 | 7.995 | 10.222 | 7.992 | 26.209 |
| **12** | IDC0843 | **m** | 42 | 100.000 | 1.635.455 | 7.417 | 24.555 | 16.119 | 48.091 | 7.130 | 23.601 | 14.762 | 45.493 | 5.637 | 9.109 | 6.875 | 21.621 |
| **Minimum** | | | 30 | 100.000 | 1.418.444 | 1.450 | 6.256 | 2.490 | 10.212 | 1.287 | 5.936 | 2.314 | 9.629 | 1.093 | 3.187 | 1.618 | 5.898 |
| **Maximum** | | | 61 | 100.000 | 2.422.696 | 15.200 | 41.733 | 24.968 | 81.901 | 14.673 | 39.800 | 23.316 | 77.789 | 9.286 | 10.222 | 7.992 | 26.209 |
| **Mean** | | | 45 | 100.000 | **1.647.309** | 7.013 | 22.855 | 12.907 | 42.775 | 6.710 | 21.900 | 11.931 | 40.541 | 4.875 | 7.552 | 5.118 | 17.544 |
| **Median** | | | 44 | 100.000 | 1.579.862 | 7.434 | 24.796 | 12.448 | 47.083 | 7.136 | 23.944 | 11.589 | 44.926 | 4.817 | 7.989 | 4.716 | 17.585 |
| **Std. deviation** | | | 12 | 0 | 258.811 | 3.503 | 9.812 | 5.521 | 18.124 | 3.402 | 9.376 | 5.161 | 17.267 | 2.245 | 2.400 | 1.578 | 5.894 |
| **Sum** | | | - | 1.200.000 | **19.767.705** | 84.153 | 274.261 | 154.887 | 513.301 | 80.517 | 262.795 | 143.176 | 486.488 | 58.499 | 90.618 | 61.416 | 210.533 |

**Table S8: HIV-1 cohort poly IgG neutralization on the 12-strain global panel**

| # | Internal ID | ART | IC50 values against global panel (µg/ml) | | | | | | | | | | | | Summary | | |
|---|---|---|---|---|---|---|---|---|---|---|---|---|---|---|---|---|---|
| | | | 398-F1_F6_20 | CNE8 | CNE55 | 246-F3_C10_2 | X2278_C2_B6 | TRO.11 | CH119.10 | BJOX002000.03.2 | 25710-2.43 | Ce1176_A3 | Ce703010217_B6 | X1632_S2_B10 | Mean (µg/ml)* | Breadth (%) | Breadth group |
| 1 | IDF0033 | Yes | 16 | 51 | 65 | 46 | 53 | 27 | 66 | 225 | 27 | 80 | 114 | 85 | 56,6 | 100,0 | High |
| 2 | IDC0527 | Yes | 253 | 240 | 813 | 289 | 79 | 277 | 129 | 282 | 166 | 287 | 449 | 291 | 253,5 | 100,0 | High |
| 3 | IDC0833 | No | >1000 | 23 | 231 | 34 | 71 | 94 | 56 | 244 | 47 | 128 | 55 | >1000 | 74,8 | 83,3 | High |
| 4 | IDC0359 | No | >1000 | 243 | >1000 | >1000 | 181 | 149 | 114 | 680 | 69 | 373 | 207 | 445 | 219,8 | 75,0 | High |
| 5 | IDC0042 | Yes | 24 | 164 | 838 | 256 | 53 | 866 | >1000 | 825 | 866 | >1000 | >1000 | 620 | 295,7 | 75,0 | High |
| 6 | IDC0094 | Yes | 101 | 490 | >1000 | 989 | 173 | 292 | 151 | 401 | 308 | 381 | >1000 | >1000 | 296,3 | 75,0 | High |
| 7 | IDC0562 | Yes | 73 | 48 | 584 | 616 | 989 | >1000 | >1000 | 717 | 64 | >1000 | >1000 | 588 | 276,3 | 66,7 | High |
| 8 | IDC0016 | Yes | 198 | 228 | >1000 | >1000 | 68 | 543 | 336 | >1000 | 201 | 801 | >1000 | 773 | 302,3 | 66,7 | High |
| 9 | IDC0577 | Yes | 38 | 42 | 601 | 876 | >1000 | 492 | >1000 | 944 | 186 | >1000 | >1000 | >1000 | 255,6 | 58,3 | Intermediate |
| 10 | IDC0465 | Yes | 198 | 337 | >1000 | >1000 | 480 | 599 | >1000 | >1000 | 653 | >1000 | 692 | >1000 | 453,2 | 50,0 | Intermediate |
| 11 | IDC0817 | Yes | >1000 | 386 | >1000 | >1000 | 135 | 539 | 717 | >1000 | 357 | >1000 | >1000 | >1000 | 372,5 | 41,7 | Intermediate |
| 12 | IDC0832 | Yes | 44 | >1000 | >1000 | >1000 | >1000 | >1000 | 323 | 894 | >1000 | 681 | >1000 | >1000 | 304,5 | 33,3 | Intermediate |
| 13 | IDC0466 | No | >1000 | 165 | >1000 | >1000 | >1000 | 376 | >1000 | 609 | 313 | >1000 | >1000 | >1000 | 330,2 | 33,3 | Intermediate |
| 14 | IDC0456 | No | >1000 | >1000 | >1000 | 90 | 21 | >1000 | >1000 | 778 | >1000 | >1000 | >1000 | >1000 | 113,0 | 25,0 | Low |
| 15 | IDC0345 | Yes | 249 | 383 | >1000 | >1000 | >1000 | >1000 | >1000 | >1000 | 324 | >1000 | >1000 | >1000 | 313,6 | 25,0 | Low |
| 16 | IDC0831 | Yes | 189 | >1000 | >1000 | >1000 | 487 | >1000 | >1000 | 624 | >1000 | >1000 | >1000 | >1000 | 385,4 | 25,0 | Low |
| 17 | IDC0790 | No | >1000 | >1000 | >1000 | >1000 | >1000 | 699 | 921 | >1000 | 179 | >1000 | >1000 | >1000 | 486,8 | 25,0 | Low |
| 18 | IDC0167 | No | >1000 | 286 | >1000 | >1000 | >1000 | 978 | >1000 | >1000 | 978 | >1000 | >1000 | >1000 | 649,2 | 25,0 | Low |
| 19 | IDC0067 | Yes | >1000 | 125 | >1000 | >1000 | >1000 | >1000 | >1000 | >1000 | 211 | >1000 | >1000 | >1000 | 162,4 | 16,7 | Low |
| 20 | IDC0792 | No | >1000 | 317 | >1000 | >1000 | >1000 | >1000 | >1000 | >1000 | 384 | >1000 | >1000 | >1000 | 349,1 | 16,7 | Low |
| 21 | IDC0816 | Yes | >1000 | 509 | >1000 | >1000 | >1000 | >1000 | >1000 | >1000 | 703 | >1000 | >1000 | >1000 | 598,1 | 16,7 | Low |
| 22 | IDC0809 | Yes | >1000 | 232 | >1000 | >1000 | >1000 | >1000 | >1000 | >1000 | >1000 | >1000 | >1000 | >1000 | 231,7 | 8,3 | Low |
| 23 | IDC0813 | Yes | >1000 | 558 | >1000 | >1000 | >1000 | >1000 | >1000 | >1000 | >1000 | >1000 | >1000 | >1000 | 558,4 | 8,3 | Low |
| 24 | IDC0846 | No | >1000 | 672 | >1000 | >1000 | >1000 | >1000 | >1000 | >1000 | >1000 | >1000 | >1000 | >1000 | 672,0 | 8,3 | Low |
| 25 | IDC0810 | Yes | >1000 | >1000 | >1000 | 714 | >1000 | >1000 | >1000 | >1000 | >1000 | >1000 | >1000 | >1000 | 714,0 | 8,3 | Low |
| 26 | IDC0836 | Yes | 756 | >1000 | >1000 | >1000 | >1000 | >1000 | >1000 | >1000 | >1000 | >1000 | >1000 | >1000 | 756,0 | 8,3 | Low |
| 27 | IDC0821 | No | >1000 | >1000 | >1000 | >1000 | >1000 | >1000 | >1000 | >1000 | 976 | >1000 | >1000 | >1000 | 975,7 | 8,3 | Low |
| 28 | IDC0837 | No | >1000 | >1000 | >1000 | >1000 | >1000 | >1000 | 978 | >1000 | >1000 | >1000 | >1000 | >1000 | 978,3 | 8,3 | Low |
| 29 | IDC0764 | Yes | >1000 | 985 | >1000 | >1000 | >1000 | >1000 | >1000 | >1000 | >1000 | >1000 | >1000 | >1000 | 984,9 | 8,3 | Low |
| 30 | IDC0788 | Yes | >1000 | >1000 | >1000 | >1000 | >1000 | >1000 | >1000 | >1000 | >1000 | >1000 | >1000 | >1000 | >1000 | 0,0 | No neutralization |
| 31 | IDC0811 | Yes | >1000 | >1000 | >1000 | >1000 | >1000 | >1000 | >1000 | >1000 | >1000 | >1000 | >1000 | >1000 | >1000 | 0,0 | No neutralization |
| 32 | IDC0828 | Yes | >1000 | >1000 | >1000 | >1000 | >1000 | >1000 | >1000 | >1000 | >1000 | >1000 | >1000 | >1000 | >1000 | 0,0 | No neutralization |
| 33 | IDC0648 | No | >1000 | >1000 | >1000 | >1000 | >1000 | >1000 | >1000 | >1000 | >1000 | >1000 | >1000 | >1000 | >1000 | 0,0 | No neutralization |
| 34 | IDC0795 | No | n/a | n/a | n/a | n/a | n/a | n/a | n/a | n/a | n/a | n/a | n/a | n/a | n/a | n/a | n/a |

* Geometric mean IC50 of all neutralized strains
n/a: not available
